# Automated design of stiffness-tunable DNA origami hollowframes for self-assembling metamaterials

**DOI:** 10.64898/2026.07.23.740378

**Authors:** Anthony J. Vetturini, Jonathan Cagan, Rebecca E. Taylor

## Abstract

DNA origami offers a route to engineering architected metamaterials with sub-nanometer precision by linking nanoscale building blocks into micron-scale assemblies. However, automated design spaces are currently limited to fixed DNA origami motifs, restricting control over the stiffness of mass-efficient nanostructures. Here, we introduce a fully automated design paradigm that converts prescribed vertices, edges, and cross-section specifications directly into manufacturable, nucleotide-level models. To demonstrate robustness, three structurally distinct nanostructures are realized under a shared experimental protocol. Further, this paradigm enables the programmed assembly of hollowframe building blocks into micron-scale architectures, including traditional and reentrant honeycomb lattices. More broadly, this work establishes a design abstraction for stiffness-tunable DNA origami nanostructures that can be rapidly translated into architected metamaterials with distinct simulation-predicted mechanical responses.

## Introduction

DNA origami (DO) (*1*) enables the manufacturing of nanoscale building blocks that span tens to hundreds of nanometers in size while containing sub-nanometer structural precision (*2, 3*). Collectively, individual DO units can be assembled through complementary sticky ends (*4–6*) or blunt end stacking (*7*) into higher-order lattices with unit cell periodicities ranging from around fifty to several hundred nanometers (*8*). These capabilities have established DO as a powerful platform for architecting periodic lattices such as photonic crystals (*4*), plasmonic metasurfaces (*9*), and mechanical metamaterials (*10, 11*). However, function in such metamaterials comes not only from the existence and periodicity of the assembled lattice, but from the geometry and mechanics of the building blocks themselves. Structural features such as topology, junction architecture, and stiffness collectively influence how a material deforms in response to external stimuli (*10, 12–14*). DO enables these mechanical properties to be tuned by arranging DNA helices into prescribed bundle cross-sections, thereby controlling a nanostructure’s persistence length and bending stiffness (*15, 16*). The cross-section is therefore a central variable for stiffness-tunable DO metamaterials.

Yet, systematically customizing these helix bundle cross-sections across complex unit cell topologies remains constrained by existing computer-aided design (CAD) platforms (*17*– *27*) that typically enforce a tradeoff between design automation and structural customization. Fully automated methods (*19, 20, 24*) convert standard polyhedral meshes directly into nucleotide-level models, bypassing the intricate DO scaffold routing and staple design steps (*3, 28*). However, these methods offer few and fixed helix bundle cross-sections based on the particular algorithm, limiting control over nanostructure stiffness and, consequently, any assembled material response. Conversely, more flexible CAD platforms (*22, 25–27*) offer some automated components alongside a broader and highly tunable design space but require iterative design-simulate-redesign cycles and more substantial designer expertise in the complex DO design principles (*3, 28*). This tradeoff is particularly limiting for lattices, where target unit cells may consist of open or concave geometries, such as reentrant honeycombs, which cannot be represented through closed polyhedral meshes without generating non-uniform edge thicknesses at shared interfaces during colloidal assembly. Thus, advancing DNA origami metamaterials requires a fully automated design framework that moves beyond simple mesh-based inputs by coupling geometry with the arrangement of DNA helices along the edges as tunable parameters rather than fixed consequences of a particular automation algorithm.

Here, we introduce a hollow wireframe, or “hollowframe,” design paradigm that resolves the tradeoff between automation and customization by automatically transforming user-defined wireframe models into nucleotide-level structures with tunable edge cross-sections. Designers specify 3D spatial vertices and connecting edges that dictate the input geometry alongside a target cross-section parameter that influences nanostructure stiffness. From these inputs, a manufacturable nucleotide-level model is automatically defined and structurally optimized through a mitering procedure to ensure global structural compatibility for colloidal assembly. Robust formation of topologically distinct DO nanostructures is demonstrated using a shared thermal annealing protocol with minimal aggregation. Furthermore, this work demonstrates micron-scale traditional and reentrant honeycomb lattices assembled from hollowframe DO building blocks. Coarse-grained molecular dynamics (*29*) simulations are used to directly connect architectural choices to emergent material response. Together, these results establish a versatile topology and stiffness design abstraction, opening access to a broader structure-property space of programmable nanoscale metamaterials.

### Graph-to-nucleotide design abstraction

To start the automated hollowframe process, a designer specifies a wireframe model represented as a graph of vertices and edges alongside an edge cross-section parameter. The cross-section parameter is defined by either a target cross-sectional diameter or as a specified number of DNA helices per edge (N helix bundle (HB) edges) (Fig. 1 A, figs. S1-3). Either specification results in a number of DNA helices arranged in a honeycomb ring-like cross-section (Fig 1 B, i) to approximately maximize the second moment of area, thus increasing the nanostructure edge bending rigidity (*15, 16*). This work’s scaffold routing algorithm (supplementary text) guarantees a solution for any even N ≥ 2 HB specification or uses a heuristic that can target odd N HB edges depending on the input geometry (fig. S2), while also allowing different numbers of helices per edge to be specified. The cross-section (Fig. 1 B i) is then combined with the input graph (Fig. 1 B ii) such that each edge contains a DNA helix bundle whose nucleotide-level positions are individually represented by their center of gravity (X, Y, Z) positions.

**Fig. 1.**
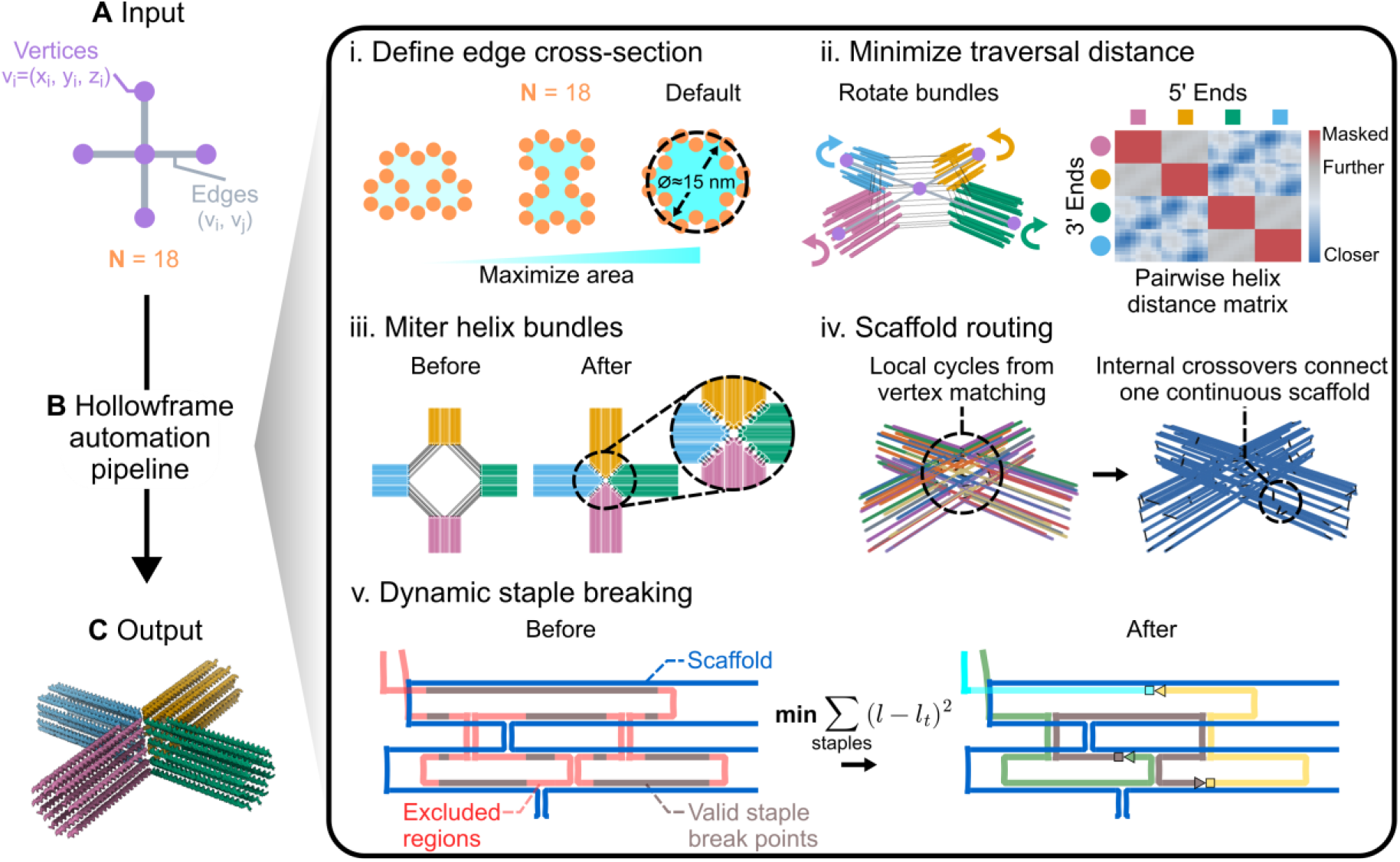
Design automation pipeline for hollowframe DNA origami. (**A**) Inputs are specified as wireframe models composed of vertices and edges alongside an edge cross-section parameter. (**B**) The automation pipeline (i) first converts the cross-section parameter to a hollow honeycomb ring of approximately maximal enclosed area to enhance bending stiffness. (ii) The honeycomb ring is placed at the midpoint of the wireframe model edges. The rings are then rotated about their respective central edge axis to achieve a state where the scaffold traverses a minimal Euclidean distance between DNA helix pairs spanning separate helix bundles. (iii) Nucleotides are populated until a threshold distance is reached between the helix pairs. (iv) Internal crossovers are used to connect a single, continuous scaffold routing. (v) Staples are broken down into manufacturable, low variance lengths by selecting break points that are not near excluded, unfavorable regions (e.g., near crossovers). (**C)** Standard DNA origami design files are then output and provided alongside the staple sequences (*30*).

Next, connections between helices belonging to separate bundles must be defined to prevent a physically infeasible scaffold routing. Individually, bundles are rotated about their central edge axes (Fig. 1 B, ii) and matched at shared vertices by minimizing the pairwise distances between the 3’ and 5’ ends between DNA helices (Fig. 1 B, ii, supplementary text, and fig. S4). Once all connections between bundles are known, the individual helices can be structurally refined. Previous work (*9*) has demonstrated that flexible helix bundle connections leads to off-target or amorphous aggregate assemblies because the DO building blocks lack uniform angles between bundles. While curved helix bundles have been used to circumvent this (*4, 9*), they are topology-dependent and require iterative simulation cycles to achieve a desired curvature. Instead, a geometric mitering step (Fig. 1 B, iii, supplementary text and fig. S5) adjusts connected helix end points by adding or removing nucleotides such that the thick helix bundle edges fit flush at vertices, forming stiff junctions that are conducive to assembly. Single-stranded DNA spacer portions of the scaffold are added to relieve backbone torsion similar to previous methods (*31*), however, oxDNA (*29*) simulations inform the number of nucleotides to use as single-stranded regions (supplementary text and fig. S6 and S7). After defining the scaffold connections between helices at the shared vertices, internal crossovers can be algorithmically applied using a minimum spanning tree algorithm (*22*) (supplementary text, fig. S8) to construct a single, continuous scaffold routing (Fig. 1 B, iv).

Finally, staple oligonucleotides are designed to match the scaffold and produce a manufacturable set of sequences. Recent literature (*32–34*) is used to inform a dynamic stapling procedure (supplementary text and Fig. S9-S11). This involves breaking staples into manufacturable lengths (defaulting to between 20 and 60 nucleotides) in a way that minimizes staple length deviation from a target of 42 nucleotides which represents a robust honeycomb DO stapling motif (fig. S9) (*23, 35, 36*). Here, hard constraints are used to ensure staple breaks are not placed near scaffold or staple crossovers to prevent kinetic traps (supplementary text and fig. S10 and S11). Due to the dynamic nature of the presented algorithms, each input condition must be individually verified to ensure compatibility.

Generally, the input graph must be connected, contain only straight edges, and have edge lengths of at least 42 nucleotides (∼14 nm), although the exact minimum depends on the bundle thickness and the angles between adjacent edges (supplementary text). Together, the described abstraction enables a simple input of vertices, edges, and a cross-section parameter to be compiled directly into a DNA origami building block (Fig. 1 A to C), observed across hundreds of input conditions (select examples in Fig. 2 and figs. S12 and 13).

**Fig. 2.**
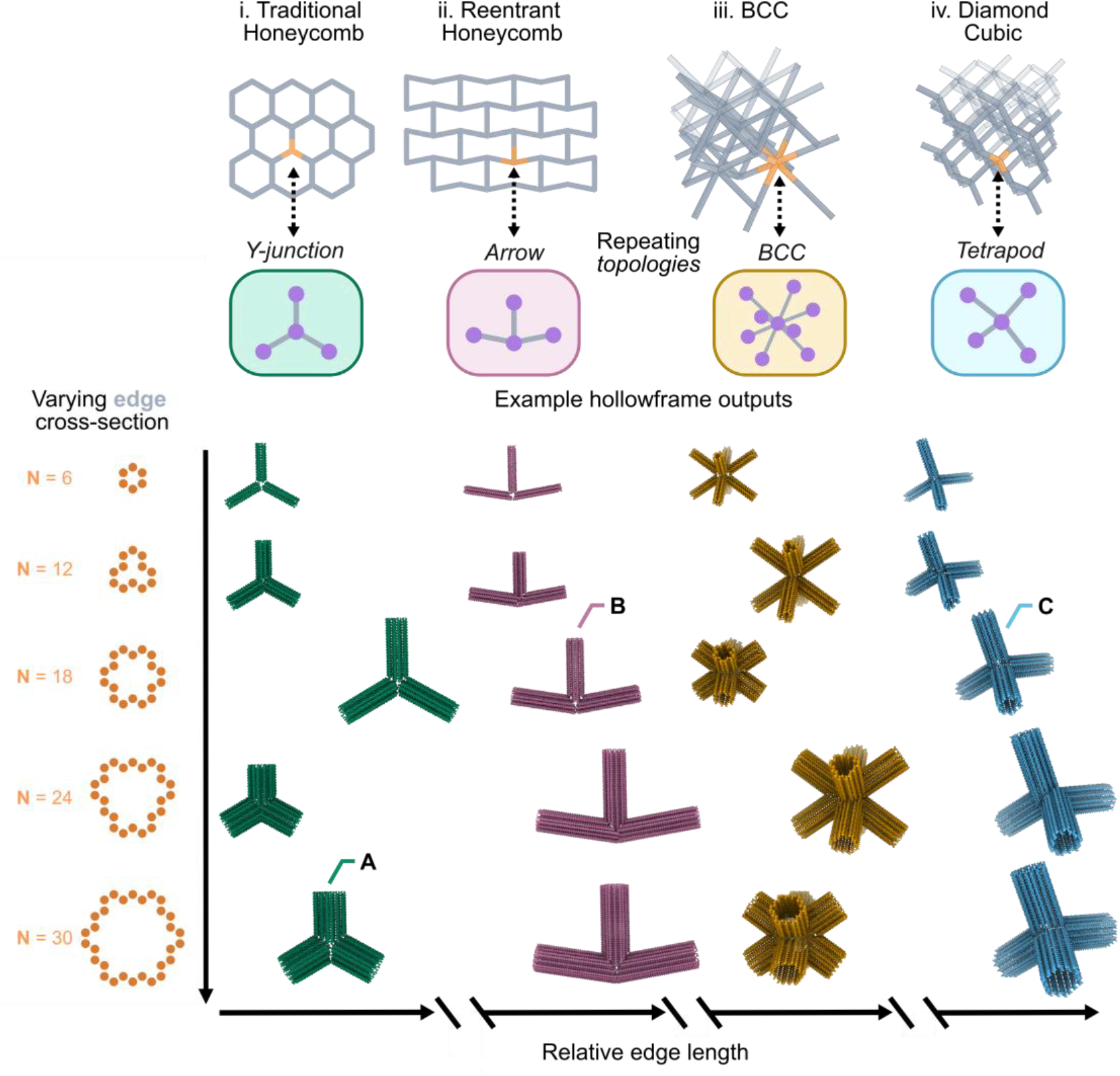
Exploring hollowframe topologies by breaking down periodic material into repeating geometries. (i) Traditional and (ii) reentrant honeycombs alongside (iii) body centered cubic (BCC) and (iv) diamond cubic lattices are shown. Each lattice is broken down into a repeating topological unit that is coupled alongside a cross-section to create hollowframe DNA origami nanostructures spanning a design space of stiffnesses and edge lengths. Designs selected for characterization (Fig. 3) are the (**A**) 30 HB Y-junction, (**B**) 18 HB arrow, and (**C**) 18 HB tetrapod.

### Automated realization of stiffness-programmable particles

In contrast to fixed-paradigm automation algorithms (*20, 24*), the hollowframe approach enables designers to explore a less-constrained design space by coupling both geometry and bundle cross-sections within a shared pipeline (Fig. 1), thereby exposing a broader spectrum of candidate DO building blocks for metamaterials design (Fig. 2). Increasing the number of DNA helices raises the predicted per-edge bending stiffness (fig. S1) and produces systematically lower RMSF and reduced global deformation as shown through oxDNA (*29*) simulations (fig. S14 - S15). However, there is a tradeoff as a larger number of DNA helices consumes a larger fraction of the scaffold, thereby constraining the achievable nanostructure size from commercially available scaffolds. Here we address this issue by utilizing hollow honeycomb ring-like cross-sections of near-maximal enclosed area (Fig. 1 B i) to provide favorable bending stiffness to scaffold use ratio of each edge bundle. To assess manufacturability, we selected three structurally distinct topologies to use with the presented hollowframe automation paradigm (Fig. 1): an 18 HB arrow, an 18 HB tetrapod, and a 30 HB Y-junction (Fig. 2 A, B, and C and supplementary tables S1 – S3).

All three structures fold using a shared 24-hour annealing protocol in standard DO buffer conditions without requiring structure-specific ramp optimizations (see Materials and Methods). Atomic force microscopy (AFM) and negative-stain transmission electron microscopy (TEM) (fig. S16 – S18) resolve the unique structural features of each design, confirming that the presented automated workflow faithfully realizes the designed nanoparticles (Fig. 3). While preferred orientation and low particle concentration restrict 3D reconstruction, Cryo-TEM 2D (*37, 38*) classification (Fig. 3 and fig. S19 – S21) complements these findings showing class averages that appear qualitatively well-aligned with their input geometries from a more natural and vitrified state. Collectively, these results support that the mitering algorithm correctly tunes the thick helix bundle topology to ensure a conformal fit at shared vertices across the tested nanostructures. A magnesium salt sweep reveals that hollowframes form distinct monomer bands between 8 and 20 mM MgCl_2_ but are most reliably folded between 12 to 16 mM MgCl_2_ with minimal aggregation as seen by agarose gel electrophoresis (Fig. 3 and fig. S22). More generally, these characterizations establish the experimental robustness and accessibility of hollowframe DNA origami monomers.

**Fig. 3.**
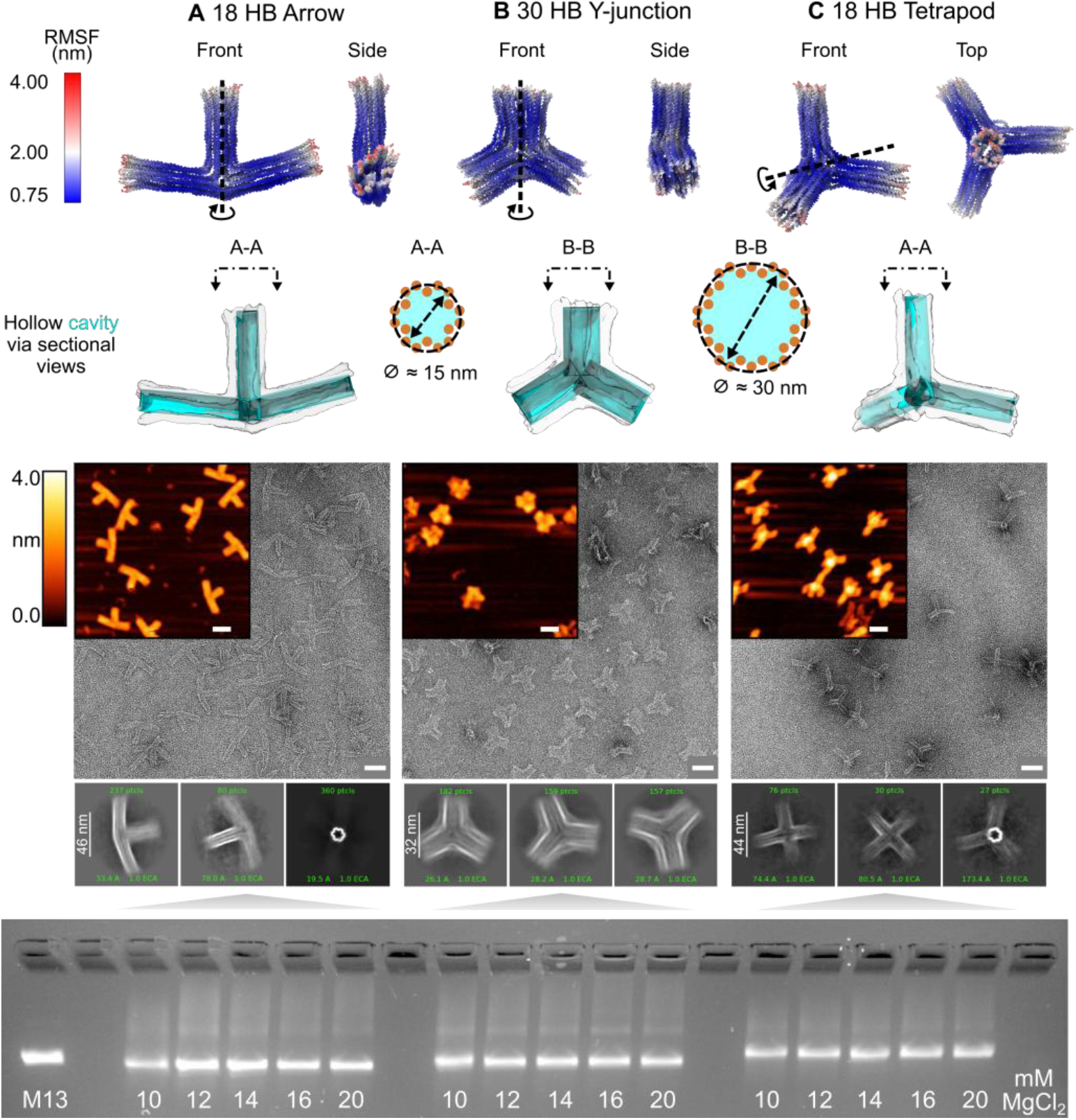
Experimental characterization of three distinct nanostructures. Three hollowframe nanostructures, (**A**) 18 HB arrow, (**B**) 30 HB Y-junction, and (**C**) 18 HB tetrapod are shown (*30*) in their centroid configurations (*39*) as simulated by oxDNA (*29*). Sectional views are provided to show the internal cavity (cyan) of these nanoparticles resulting from the hollow cross-section diameter. Atomic force microscopy (AFM) and negative-stain transmission electron microscopy (TEM) show robust nanostructure formation and select cryo-TEM class averages (*37, 38*) show the structure in a more natural, vitrified state (figs. S19-S21 for full class averages). Agarose gel electrophoresis shows well-folded DNA origami across a range of magnesium salt concentrations (fig. S22). All scale bars are 50 nanometers unless otherwise specified.

### Assembling micron-scale 2D DNA origami materials

With monomer hollowframe DO nanostructures established, investigation turns to their ability to be assembled into periodic lattices. Complementary sticky ends provide a direct route to programmatically orient and position DO building blocks for colloidal assembly (*4*– *6*). NUPACK 4 (*40*) is used to screen for candidate sticky end sequences with similar melting temperatures and evaluate the free energy differences between candidates. A GPU-based simulated annealing algorithm (*41, 42*) (supplementary text) searches the vast design space for a set of sticky ends whose off-target interactions are well separated from the on-target sticky end interactions (supplementary text and fig. S23 and S24). The 18 HB arrow and 30 HB Y-junction use six and ten sticky ends per face, respectively. The remaining helices on each face contain 3-nucleotide long poly-T single-stranded DNA brushes to discourage off-target blunt end interactions (*43*). The sticky end and poly-T overhangs are combined with the core staple strands alongside standard DO buffer conditions to assemble these 2D periodic materials using a one-pot, three-day thermal anneal (see Materials and Methods).

AFM and negative-stain TEM reveal the robust formation of micron-scale sheets in both systems. The 30 HB Y-junction assembles into a conventional honeycomb lattice (Fig. 4 A and fig. S25) while the 18 HB arrow yields the programmed reentrant honeycomb lattice (Fig. 4 B and fig. S26). These lattices are architecturally unique and exhibit distinct simulated mechanical responses (Fig. 5) in part due to their underlying geometries (*14*).

**Fig. 4.**
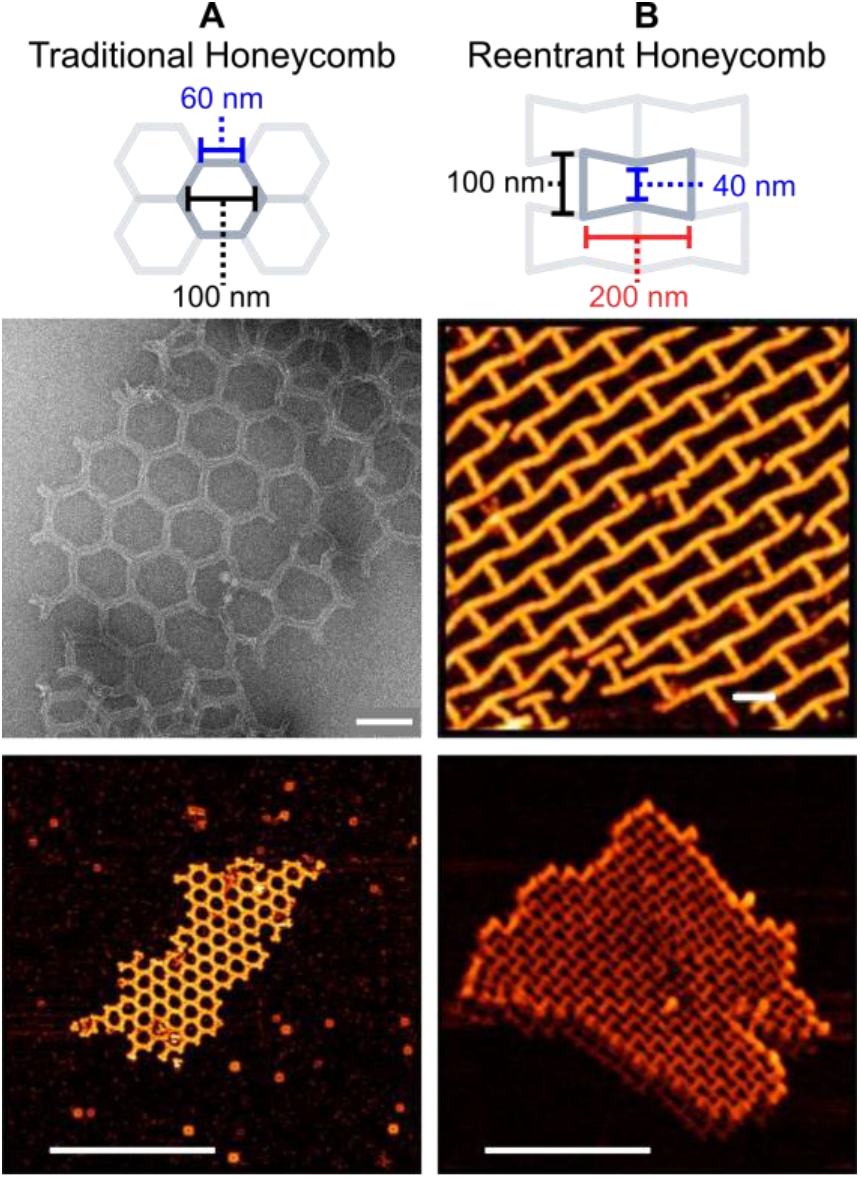
Experimental characterization of 2D DNA origami lattices. Micron-scale (**A**) traditional and (**B**) reentrant honeycomb lattices are designed with nanometer-level precision and assembled by one-pot assembly (see Materials and Methods) as shown by AFM and negative-stain TEM. Scale bars are 100 nm (top row) and 1 µm (bottom row).

**Fig. 5.**
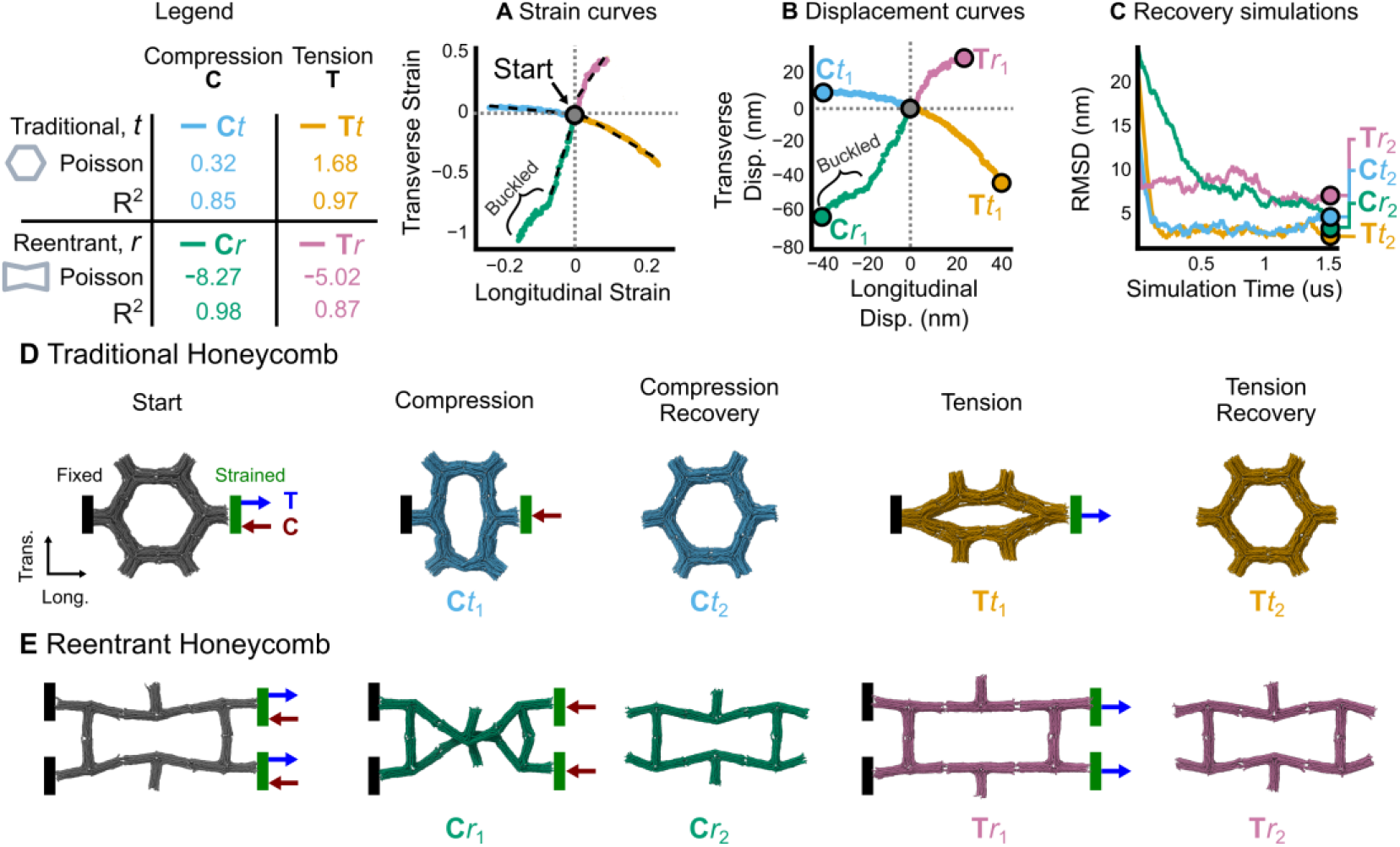
CG MD simulations of honeycomb unit cells under uniaxial strain. (**A**) Transverse versus longitudinal strain profiles and (**B**) displacement curves for traditional and reentrant honeycombs under tensile or compressive loading. The reentrant honeycomb exhibits negative Poisson ratio behavior and structural buckling yet (**C**) subsequent unconstrained simulations show global structural recovery toward the initial state. Structural snapshots for the (**D**) traditional and (**E**) reentrant honeycombs are shown for loaded and recovered states (figs. S27 to S31).

Furthermore, both lattices are generated from a shared design automation and optimization workflow that treats an input geometry and cross-section as coupled and tunable parameters (Fig. 1 and 2 i and ii). Their successful realization demonstrates how the hollowframe paradigm provides access to a 2D DNA origami metamaterials design space in which the building-block geometry, nanostructure stiffness, and sticky-end connectivity can be programmed together.

### Simulation-informed mechanical response of architected materials

Coarse-grained molecular dynamics (CG MD) oxDNA simulations (*29, 44*) quantify unit cell response under displacement-controlled uniaxial loading. Harmonic trap forces (see Materials and Methods) impose both compressive and tensile displacements on individual traditional and reentrant honeycomb unit cells (supplementary movies S1 and S2, respectively). While conventional bulk isotropic materials typically have Poisson ratios between 0 and 0.5 (*45*), an architected lattice can be characterized by a geometric ratio capturing the kinematics of the unit cell under displacement and do not need to fall within this range. Here, the Poisson ratio is found as the negative slope of the linear regression between transverse and longitudinal strain data fit within the linear-response regime. Strains are found by tracking relative spatial displacements between averaged nucleotide groups along the principal structural axes throughout the simulation (fig. S27).

The traditional honeycomb exhibits a positive Poisson ratio, meaning a negative slope of transverse to longitudinal strain (Fig. 5 A **C***t* and **T***t*), throughout the simulation (Fig. 5 D). This results in the multi-DO unit contracting transversely under tension while expanding under compression. The lattice can structurally undergo large amounts of deformation due to the thickness of the 30 HB cross-section, and the honeycomb does not buckle out of plane in the allotted simulation time (fig. S28). Furthermore, the traditional honeycomb rapidly recovers its initial, unstrained state when simulated with the enforced strain conditions removed (Fig. 5 C, fig. S29, and supplementary movie S3).

Conversely, the simulated reentrant honeycomb exhibits auxetic behavior, having a negative Poisson ratio with a positive slope of transverse to longitudinal strain (Fig. 5 A **C***r* and **T***r*). An auxetic is a class of mechanical metamaterial that expands transversely under tension while contracting under compression (*14*). However, under high load, the reentrant material undergoes out-of-plane buckling in compression and exhibits a structural failure in tension as the sticky end connections between building blocks dissociate during simulated straining (Fig. 5 E and fig. S30). Interestingly, the buckling is largely reversible over the simulated timescale, as the global RMSD tends toward the initial reference configuration after the strain conditions are removed (Fig. 5 C, fig. S31, and supplementary movie S4).

This suggests these DO lattices can recover their initial global conformation even under severe mechanical strain, provided sticky ends remain intact. Simulations of a 30 HB arrow assembled into the reentrant honeycomb using ten sticky end connections per face show that the increased stiffness of the thicker cross-section shifts the onset of buckling deeper into the compression window, however, it also structurally fails in tension (fig. S32). Together, these results show that hollowframe design choices produce distinct unit-cell responses under simulated loading conditions.

## Discussion

We have introduced a fully automated hollowframe paradigm that treats a graph-based geometry and edge stiffness as coupled design variables and directly translates them into manufacturable nucleotide-level structures, expanding the accessible structure-property space for DO materials beyond fixed mesh paradigms (*19, 20*). Owing to a strict geometry-dependence of the underlying automation algorithms (Fig. 1), hollowframe nanostructures are extendable to non-honeycomb cross-sections and allow per-edge N HB counts to be specified (fig. S12, S13, and S33). Pairing this paradigm with a recent geometry-independent assembly algorithm (*5*) could enable researchers to sculpt complex, nano-to-micronscale lattices using DNA origami without requiring extensive designer expertise.

Simulations predict that hollowframe DO unit cells can structurally undergo uniaxial deformation and exhibit effective behaviors such as negative Poisson’s ratio (Fig. 5). Small amounts (∼2 to 5 nm) of lateral displacement, potentially activated through strand displacement mechanisms (*13, 46*), are mechanically amplified into larger perpendicular displacements (∼5 to 13 nm) under tension (Fig. 5 B). This behavior could be incorporated into discrete hollowframe unit cell structures, as opposed to periodic materials (Fig. 4), to engineer nanoscale mechanical testing apparatuses (*47*). The automation package is provided as free open-source software that can be accessed through an online, web-based GUI or through a Python API (*48*).

## Acknowledgments

We would like to thank Dr. James Conway at the University of Pittsburgh School of Medicine for conducting the TEM imaging. The Pittsburgh Center for CryoEM (RRID:SCR_025216) used for data collection in this project was supported, in part, by the University of Pittsburgh, the School of Medicine, the Department of Structural Biology, and the National Institutes of Health (grants S10-OD-019995 and S10-OD-025009). The content is solely the responsibility of the authors and does not necessarily represent the official views of the National Institutes of Health. Molecular graphics and analyses performed with UCSF ChimeraX, developed by the Resource for Biocomputing, Visualization, and Informatics at the University of California, San Francisco, with support from National Institutes of Health R01-GM129325 and the Office of Cyber Infrastructure and Computational Biology, National Institute of Allergy and Infectious Diseases. We also would like to thank Anuhya Adupuganti, Isabella Ferranti, Logan Carpenter, Mangalam Sahai, Praneetha Prakash, and Vismaya Walawalkar for their thoughtful input and assistance in the development of this work.

## Author contributions

Conceptualization: AJV, JC, RET

Methodology: AJV, JC, RET

Investigation: AJV

Visualization: AJV

Funding acquisition: AJV, JC, RET

Project administration: JC, RET

Supervision: JC, RET

Writing – original draft: AJV, JC, RET Writing – review & editing: AJV, JC, RET

## Data, code, and materials availability

The data, code, and materials underlying this article are available within the article itself and in its online supplementary materials. The actively maintained code and web-based GUI are freely available online via GitHub (https://github.com/ajvetturini/hado) alongside API documentation. The initial version of the code and data collection files has been permanently deposited into Zenodo and are available at https://doi.org/10.5281/zenodo.21515951.

## Supplementary Materials

Materials and Methods

Supplementary Text

Figs. S1 to S33

Tables S1 to S3

References (*22, 23, 29, 31, 33–42, 44, 48–52*)

Movies S1 to S4

sData Files S1

