## supplementary text for "Automated design of stiffness-tunable DNA origami hollowframes for self-assembling metamaterials"

### Materials and Methods

Individual DNA origami sample preparation

The required staple single stranded DNA sequences for the 18 HB Arrow, 30 HB Y-junction, and 18 HB Tetrapod (supplementary table S1, S2, and S3, respectively) were produced using the design automation framework proposed in this work. Staples were ordered in 96-well plates at 200 µM concentration in IDTE (8.0 pH) (Integrated DNA Technologies, Coralville, Iowa, USA). All non-sticky end staples (i.e., staple oligonucleotides labelled *core* or *PolyT* in tables S1-S3) were combined in equal parts to form a staple mix calibrated to 1000 nM. All structures shown used the M13mp18 scaffold (Bayou Biolabs, Metairie, Louisiana, USA) at 450 nM concentration. For annealing of the nanostructures, 10 nM of the scaffold was combined with 100 nM of the staple mix into 1X TAE (8.3 pH) and 16 mM $MgCl_2$. Finally, all structures were annealed using the following thermal ramp over the course of approximately 24 hours: 95°C for 5 minutes, 70°C for 15 minutes, 70°C to 30°C at -1°C per 30 minutes, 30°C to 20°C at -1°C per 5 minutes and then hold samples at 4°C before refrigerating.

One pot assembly of DNA origami 2D materials sample preparation

The 18 HB arrow and 30 HB Y-junction are geometrically capable of tiling to a reentrant and traditional honeycomb lattice, respectively. The designed sticky end staples (supplementary text) were ordered in 96-well plates at 200 µM concentration in IDTE (8.0 pH). These staples (i.e., those labelled with a *color* in supplementary table S1 and S2) were then combined into a binding staple only mix calibrated to 1000 nM. To accomplish the assembly, a one-pot anneal was conducted using 10 nM of the scaffold combined with 100 nM of the core staple mix and 75 nM of the binding staple mix into 1X TAE (8.3 pH) and 12 mM $MgCl_2$. An approximately 3-day long annealing ramp (95°C for 5 minutes, 65°C for 2 hours, 65°C to 35°C at -0.5°C per 1 hour, 35°C to 20°C at -1°C per 15 minutes, then refrigerate and hold samples at 4°C) was used to create the materials shown (main article, Fig. 4).

Amicon purification

The individual DNA origami samples were purified from the excess staples and folding buffer using an Amicon Ultra 0.5 mL spin filter column with a molecular weight cutoff of 100 kDa (Thermo Fisher Scientific, Waltham, MA, USA). First, the filter was soaked overnight in 500 µL of a 5% Tween-20 (MilliporeSigma, Burlington, MA USA) mix. Then, the Tween mix was carefully aspirated out without damaging the filter membrane. The filter was then washed using 500 µL of autoclaved reverse osmosis deionized (RODI) water and spun at 10000 rcf for 3 minutes. The spin-through fluid was discarded, and the process repeated for 5 total washes. The filter was then filled with 500 µL of a low salt buffer (LSB) (1X TAE, 5 mM $MgCl_2$) and spun at

10000 rcf for 3 minutes. Then, 100 µL of DNA origami sample and 400 µL of LSB were added and spun at 2000 rcf for 15 minutes. Finally, samples were washed 3 more times at 3000 rcf for 5 minutes each using 500 µL of the LSB per-wash before flipping the filter upside down into a new tube and spinning at 3000 rcf for 3 minutes to extract the purified sample.

Gel electrophoresis

For all DNA origami samples, 15.30 µL of the 10 nM annealed sample was combined with 1.70 µL of 10X gel loading buffer (Thermo Fisher Scientific, Waltham, MA, USA). Then, 15 µL of this solution was analyzed by electrophoresis in 2% agarose gel in 1X TBE buffer with 12.5 mM $MgCl_2$ and 1X SYBR Safe DNA gel stain (Thermo Fisher Scientific, Waltham, MA, USA) run at 100 V for 75 minutes. The gels were imaged using the ChemiDoc Imaging System (Bio-Rad, Hercules, CA, USA).

Atomic force microscopy (AFM) sample preparation

For AFM, either purified or unpurified DNA origami sample was diluted to 20 µL at 2 nM using buffer matching the sample salt conditions. First, a mica sheet was fixed to a metal wafer (Park Systems Corporation, Gwacheon City, South Korea). The sample was then deposited onto a freshly cleaved mica surface and incubated in a humidity chamber for 5 minutes. The samples were then blow-dried with nitrogen and 75 µL of RODI water was deposited and let sit for 5 minutes. This step was repeated for 2 total washes before being scanned using the NX10 AFM system (Park Systems Corporation, Gwacheon City, South Korea) in non-contact mode.

Negative stain transmission electron microscopy (TEM) sample preparation

For negative stain TEM, 2.5 µL of the purified DNA origami samples were deposited onto a freshly glow-discharged copper grid with a support film of graphitized carbon. The sample was side-blotted, washed on a drop of water, blotted again, and washed in a 1% solution of uranyl acetate before a final blot. Grids were air-dried for 5 minutes before imaging in a Tecnai TF20 transmission electron microscope operating at 200 kV (Thermo Fisher Scientific, Waltham, MA, USA). Images were collects on a TVIPS XF416 CMOS camera using the TVIPS Emplified software (TVIPS GmbH, Gilching, Germany).

Cryo-TEM sample preparation and image processing

For cryo-electron microscopy, 2.5 µL of the purified DNA origami samples were deposited onto freshly glow-discharged Quantifoil R2/1 grids (Quantifoil Micro Tools GmbH, Großlöbicha, Germany), blotted and plunge-frozen into a 60:40 mixture of liquid propane:liquid ethane using a TFS Vitrobot Mk 4 (51). Vitrified grids were clipped into Autogrids and inserted into a TFS Krios TEM operated at 300 kV and equipped with a Selectris energy filter and Falcon 4i direct electron detecting camera. Movies were collected in electron counting mode under the control of the TFS EPU software using a total dose of ~30 e/Å$^2$. Particles were imaged at 130kx magnification, corresponding to a pixel size of 1.17Å at the sample.

High-resolution EER files from the Cryo-TEM collection were first imported into CryoSPARC (36) and preprocessed via patch motion correction and CTF estimation using the default import settings. Micrographs with poor ice quality or zero-particle presence were then excluded prior to particle picking. An initial set of 100 particles were manually picked to train a Topaz (37) model for each TEM dataset that was then used to extract the particles from the remaining micrograph stack. Overall, low particle concentrations were observed across all three nanostructures, and therefore 3D reconstruction was not pursued. Instead, a 2D classification of the Topaz-picked particles across 50 classes was used to qualitatively compare particle morphology alongside AFM and negative-stain TEM.

Coarse-grain molecular dynamics (CG MD) simulations

CG MD simulations are used to systematically probe the structural behavior of the nanostructures and materials in this work. Structures were first minimized for 1000 timesteps to resolve physical overlaps in the initial simulation model. Then, a relaxation step equilibrates the structure for 1e6 timesteps and a Langevin thermostat. Following the equilibration, the structure was simulated for 5e7 timesteps using the oxDNA2 model (29, 43) with a salt concentration of 1.0 and was performed on an RTX A6000 (48GB VRAM) GPU before being analyzed using the oxDNA analysis toolkit (38). The minimization, relaxation, and simulations all use a time step of 0.005 and example oxDNA input files are provided in the deposited code repository (47).

Strain simulations (main article, Figure 5) were performed on a single unit cell that was manually connected via the programmed sticky end connections using oxView (38). This unit cell was then simulated using the above described minimize-relax-simulate procedure and the centroid configuration was evaluated using oxDNA analysis tools. The centroid was then aligned with the global coordinate system such that the strain is applied in the ±X direction (fig. S27 A). Fixed harmonic trap forces (with a rate of 0) were placed on the single stranded DNA nucleotides that would otherwise be connected via sticky ends to connecting unit cells. Harmonic traps with a rate of 1e-6 in the X (i.e., [1, 0, 0]) direction were placed on the opposing ends sticky end nucleotides to simulate a strain test (fig. S27 B). These simulations were run for 1e7 timesteps and trajectories were analyzed via custom scripting to measure the Poisson ratio (fig. S27 C). Lastly, for recovery simulations (main article, Figure 5), the initial state is simply the final configuration from the strain simulation. No minimization or relaxation is necessary from this strained state, and thus standard oxDNA simulations were performed for 5e7 timesteps using a time step of 0.005. The RMSD reported by the recovery simulations (main article, Fig. 5 ii) is the Kabsch-aligned RMSD between the pre-strained centroid configuration and the trajectory of the recovering material.

### Supplementary Text

Defining the input geometry

This framework takes two key inputs: 1) An input wireframe model represented as a graph (referred to as WM in what follows) and 2) an edge thickness parameter. The WM contains nodes representing 3D spatial Cartesian coordinates in units of nanometers and edges defining the connectivity between the vertices. The WM input is not restricted to closed surface meshes, and arbitrary 3D wireframe networks defined by interconnected line segments are valid. The edge thickness parameter determines the number of DNA helices that will populate each edge of the WM. This number, N, of DNA helices is then arranged into a helix bundle (i.e., N HB) that is aligned with each edge of the WM.

There are a few limitations to enable successful conversion from the input representation to the nucleotide-level nanostructure. Firstly, the edge lengths in the input WM must be sufficiently long, typically at least ~14 nanometers (42 nucleotides presuming a helical rise of 0.34 nm per basepair for B-DNA) long. Overall, the actual minimal edge length will depend on a combination of the bundle thickness (i.e., how many DNA helices per edge) and the angles between edges sharing a vertex. Due to the dynamic nature of the mitering and stapling algorithms, the actual minimal edge-length will therefore be specific to the input geometry. Furthermore, the input WM, when represented as a graph, must contain a single connected component. Lastly, all edges are considered as straight lines, and curvilinear input WMs (i.e., DNA origami with curved bundle features) are not yet supported.

<u>Converting an edge thickness to DNA helix bundle cross-sections</u>

The first step in converting the WM-and-thickness input into a DNA origami nanostructure is to define the cross-section of the DNA helices along each edge. A honeycomb grid was selected to represent the geometry of the cross-section due to its reliable design motifs (helical pitch of 10.5 bp/turn and crossovers every 7-nucleotides, depending on the neighboring helices). While other grid styles can be used, such as the square grid (fig. S33), they require more careful validation to account for staple design features that may result in severe global structural twist.

Designers can specify the edge thickness as either a target outer diameter (in units of nanometers) or a desired number of DNA helices to place along the edge (defined as an integer value greater than or equal to 2). If a diameter is specified, then a contiguous honeycomb ring with an even total number of DNA helices is found whose diameter is closest to the specified diameter (fig. S1 A). This diameter is computed using the center-to-center spacing of the DNA helices, which in this work is defined by the diameter of DNA (assumed to be 2.25 nanometers) added to an inter-helix spacing distance due to electrostatic repulsion of the negatively-charged DNA backbone (assumed to be 1.00 nanometers) (fig. S1 A). Here, it's noted that this work focuses on constant thickness cross-section inputs along each edge, although the underlying algorithms support per-edge assignments (fig. S12 and S13).

A designer may otherwise specify the per-edge thickness as a discrete number of DNA helices. If the specified number of helices is even, then the scaffold routing algorithm (next section) can guarantee a single continuous scaffold routing without checking the WM topology (fig. S2 A). Otherwise, this algorithm also supports specifying an odd number of DNA helices per edge that is approximated using a custom graph heuristic (fig. S2 B). First, each edge is presumed to have $\lfloor N/2 \rfloor$ helices running in opposite directions. For example, if N was 3, then each edge in the graph is first aligned with anti-parallel running helices. An Eulerian tour of the original graph would provide a perfectly matching scaffold routing from this stage, as each edge is visited once while returning to a start vertex (ensuring scaffold circularity), and thus the target N would be observed on each edge. However, the input WM topology will dictate if an Eulerian tour is topologically possible, and if not, then a few heuristics are used to approximate a tour. First, the WM is checked to ensure that it does not contain any pendant (degree-1) vertices, as those would represent an edge that would always require two traversals (out-and-back) to cover (fig. S2 A), thus resulting in a required even-number of helices for that edge. Presuming there are no degree-1 nodes, then the first heuristic constructs a metric closure of the WM and applies Christofides' algorithm (50) to find a vertex-visiting routing that minimizes extra edge visits. The second approach approximates an Eulerian tour by iteratively extracting cycles via depth-first traversals and stitching them together at shared vertices to form a global tour. Any cycles that cannot be stitched through a shared vertex are discarded. The two solutions are then compared to see which better approximates the target number of helices per edge (fig. S2 iv).

Regardless of the input mechanism, a honeycomb-like ring of DNA helices is populated where each helix has a known Euclidean (X, Y) position (fig. S3). The centroid of the cross-section is then translated to align with an arbitrary local origin of (0, 0). Then, each helix position has a Z dimension appended (Z=0) as the cross-section will eventually be translated and rotated into 3D space along the WM edges in future steps. Lastly, it's noted that while this work focuses solely on honeycomb grids with at least 2 DNA helices (i.e., 2 HB) for the edge cross-sections, designers may specify their own custom positionings using the Python API (47). Currently, this is only possible if scaffold and staple crossovers occur at regular intervals for each turn of DNA for the custom cross-section specified. However, any structural artefacts, such as global twist, that may be induced by custom motifs must then be individually tuned.

Connecting a continuous scaffold routing

Once the cross-section(s) for each edge are defined, a continuous, single scaffold routing can be determined. First, a new graph is initialized to represent the nucleotide-level abstraction of the input WM. The number of nucleotides equal to the edge length divided by the helical rise in B-form DNA (0.34 nanometers) is calculated. Each helix within a cross-section is specified using this number of nucleotides, extending along the +Z direction. These nucleotides are stored as graph nodes and the edges connecting individual helix runs of nucleotides are set in the nucleotide level graph. Then, the spatial positions of nucleotides are translated and rotated to align with the corresponding WM edge using the Rodrigues' rotation formula. This process is repeated for all edges in the input WM (fig. S4 A).

Next, each bundle of nucleotides is rotated about its edge as the central axis in discrete angular values. At each vertex containing at least two edges, a matching problem is constructed to find a rotation that minimizes a scaffold traversal cost between helices. Here, helices are first partitioned into two sets based on the directionality of DNA corresponding to the 3' and 5' ends of the individual helices. A cost matrix is then calculated based on the pairwise Euclidean distances between the two sets (fig. S4 B). The Hungarian algorithm is used to identify the set of helix pairings that minimizes the total Euclidean distance cost at a vertex in polynomial time (48). A greedy stochastic search is used to identify a state containing a discrete rotation per edge that minimizes the resulting scaffold traversal cost. This search is computationally efficient as the honeycomb cross-section exhibits 60-degree rotational symmetry, restricting the number of distinct rotation states that must be considered to find a quality state that does not have the scaffold traversing unphysically large distances (fig. S4 B).

The local pairings found by the Hungarian algorithm above do not yet define a complete, single global scaffold routing. To resolve this, an adjacency graph is defined whose nodes represent the cycles formed by the paired helix segments and whose edges represent cycles that can be combined using an internal scaffold crossover. A minimum spanning tree algorithm (fig. S8) is used to collapse the many local cycles found into one continuous scaffold routing through use of internal scaffold crossovers, similar to the procedure described by MagicDNA (22). Briefly, the adjacency graph has edge weights corresponding to a count of the number of scaffold crossover locations that connect two cycles into one. The minimum spanning tree ensures that the most constrained (i.e., lowest-crossover count) sub-cycles are first connected prior to the lesser constrained regions. Lastly, this work's scaffold routing algorithm can support any combination of even DNA helices per edge as well as the heuristic determined even-odd (or odd-only) helix edges (fig. S2 A). At this point, the per-helix nucleotide positions are known but the internal crossovers are not yet assigned as the design must first be mitered which will selectively add and remove more nucleotides to the helices.

Geometric mitering for global stiffness

With the helix connections and scaffold routing determined, a geometric mitering algorithm finetunes the ends of the helix pairs (defined by the Hungarian algorithm in the previous section). At each multi-edge vertex, the mitering algorithm begins looping over all paired helices starting from the mid-point of the edge. Nucleotides are kept, added, or removed one base at a time along the helical axes using the axial rise value of 0.34 nanometers until the paired endpoints reach a Euclidean distance, $d_{cut}$, of 2.0 nanometers (fig. S5). The final distance found is then divided by a presumed nucleotide-to-nucleotide spacing of single stranded DNA (ssDNA), $d_{ssDNA}$, to determine a discrete number of single stranded scaffold nucleotides placed at the 3' end of the connected helices (fig. S6 A). The use of single stranded overhangs reportedly reduces torsion experienced in the backbone and enables structural uniformity (31).

While previous work used a reported value of 0.42 nanometers for $d_{ssDNA}$, the described work uses 0.60 as determined through oxDNA simulations across a variety of input geometries (fig. S7). These simulations tested combinations of final point-to-point Euclidean distances (1.0, 1.5, 2.0, 2.5, 3.0 nanometers) and ssDNA distances (0.42, 0.48, 0.54, 0.60 nanometers). The root mean square deviation (RMSD) between the centroid configuration of a simulation and the input geometry was used to infer how deformed a nanostructure is. Having lower RMSD generally results in a more structurally stable design that is more suitable for constructing periodic materials. While no clear choice in parameters is seen across all geometries, simulations show that a lower RMSD across designs was found when the final point-to-point $d_{cut}$ distance was set to 2.0 nanometers and a single stranded DNA spacing, $d_{ssDNA,}$ of 0.60 nanometers was used, however no obvious trend is observed between the variables (fig. S7).

Once mitering is applied to each multi-edge vertex, the scaffold crossovers can be determined at the individual nucleotide-level using the periodic motifs of the honeycomb grid. Internal crossovers found from the minimum spanning tree (fig. S8) are located centrally along the individual helices in each bundle along each edge. Scaffold crossover positions are determined by the periodic crossover rules (every 7 nucleotides) of the honeycomb lattice resulting in a nucleotide-level graph abstraction containing a single, continuous scaffold routing.

Automated stapling algorithm

With the scaffold completely defined, stapling can be performed to complete the nucleotide-level design representation. First, staple nucleotides are placed onto the design pairing with the scaffold nucleotides, excluding the single stranded spacer regions. Then, a number of single stranded poly-T overhangs are then added to the 3' ends of the staples. This is found by dividing the final Euclidean distance found from the mitering step by 0.60 nanometers, as described in the previous geometric mitering subsection (fig. S6 B). Then, the staple 3' and 5' ends connecting the staple nucleotides between helix bundles are connected using the helix connections found by the Hungarian algorithm in the scaffold routing subsection (fig. S9 A - C).

Next, all allowable staple crossovers between neighboring helices within a bundle are introduced, which generally produces long, circular staples (fig. S9 D). However, there are three constraints imposed when placing crossovers: 1) no crossovers are placed in the single-stranded poly-T overhangs, 2) no staple crossovers are placed within 5 nucleotides of the ends of the double-stranded staple segment and 3) staples must run (i.e., bind with the scaffold) for at least 5 nucleotides after crossing over from another helix before another crossover can be taken (fig. S10 A). Generally, the value of 5 nucleotides was chosen to encourage enough binding between staple-scaffold nucleotides before entering another kinetic event (i.e., connections between two helix bundles or internally within a bundle).

The resulting long staples must then be broken down into lengths between a lower and upper bound (LB and UB). By default, LB is 20 and UB is 60 nucleotides, as these are currently the bounds of synthetic single stranded DNAs that can be reliably manufactured at a standard cost. This is accomplished using a staple auto-break algorithm that minimizes the squared deviation from a target staple length of 42 nucleotides. The value 42 represents a honeycomb-grid DNA origami staple motif that has been shown to have improved assembly yields due to continuous 14-nucleotide duplex domains flanked by the repetitive staple crossovers between neighboring helices (23, 34, 35). The first step in the auto-break is to identify all valid nick positions along all long staple runs. The following positions are excluded: 1) in the single stranded domain of a staple, 2) within 3 nucleotides of crossing over to encourage binding and 3) within 3 nucleotides of any scaffold crossovers on the same helix (fig. S10 B). This 3-nucleotide exclusion follows a recent automated thermodynamics-driven stapling algorithm (33) that cannot be directly applied to hollowframe structures as it relies explicitly on thick lattice-based packing motifs.

Once the nick positions have been validated, the staple breaking begins looping over all long staples and treating each individually as a one-dimensional partitioning problem. This is solved using a dynamic programming (DP) approach, where a DP table is constructed in which each entry stores the minimum cumulative penalty for placing a given number of nicks up to a given candidate position. Here, the penalty assigned to a staple segment that is spanning between two candidate nick positions, $n_i$ and $n_j$, is

$$C(n_i, n_j) = \left((n_j - n_i) - L_{target}\right)^2.$$

This cost function favors staples whose lengths most closely match the target length of 42 nucleotides. Segments shorter than LB (20 nucleotides) are disallowed and their cost set to infinity, whereas segments longer than UB (60 nucleotides) are assigned a large penalty so that they are avoided unless no valid alternative (i.e., less than 60 nucleotides nick point) exists. The constraint values used as described in this section come from a combination of literature-defined heuristics (33) alongside an ablation study of their ranges (fig. S11). If the constraints are set to be overly restrictive (i.e., too large), then the cost explodes as no nick points exist. Conversely, removing the constraints enables the cost function to be further minimized, however, introduces potential kinetic trapping due to poor staple design. Therefore, the constraint values were set to be as large as possible (to encourage maximal binding between scaffold and staple) while not being prohibitively large across the designs analyzed. Once the DP table is filled in, the optimal set of nick positions is recovered by backtracking from the minimum-cost terminal state. This yields the partition of the staple into more uniform lengths, and then this partition procedure is repeated for the remaining staples (fig. S10 C).

Computational design of sticky end sequences

Sticky end sequences are used to programmatically assemble the DNA origami units into a prespecified lattice. To accomplish this, NUPACK 4 (39) is used to inform a heuristic-driven search (40) for a set of sticky end sequences that are orthogonal as defined by the NUPACK ensemble free energy. The search space here is constrained using three parameters: the sequence length, a target melting temperature, and a specified number of GC nucleotides included in the sequence. The melting temperature here is used only to screen for thermodynamically similar sticky end overhangs and should not be interpreted as the melting temperature of assembly.

First, all candidate sequences are enumerated, which is computationally feasible because there are only 4 possible nucleotides and the max length a sticky end for assembly is typically 8 nucleotides (i.e., $4^8$ = 65,536 potential sequences). Longer sticky ends are generally avoided as their melting temperature tends to grow too large that leads to assembly defects that lock in early on during the thermal annealing ramp. These 65,536 sequences are first screened down by ensuring sequences do not contain any more than three consecutive nucleotides (e.g., GGGGAAAA and TTTTCCGG would be invalid). Then, NUPACK is used to determine the sticky end melting temperature to find all sequences that fall between the specified melting temperature ± a threshold of 0.5 C to prevent being overly-restrictive (fig. S23 A). The sticky ends used in this work are 8 nucleotides long, contain 4 G or C nucleotides, and have a melting temperature of 15C yielding a candidate pool of 669 sticky end sequences. If one were to choose N=8 total sticky ends from this pool, there would be ~9.54e17 combinations, thus brute-force optimization is not viable. Instead, a GPU-enabled parallelized simulated annealing (40) algorithm is used to search the space where the state corresponds to a subset of N sequences drawn from the pool of 669.

A NUPACK-derived matrix is precomputed by evaluating the ensemble free energies (esf), $\Delta G$, between the candidate sticky end sequences, $s$, and all other candidate sequences, as well as their reverse complements, $\bar{s}$ (fig. S23 B). Ensemble free energies are negatively valued with more negative values corresponding to stronger interactions between two sequences. The diagonal terms, $\Delta G(s_i, \overline{s_i})$, are the on-target bindings, whereas the off-diagonal terms represent undesired interactions. The state is represented as the N selected sequences $S = \{s_1, s_2, \dots, s_N\}$, and the objective function looks to maximize the worst-case thermodynamic margin between each sticky end's on-target interaction and its strongest off-target interaction. This is mathematically defined as:

$$Margin(S) = \min_i[\min\left(\min_{j \neq i} \Delta G(s_i, \overline{s_j}) - \Delta G(s_i, \overline{s_i}), \min_j \Delta G(s_i, s_j) - \Delta G(s_i, \overline{s_i})\right)].$$

This margin-based objective function therefore reports the weakest thermodynamic separation anywhere in the current set of N sequences. Maximizing it favors a set of sticky ends whose worst-off target interactions are well-separated from the on-target interactions. Visual inspection of the esf matrix between the initial versus optimized sets of sticky ends shows sticky ends with larger off-diagonal (i.e., less-interacting) interactions (fig. S23 D).

During the optimization process, the state is iteratively perturbed by randomly swapping sequences in the set $S$. The acceptance of a new state is guided by the Metropolis criteria (49) where early on in the search the optimizer is more likely to hop out of local minima before converging to an optimally-directed set of sticky ends. Here, a geometric cooling schedule is used with a cooling rate of 0.9900 and the initial temperature is set by taking the standard deviation of a random walk for 1000 steps (41) (fig. S24 A). This work uses 4096 parallel instances where each instance runs for 50000 steps (fig. S24 B and C). GPU computation 1) enables one to verify convergence to an objective function value in a single run of the script (fig. S23 C) and 2) is not restricted to an explicit thread count (e.g., a CPU), thus running thousands of parallel instances is feasible. Finally, the resultant sticky ends can be exported alongside their on-target reverse complements to be attached to user-designed overhangs. The sticky end optimization code is provided (47); however, you must have your own personal NUPACK license and its Python module installed to use this portion of the code.

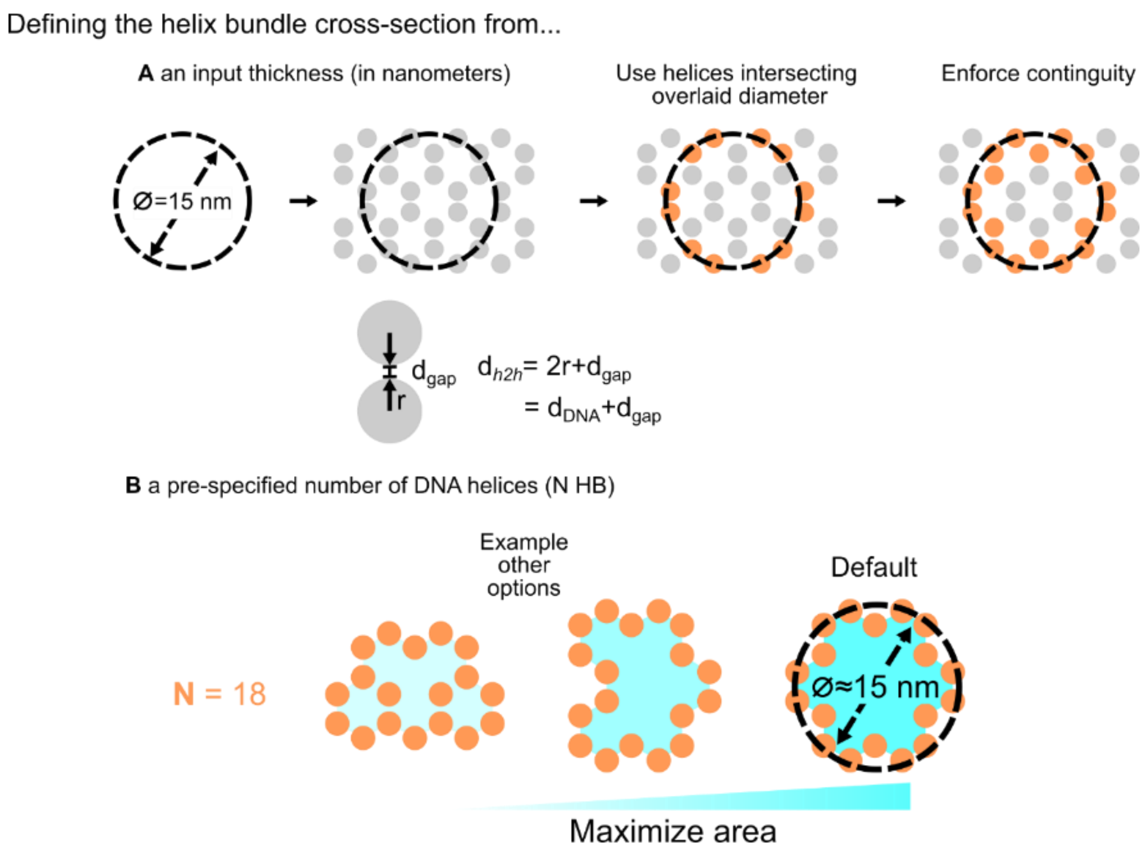


**Fig. S1.** The DNA helices are arranged in a honeycomb grid whose center-to-center spacing is determined by the diameter of B-form DNA ($d_{DNA}$) added to an interhelix gap ($d_{gap}$) distance due to the electrostatic repulsion of the negatively charged backbone. (A) If a thickness is specified, a number of helices are placed into a contiguous ring, otherwise (B) if a predefined number of helices is specified then they are arranged into a contiguous ring of maximal cross-sectional area.

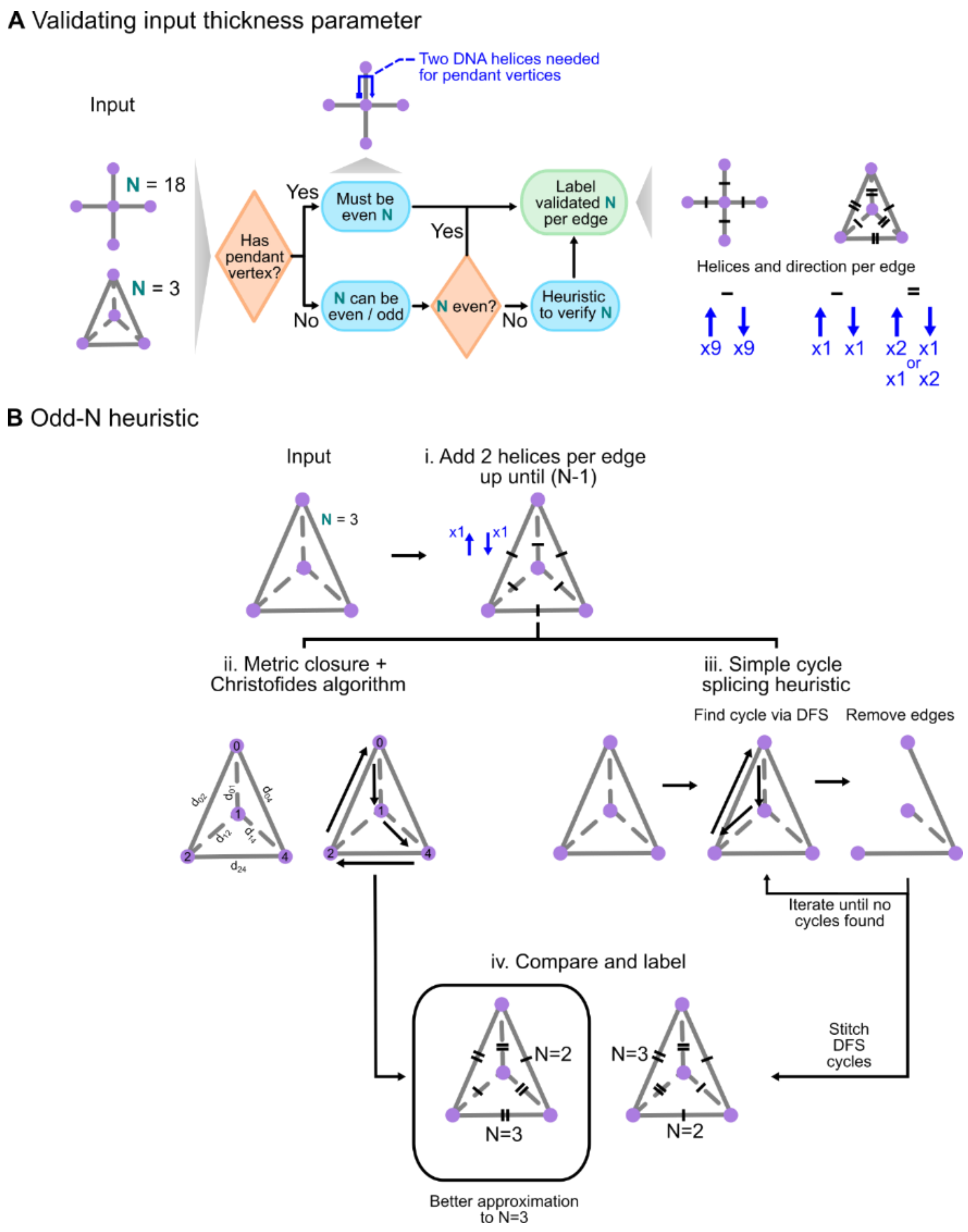


**Fig. S2.** (A) The input graph topology and specified N DNA helices per bundle must be validated. If N is even, then a continuous scaffold routing can be guaranteed (fig. S8) regardless of topology by assigned N / 2 opposite-running helices per edge. (B, i.) If N is odd, then first each edge gets assigned 2 helices (opposing-running) until N-1 is reached. (ii) A metric closure is defined, and Christofides algorithm is compared with a (iii) simple depth first search of cycles heuristic for determining an Eulerian tour. (iv) These traversals are compared to see which approximates the specified N more closely across all edges.

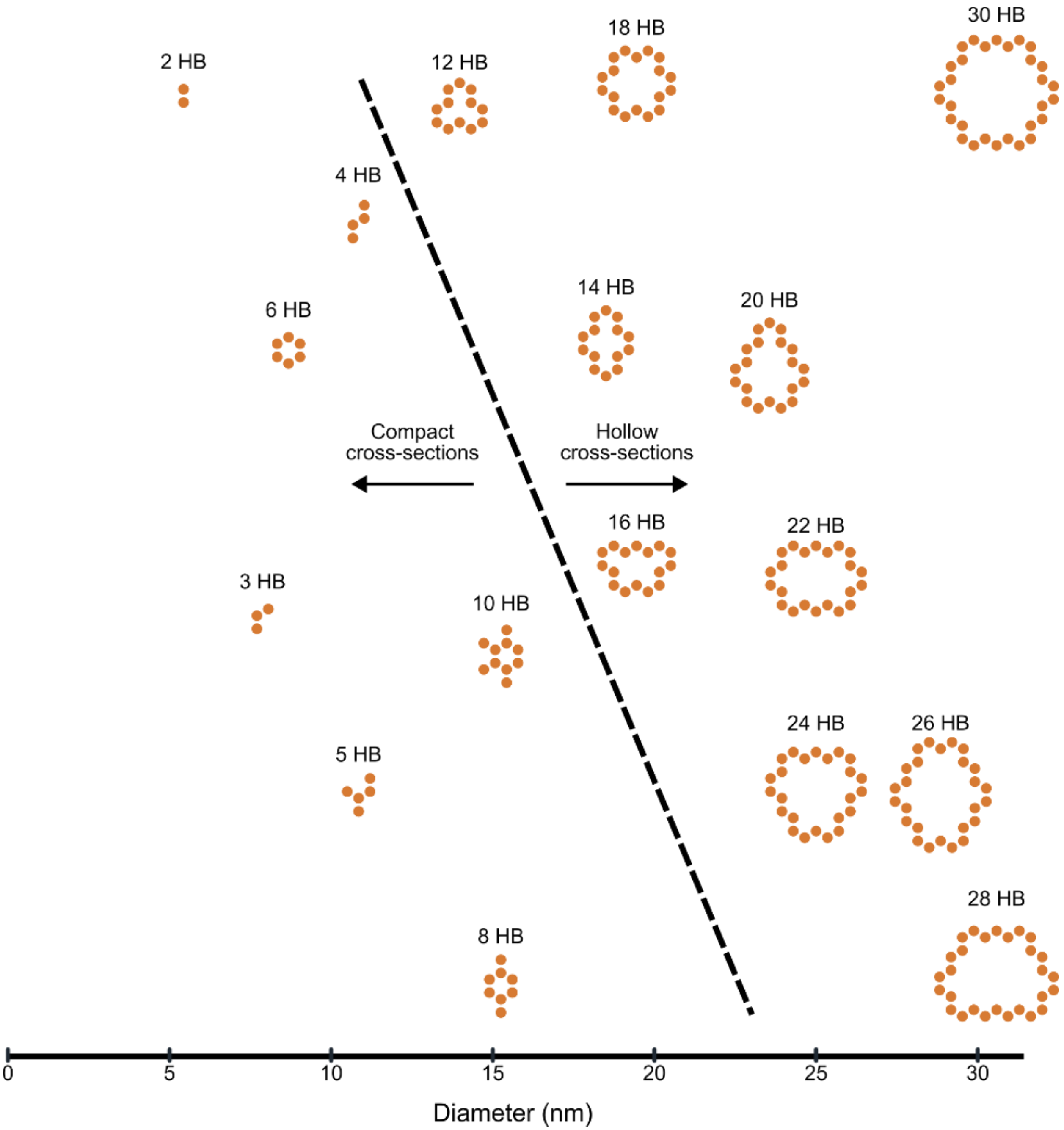


**Fig. S3.** Helix cross-sections for input target diameters (fig. S1) ranging from 0 to 30 nanometers. Whenever at least 12 HB are set, then cross-sections contain a “hollow” cavity within the honeycomb lattice.

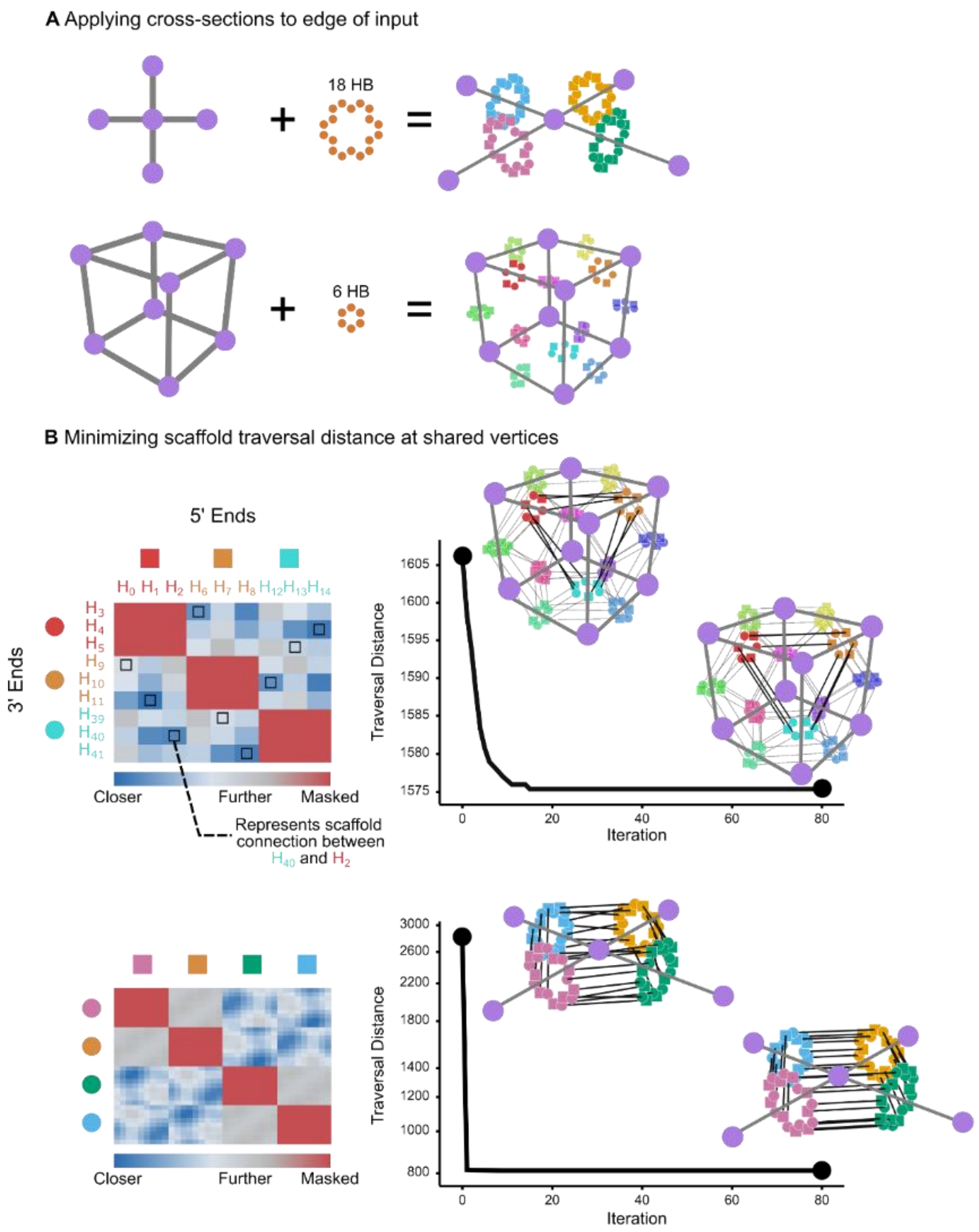


**Fig. S4.** (A) Cross-sections are first aligned along each edge of the input wireframe model. The individual 3’ (circle) and 5’ (square) helices have 3D positions. (B) The cross-sections are then rotated about the central edge axis in discrete settings leveraging the 60-degree rotational symmetry of the honeycomb grid. At each rotation, a pairwise distance matrix is evaluated at each vertex with at least two edges to find minimal-traversal distance pairings to route the scaffold through.

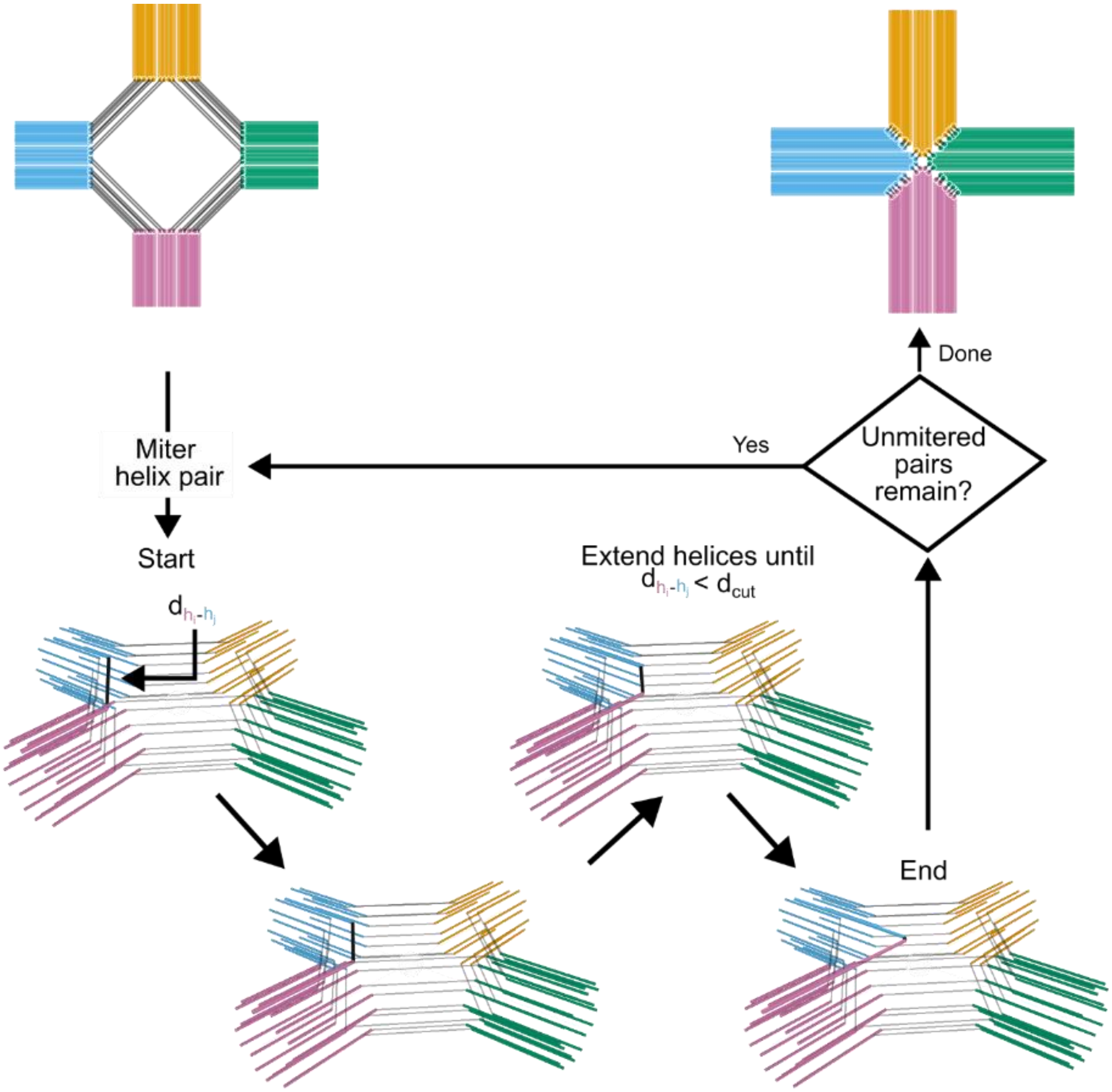


**Fig. S5.** Mitering is applied to the helix pair found by the vertex matching (fig. S4). For each pair, the mitering algorithm extends the individual helices along their respective central axes one nucleobase (0.34 nanometers) at a time until the helix-to-helix distance is less than a cut-off distance ($d_{cut}$ = 2.0, fig. S7).

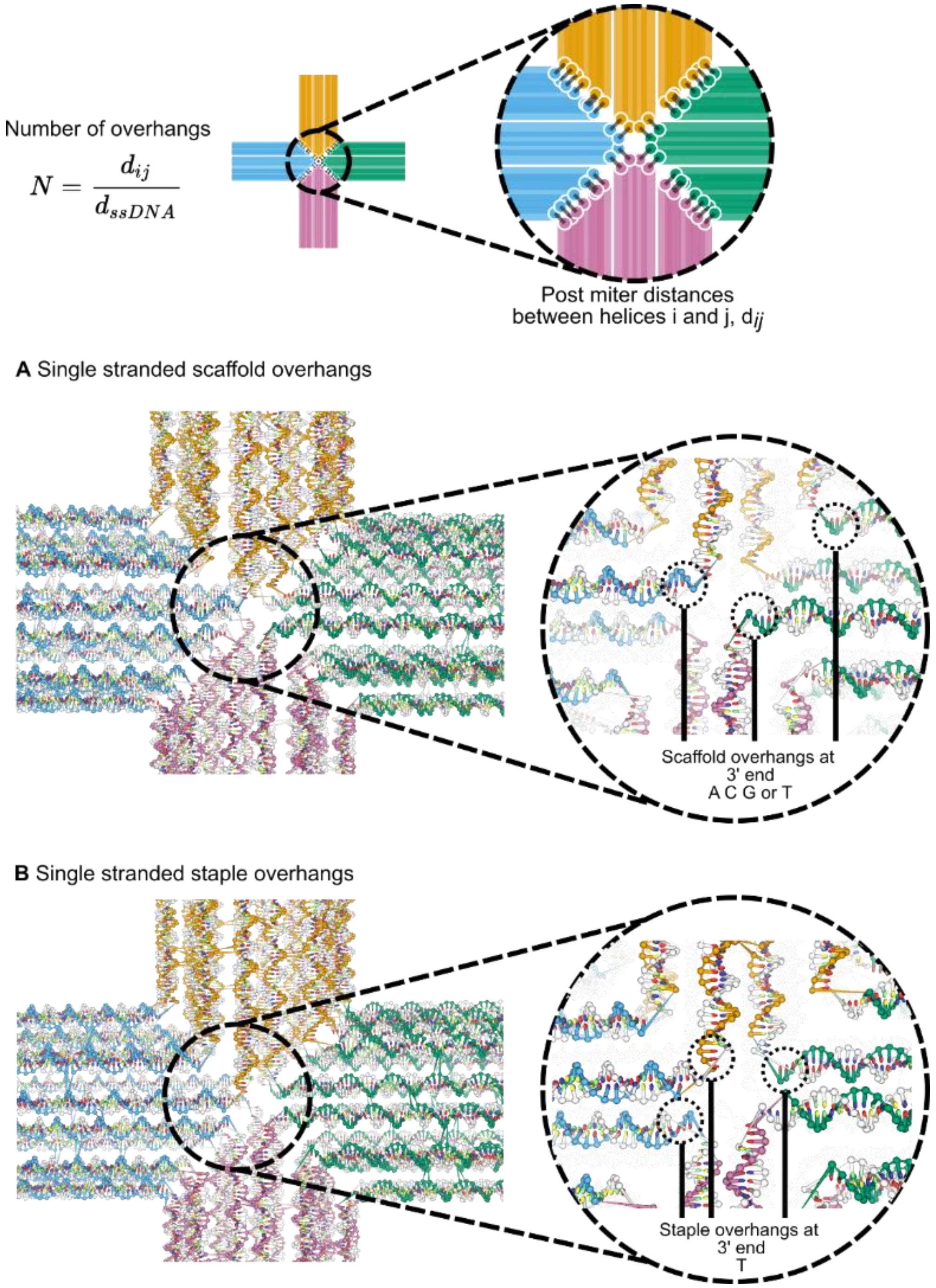


**Fig. S6.** Single stranded overhangs are added at the 3' end of the scaffold and staple connections to minimize backbone torsion. The number of overhangs is equal to the final found distance (approximately $d_{cut}$ = 2.0) divided by a presumed nucleotide-to-nucleotide spacing of single stranded DNA ($d_{ssDNA}$ = 0.60, fig. S7). (A) Scaffold single stranded overhang regions are scaffold-sequence dependent, and maybe any of A C G or T. (B) Staple single stranded overhangs are always T (by default).

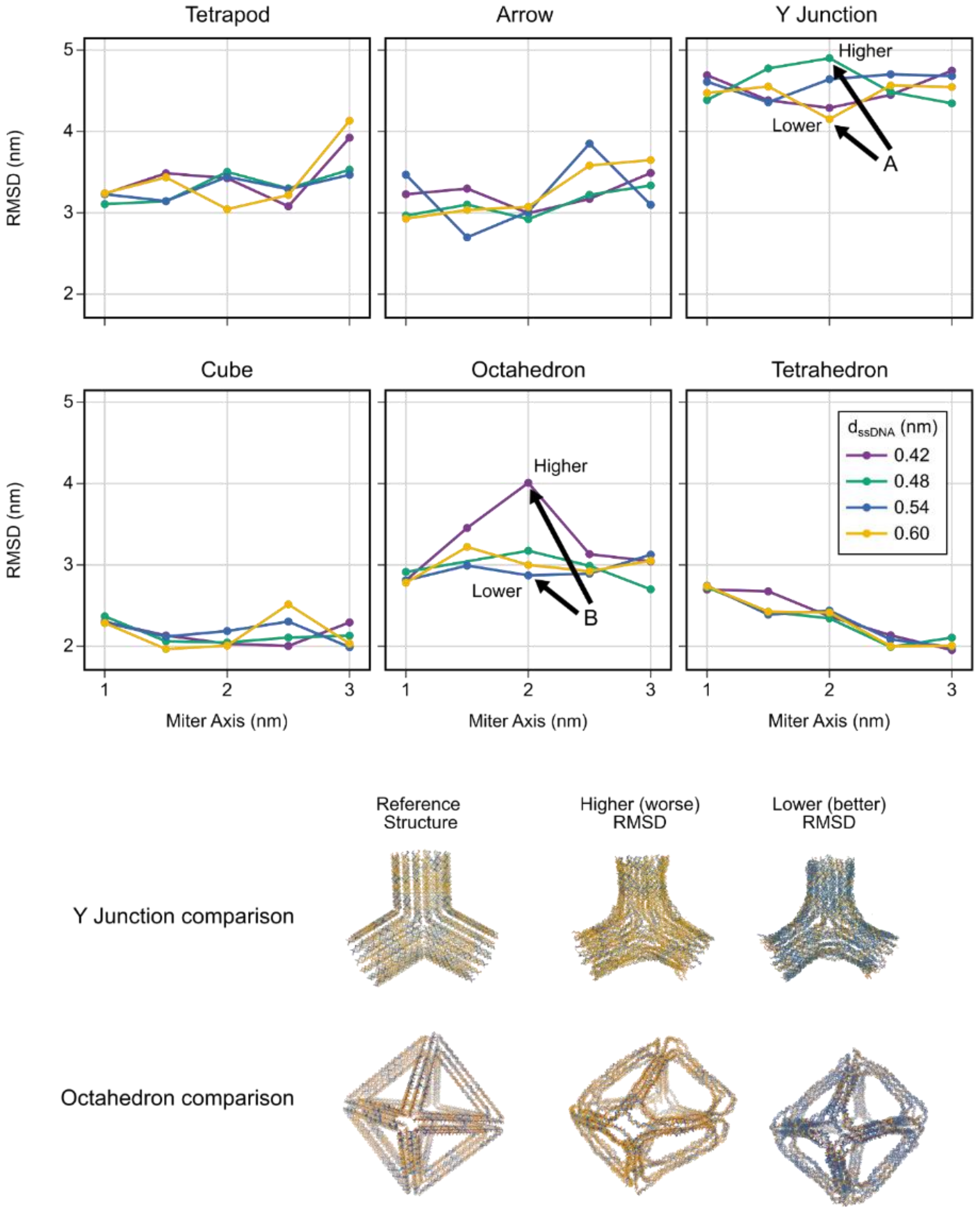


**Fig. S7.** Default values for the $d_{cut}$ and $d_{ssDNA}$ (fig. S5 and S6) parameters were determined through oxDNA simulations across combinations of their settings. The Kabsch-aligned RMSD between the ideal, reference structure and the centroid configuration post-oxDNA simulation is used to inform the selection of the parameters.

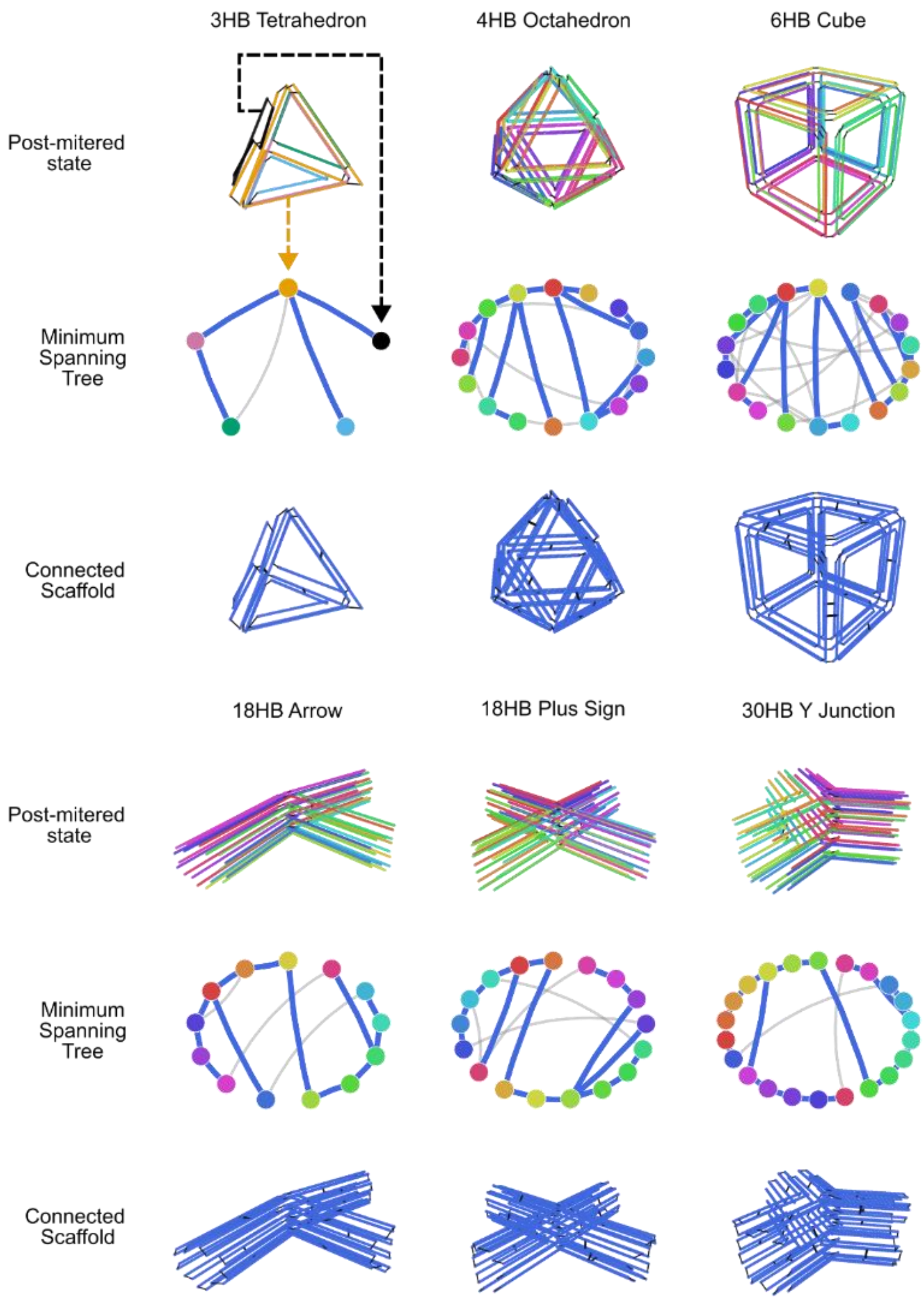


**Fig. S8.** A minimum spanning tree algorithm is used to add internal double stranded scaffold crossovers (22) to connect a single scaffold routing. Nodes are the individual cycles found by the rotation algorithm (fig. S4) and the edges are placed between cycles where an internal double stranded crossover can be placed, thus connecting those two cycles. The total number of potential crossovers is found and applied to each edge, and the minimum spanning tree (blue thicker edges) is used to select which ones are placed to connect a single, continuous scaffold.

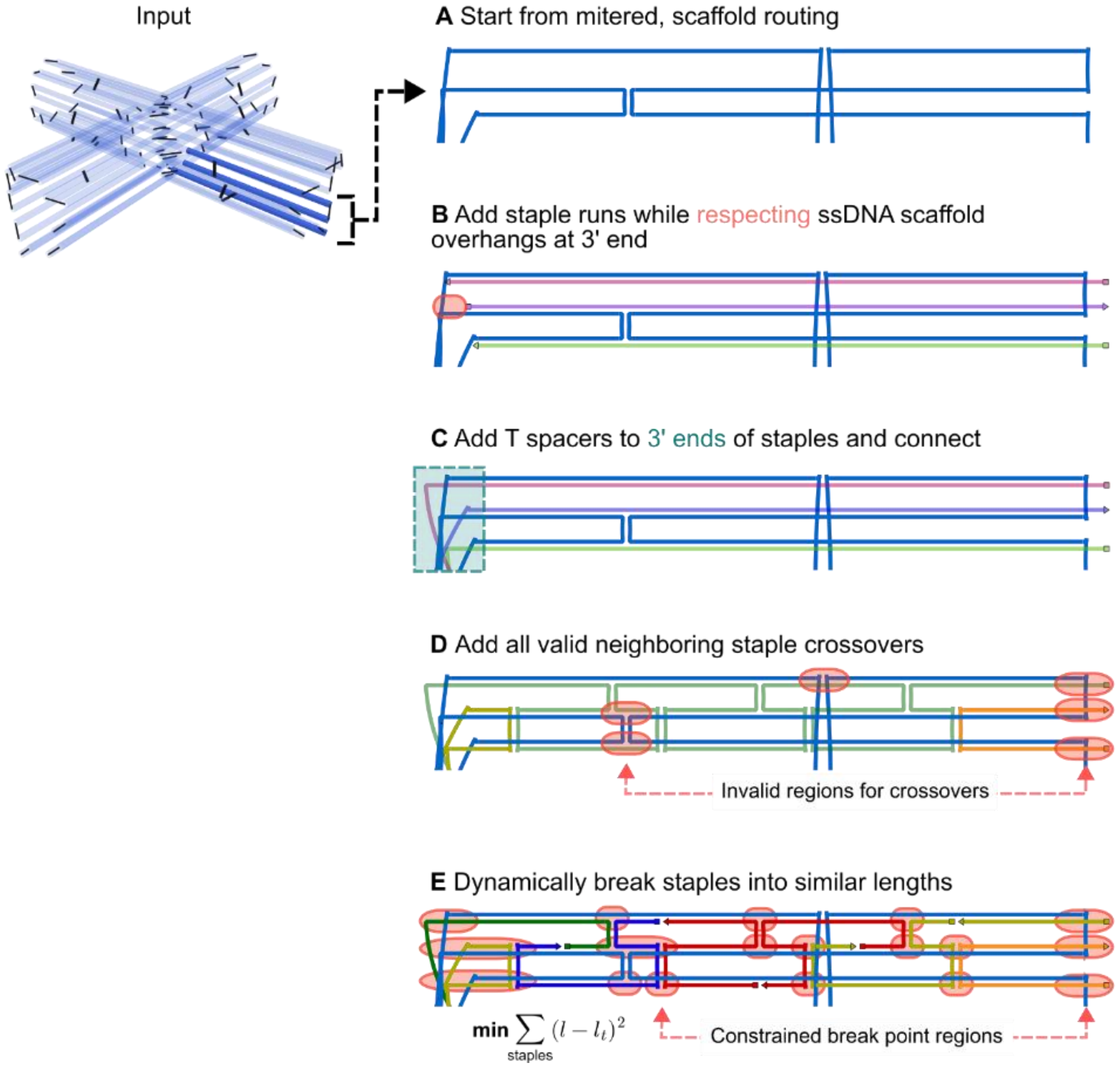


**Fig. S9.** (A) The stapling algorithm starts by looking at the scaffold-routed design (fig. S8) and (B) staple runs are added to pair with each staple nucleotide while ignoring the single stranded regions (fig. S6). Also, single stranded overhangs are added at any terminal ends to serve as p`oly-T brushes. (C) Then, single stranded thymine spacers are added to the 3’ ends of the staples (fig. S6 B), and all staple ends are connected based on the final rotated state (fig. S4 B). (D) All staple crossovers based on the honeycomb-style grid that are not near scaffold crossovers are added (near being within 3 nucleotides, by default). (E) Finally, long staples are dynamically spliced into similar length segments (fig. S10) while respecting various constrained regions that could lead to kinetic traps during physical folding.

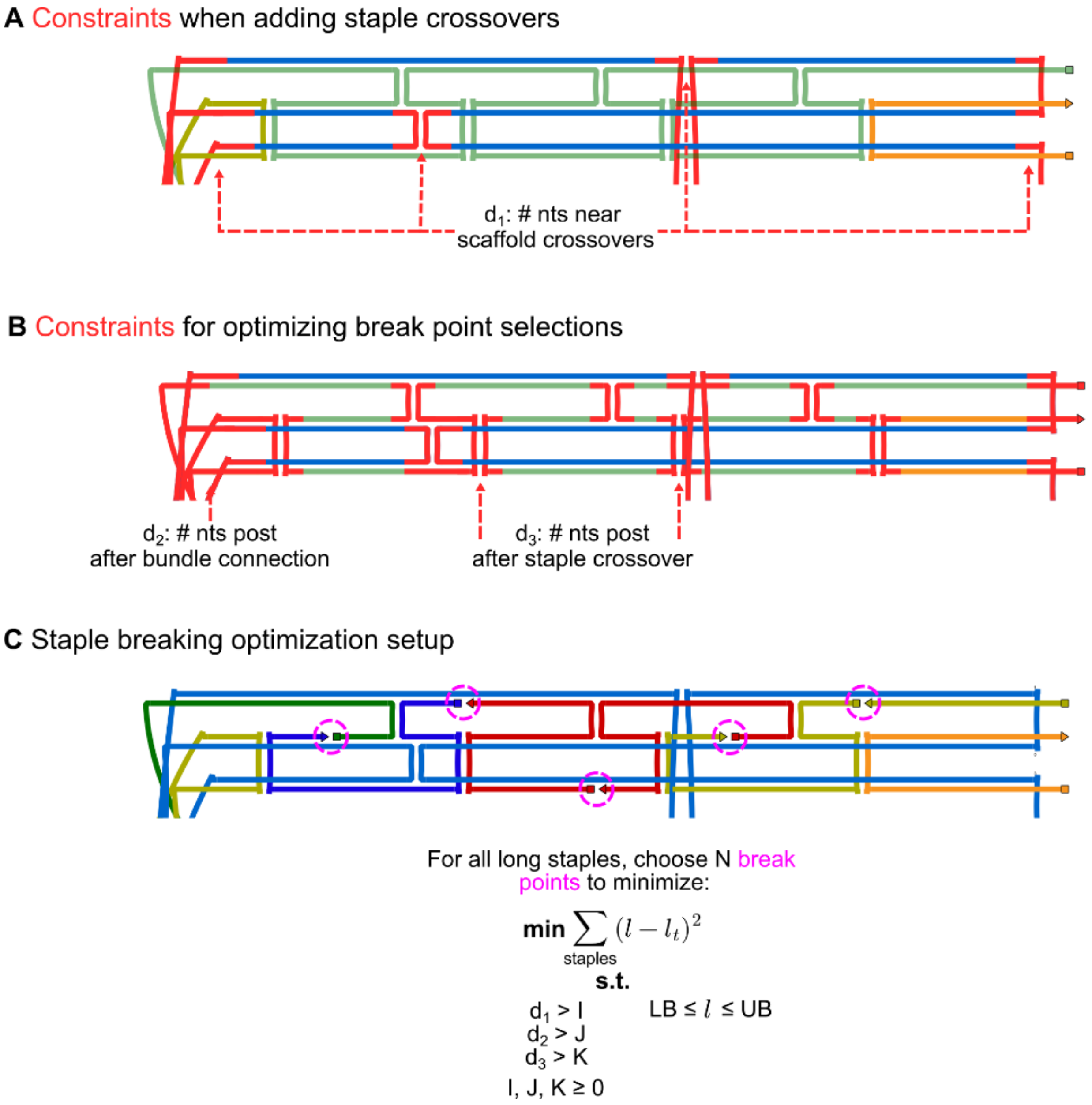


**Fig. S10.** The automated staple breaking algorithm starts by identifying the constrained nucleotides (nts) that are within (A) $d_1$ nts of scaffold crossovers, (B) $d_2$ nts of a previous bundle connection, or $d_3$ nts after a staple crossover. (C) Then, for all staples that are longer than an upper bound (UB), N break points are chosen that minimize mean square error towards a target staple length subject to the constrained nucleotides above and ensuring all staples are bounded between a lower bound (LB) and the UB. By default (fig. S11), $d_1 = 3$, $d_2 = 5$, $d_3 = 3$, LB = 20, and UB = 60 (all in units of nts).

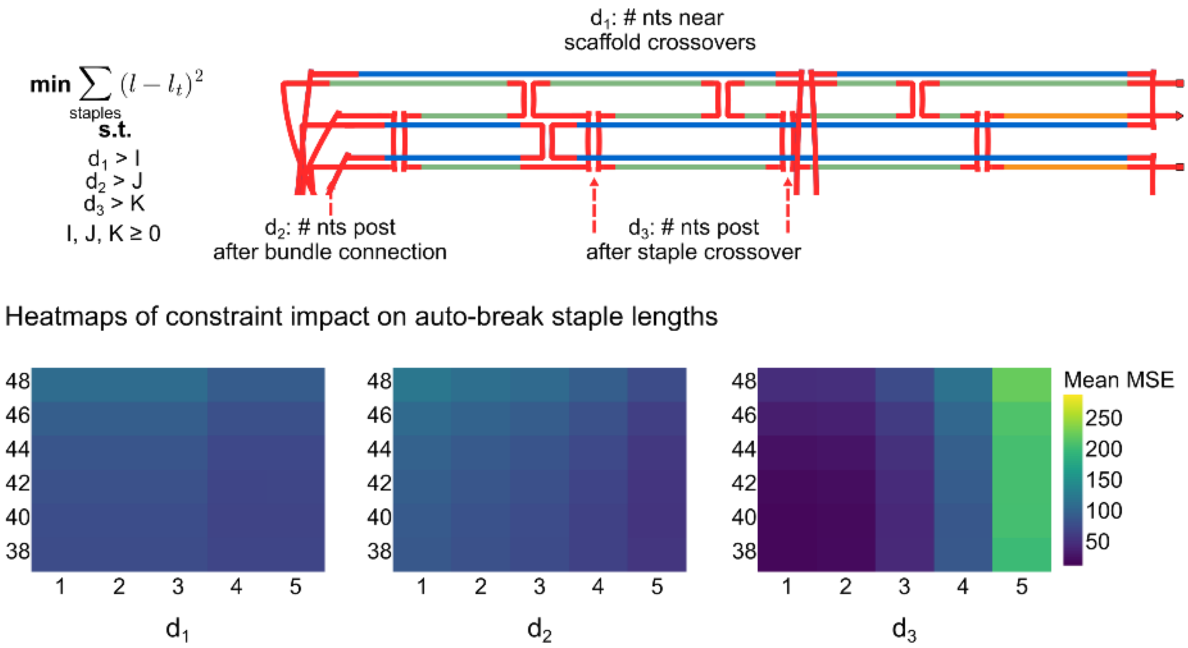


**Fig. S11.** Analysis of 95,000 designs generated by various topology, edge length, and cross-sections (some shown in fig. S12 and S13). The staple constraints ($d_1$, $d_2$, $d_3$ in fig. S10 C) alongside the target staple length were also swept through. The mean MSE is the average per-staple objective function valuation, and as $d_3$ is set larger, the MSE greatly increases as there are few break points for selection. Comparatively, $d_1$ and $d_2$ can be set to larger values (e.g., 5 nucleotides of binding between the staple and scaffold) without impacting the MSE greatly.

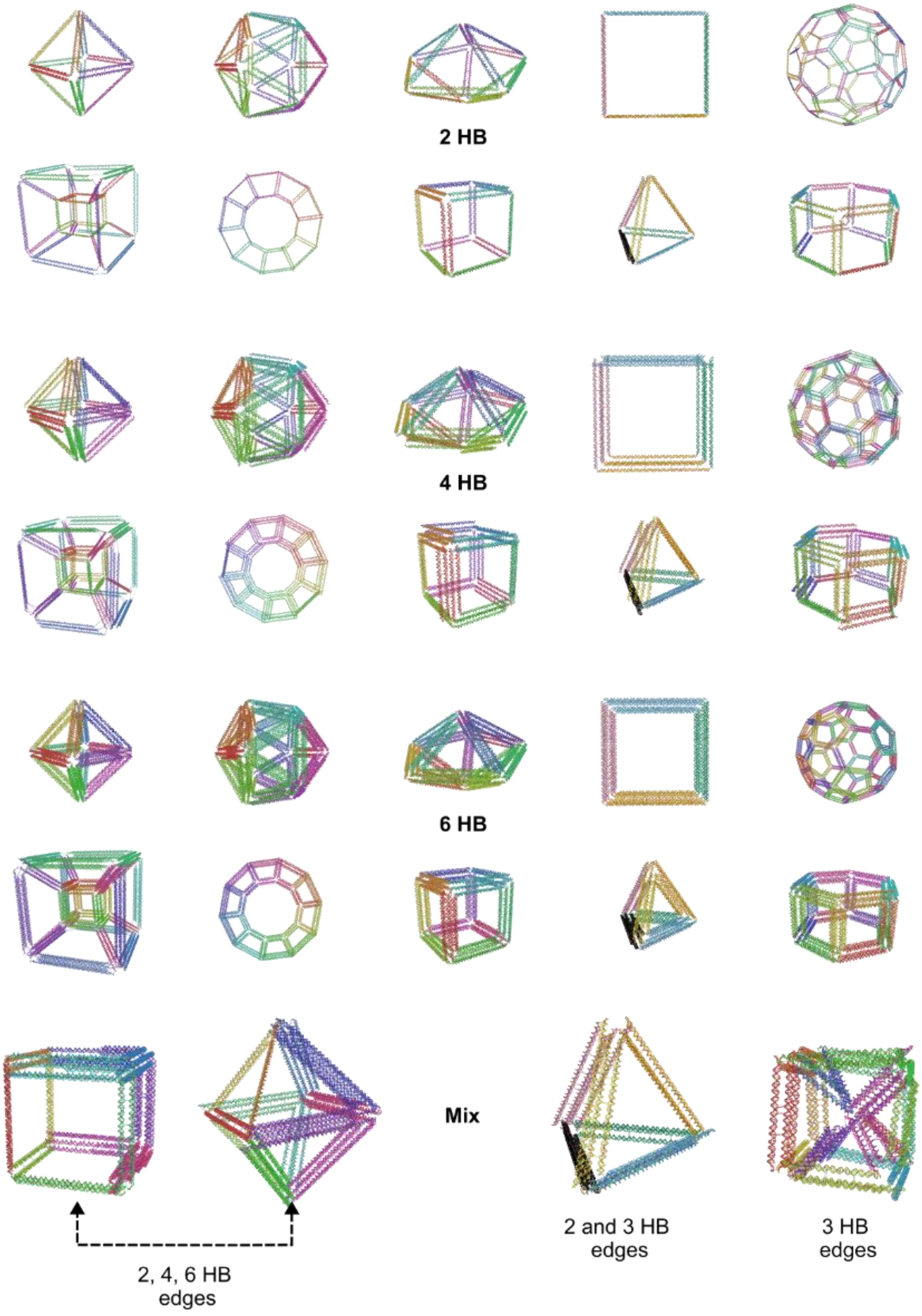


**Fig. S12.** A range of traditional wireframe DNA origami nanostructures with constant 2, 4, or 6 HB edges or variable, mixed-edge assignments are shown.

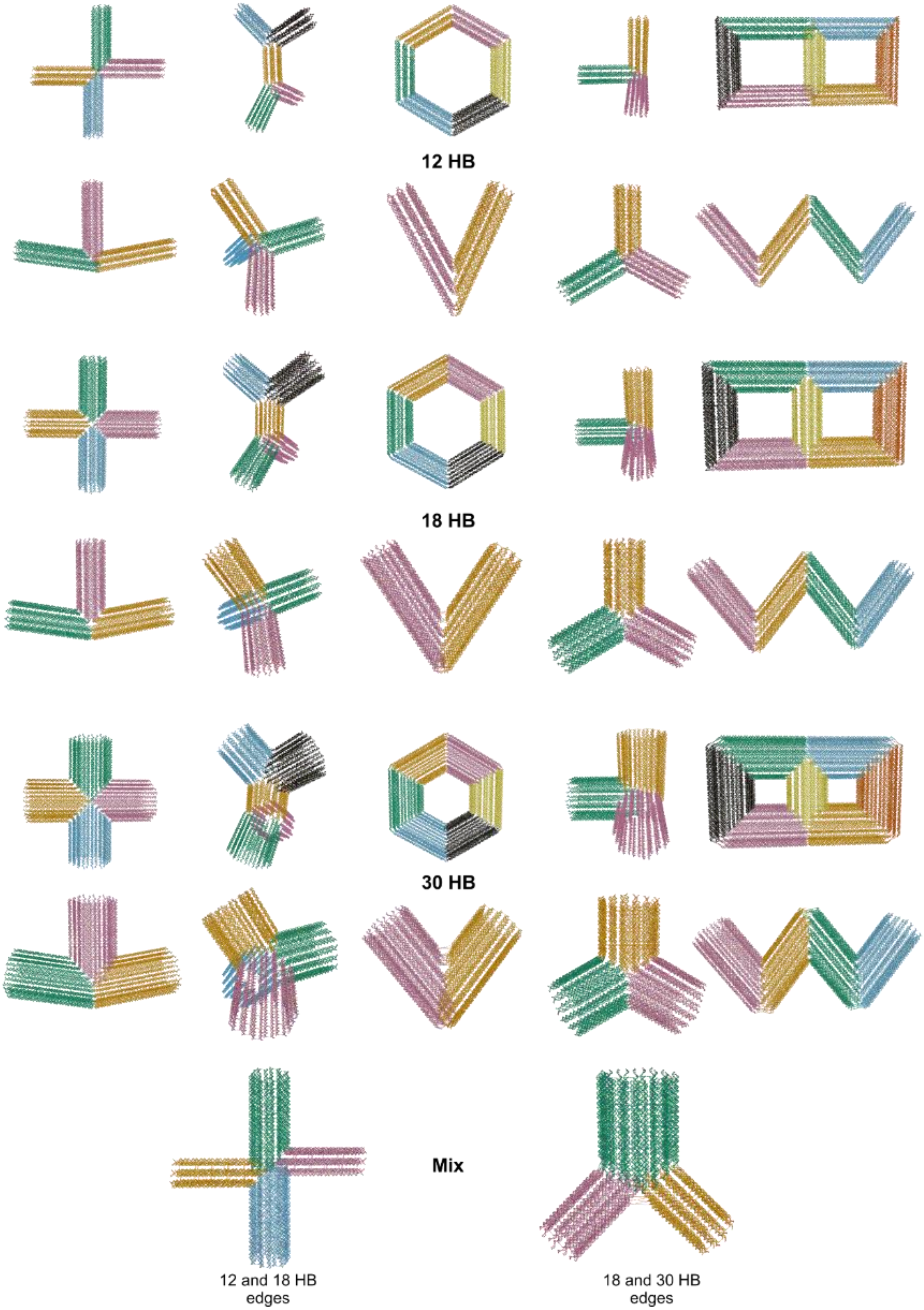


**Fig. S13.** A range of hollowframe DNA origami nanostructures with constant 12, 18, or 30 HB edges or variable, mixed-edge assignments are shown.

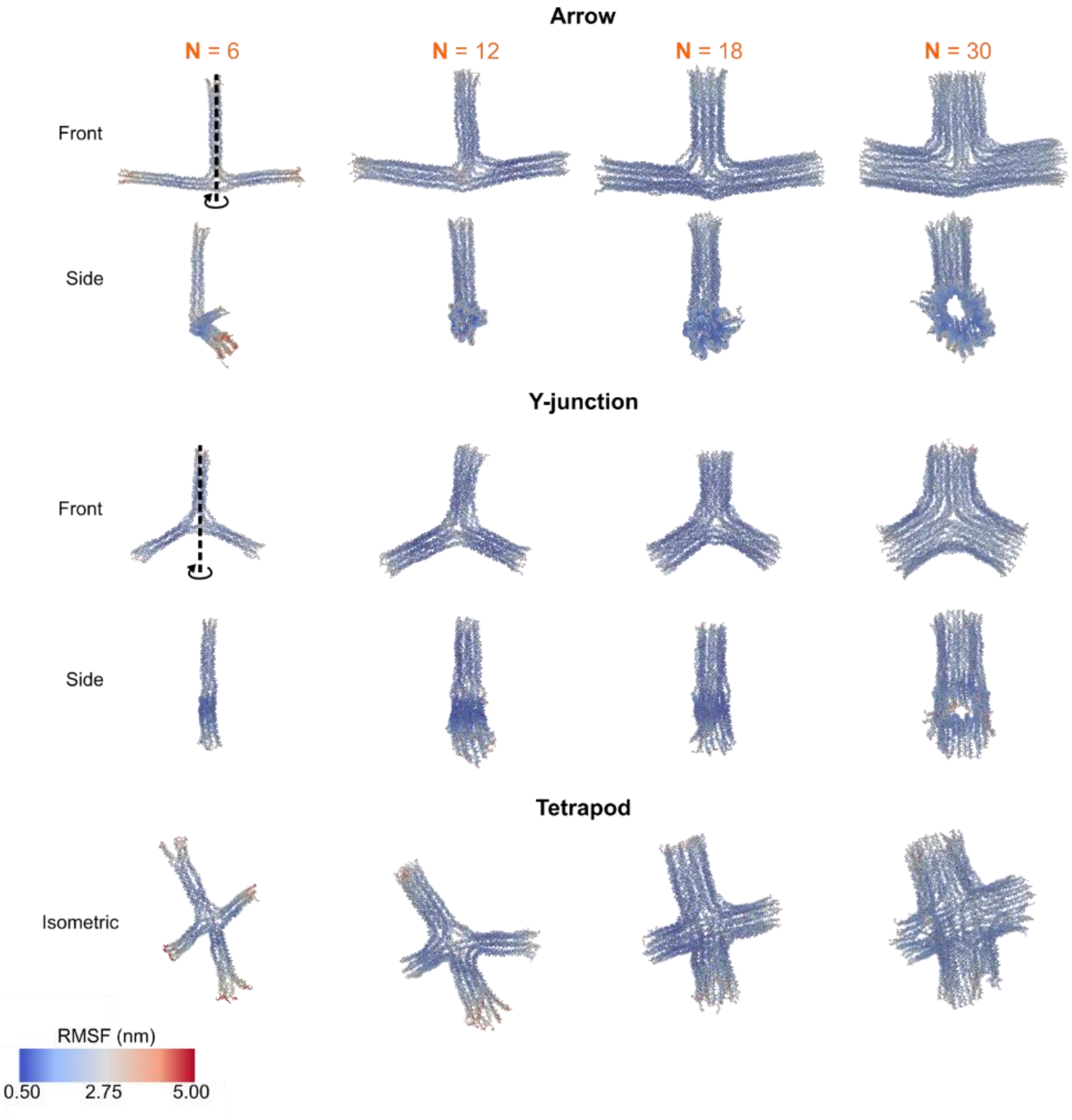


**Fig. S14.** oxDNA simulations show nanostructure flexibility with increasing the number of DNA helices per edge, reducing out of plane (side) fluctuations. All simulations run for 5e7 timesteps using a standard minimize-relax-simulate procedure (see Materials and Methods).

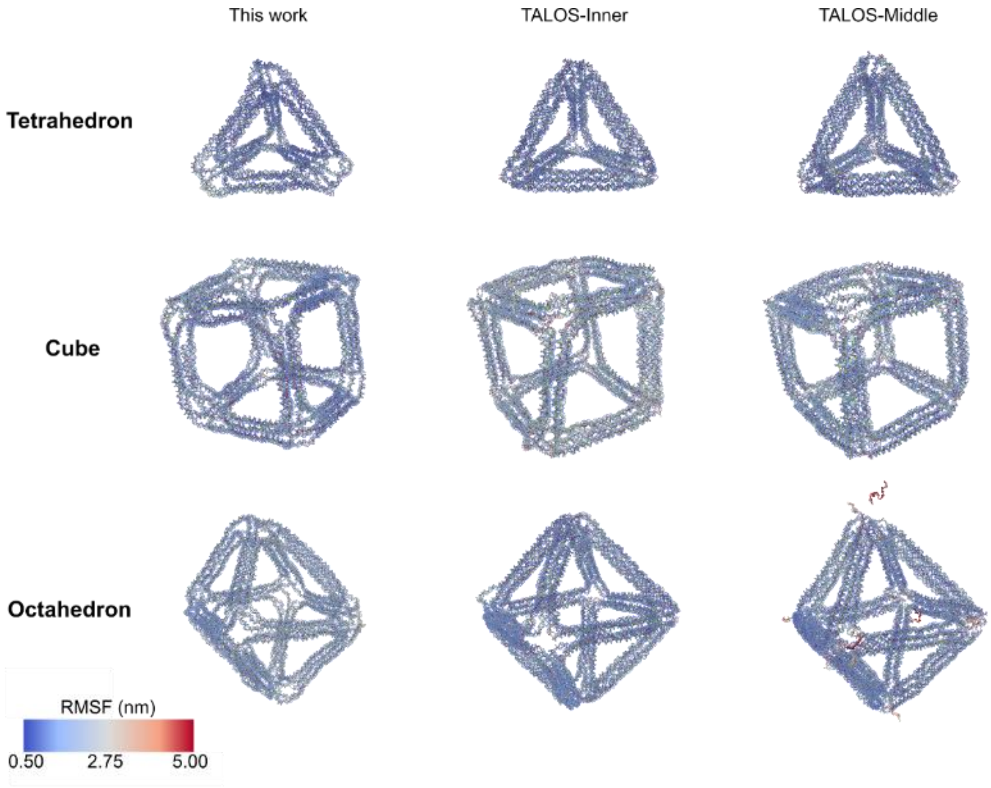


**Fig. S15.** oxDNA simulations comparing 6HB traditional wireframe designs produced by this work with TALOS (23). All simulations run for 5e7 timesteps using a standard minimize-relax-simulate procedure (see Materials and Methods).

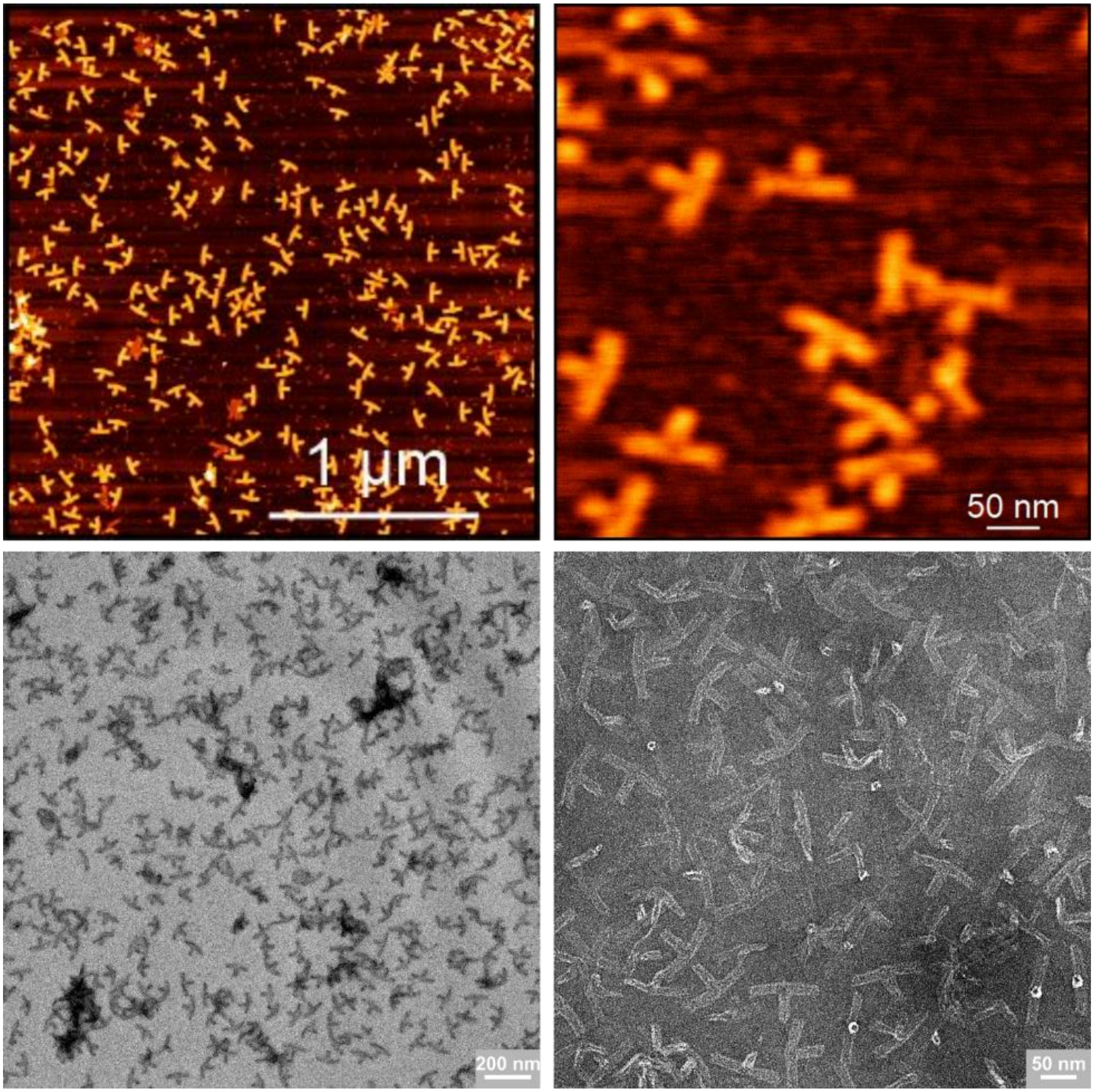


**Fig. S16.** Extended AFM and negative stain TEM images of 18 HB arrow. Scale bars as labelled.

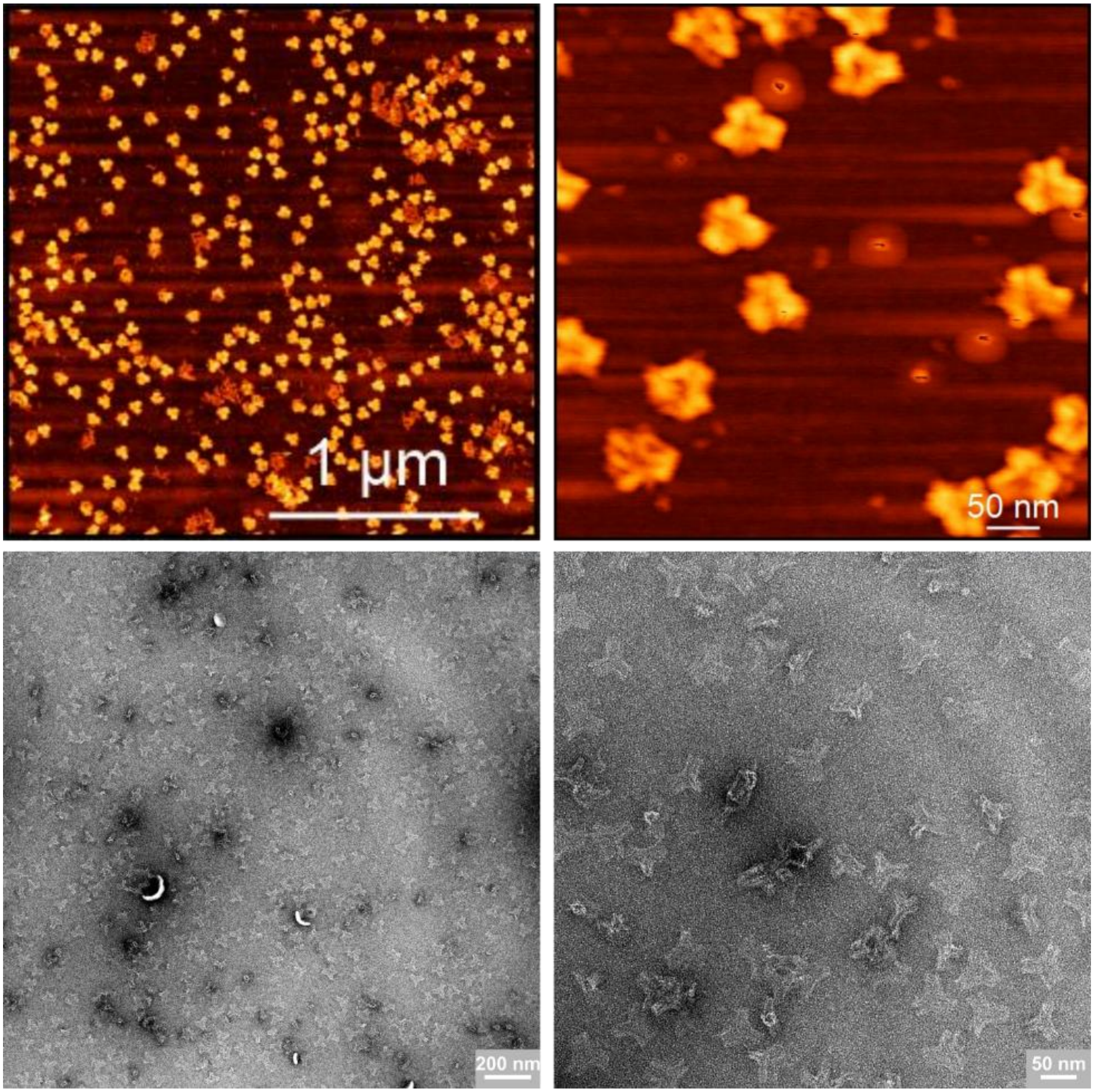


**Fig. S17.** Extended AFM and negative stain TEM images of 30 HB Y-junction. Scale bars as labelled.

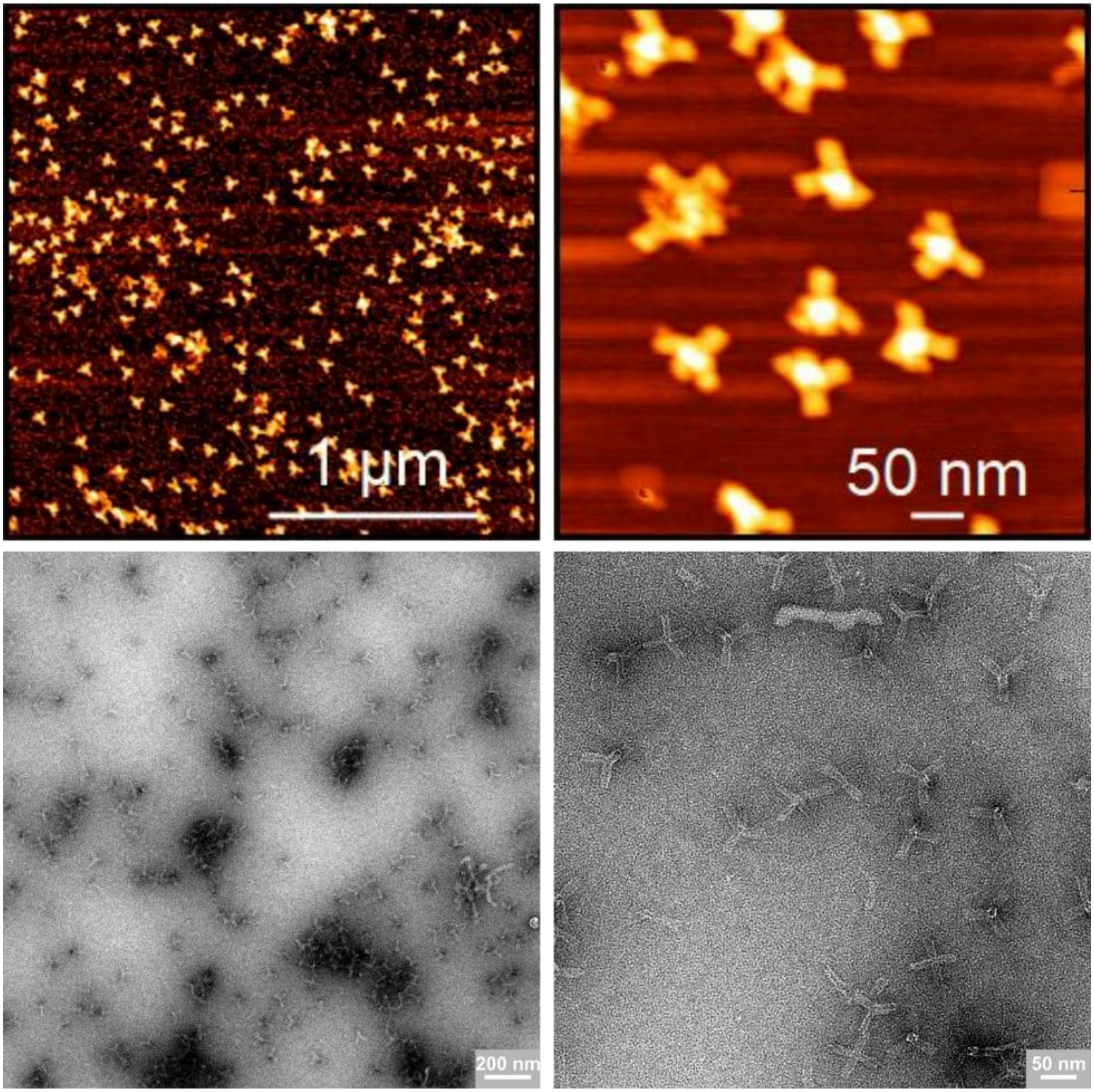


**Fig. S18.** Extended AFM and negative stain TEM images of 18 HB tetrapod. Scale bars as labelled.

**A** 18 HB Arrow Cryo-TEM class averages 105 kx

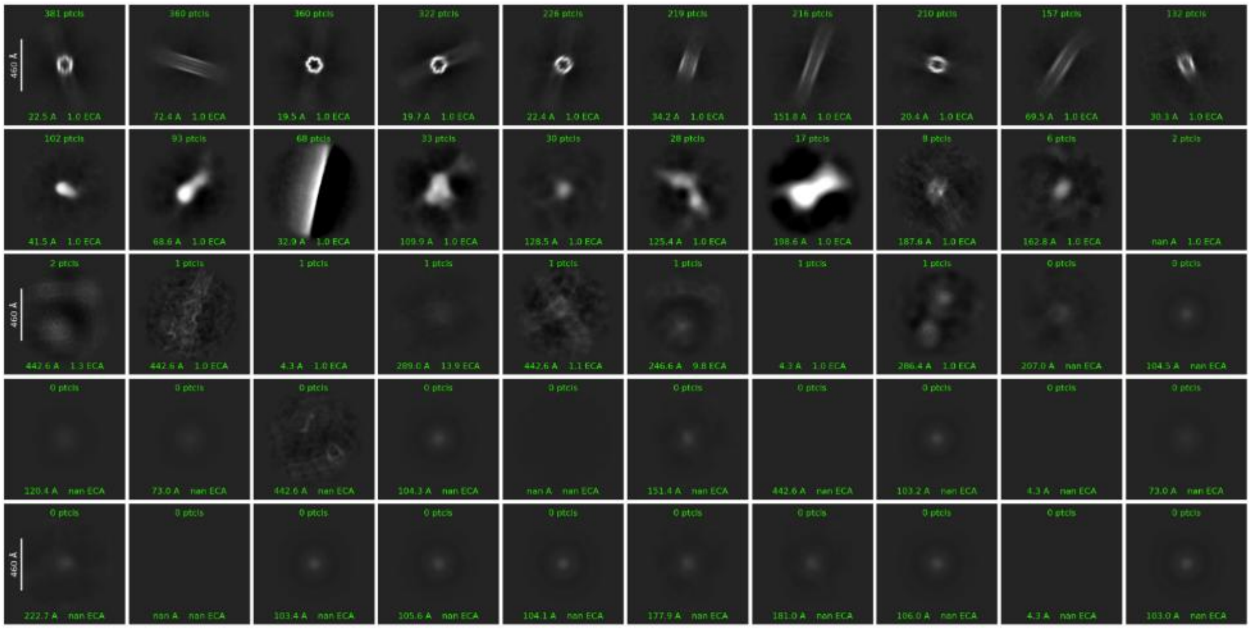

**B** 18 HB Arrow Cryo-TEM class averages 130 kx

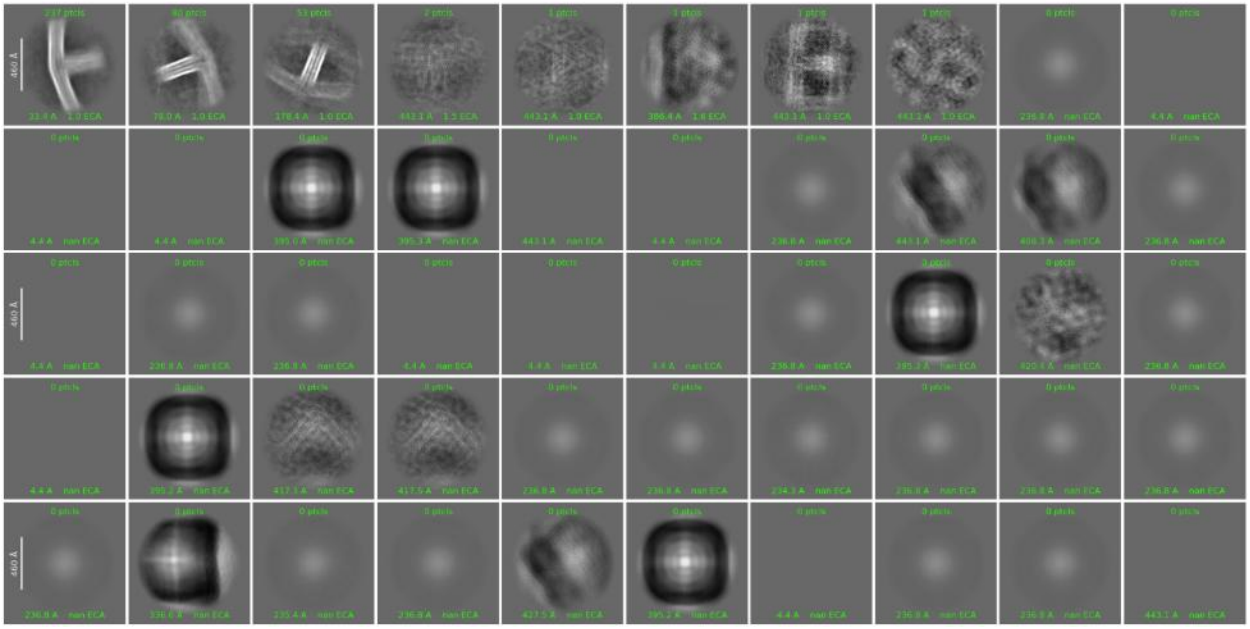

**Fig. S19.** Full Cryo-TEM 2D class averages of the 18 HB arrow at both (A) 105 kx (3,389 particles) and (B) 130 kx (517 particles) magnification.

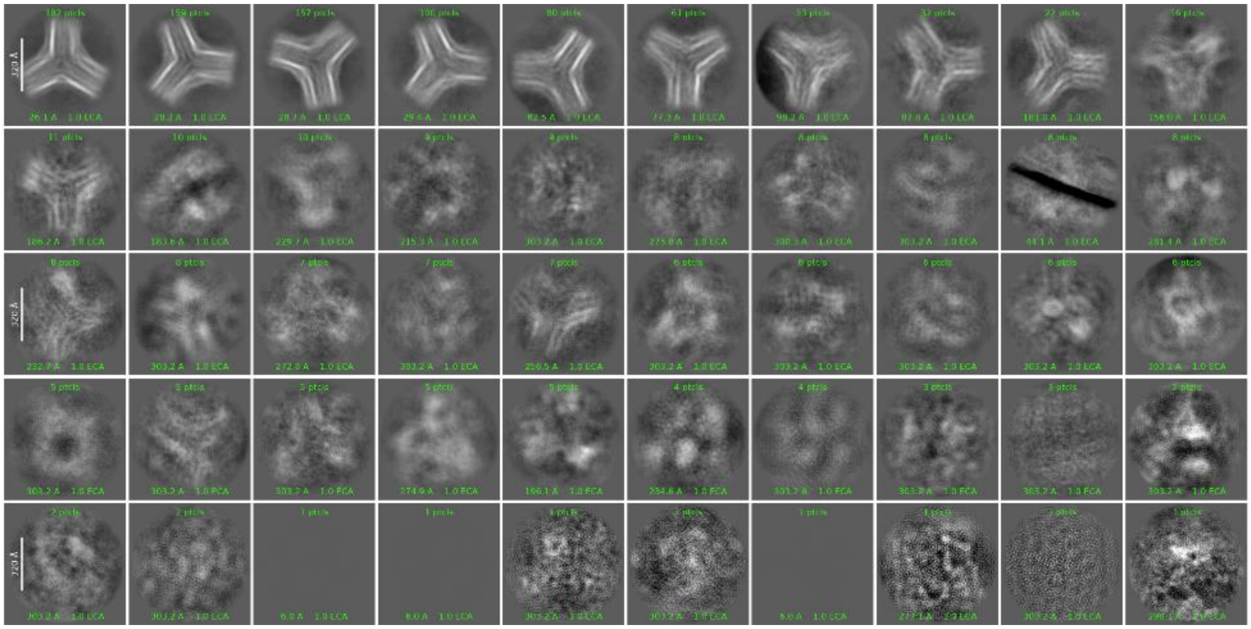

**Fig. S20.** Full Cryo-TEM 2D class averages of the 30 HB Y-junction at 105 kx (1,048 particles) magnification.

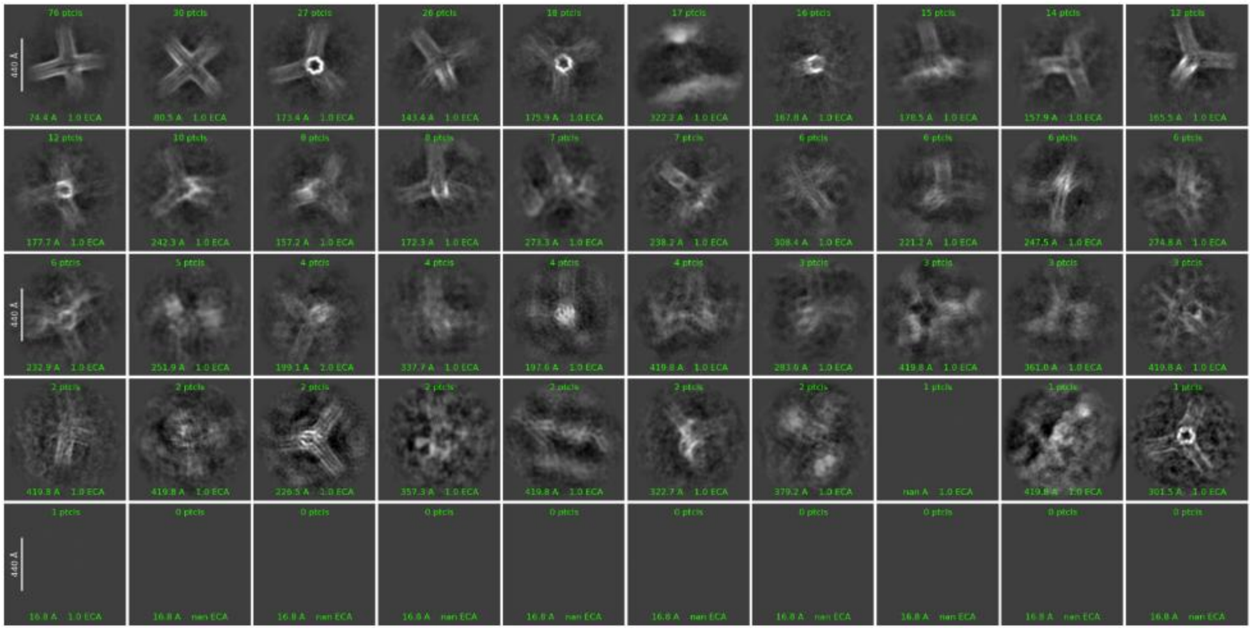

**Fig. S21.** Full Cryo-TEM 2D class averages of the 18 HB tetrapod at 105 kx (215 particles) magnification.

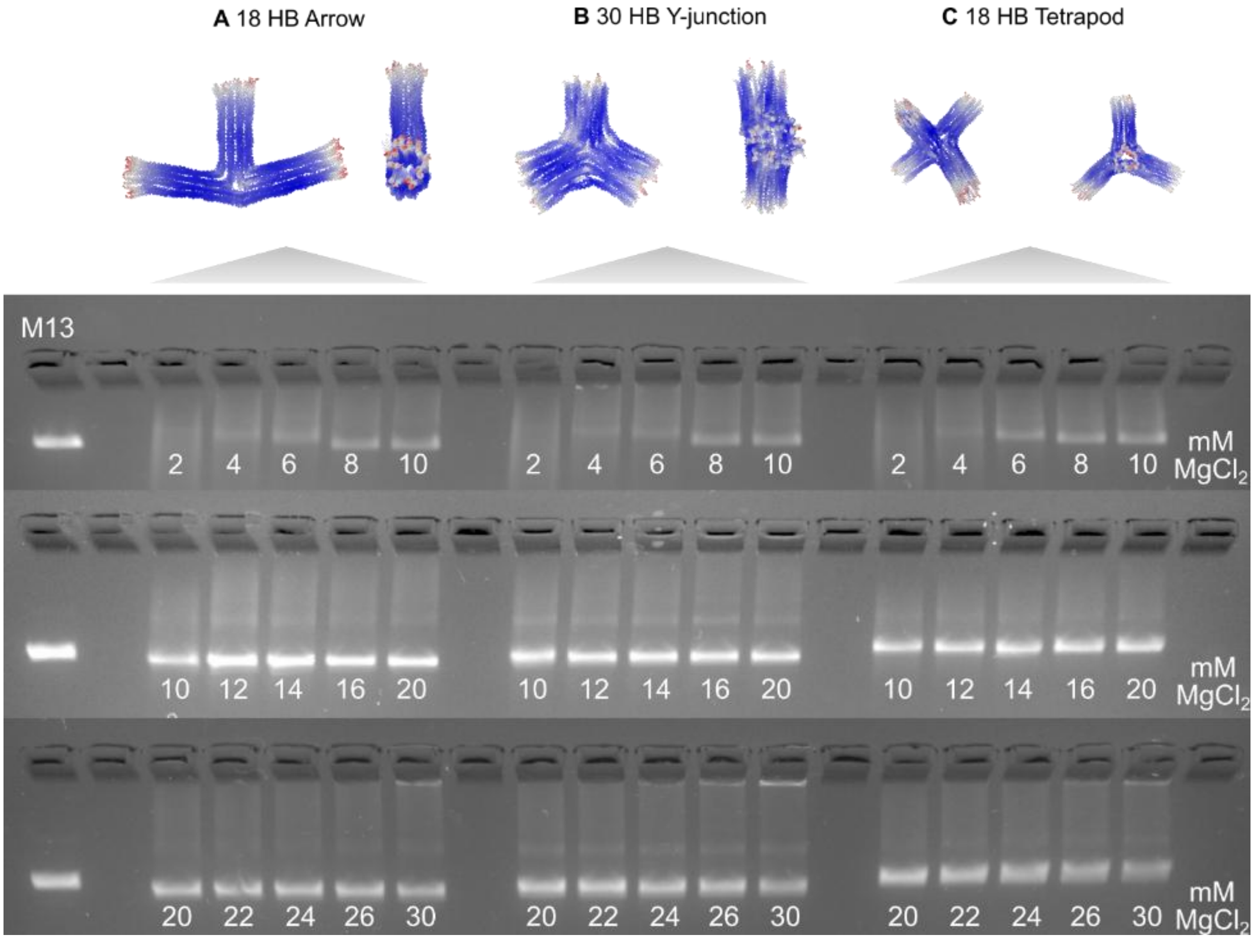


**Fig. S22.** Agarose gel electrophoresis of magnesium chloride (MgCl2) concentrations ranging from 2 to 30 mM for the (A) 18 HB arrow (B) 30 HB Y-junction and (C) 18 HB tetrapod.

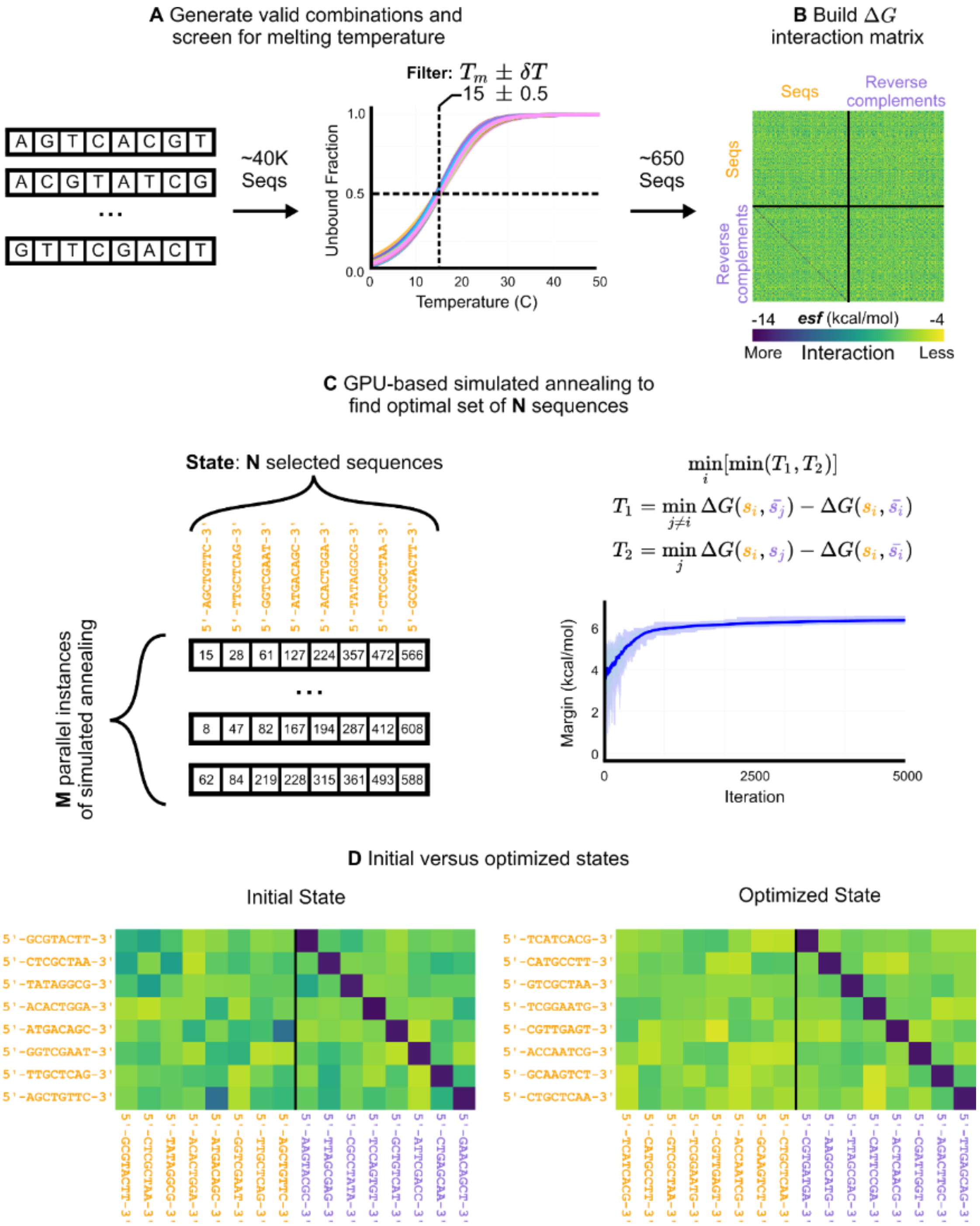


**Fig. S23.** Sticky ends are (A) exhaustively populated and screened based on melting temperature with their respective reverse complement. (B) A pairwise matrix is setup between the sequences and themselves as well as the reverse complements using the ensemble free energy function (39) where more negative values are higher interacting sticky ends. (C) M parallel independent randomized instances of simulated annealing are placed onto GPU to minimize the thermodynamic margin (see Materials & Methods). (D) An example initial versus optimized state is shown, where the darker (higher interacting) regions are removed (excluding the on-target diagonal between the sequences and their reverse complements).

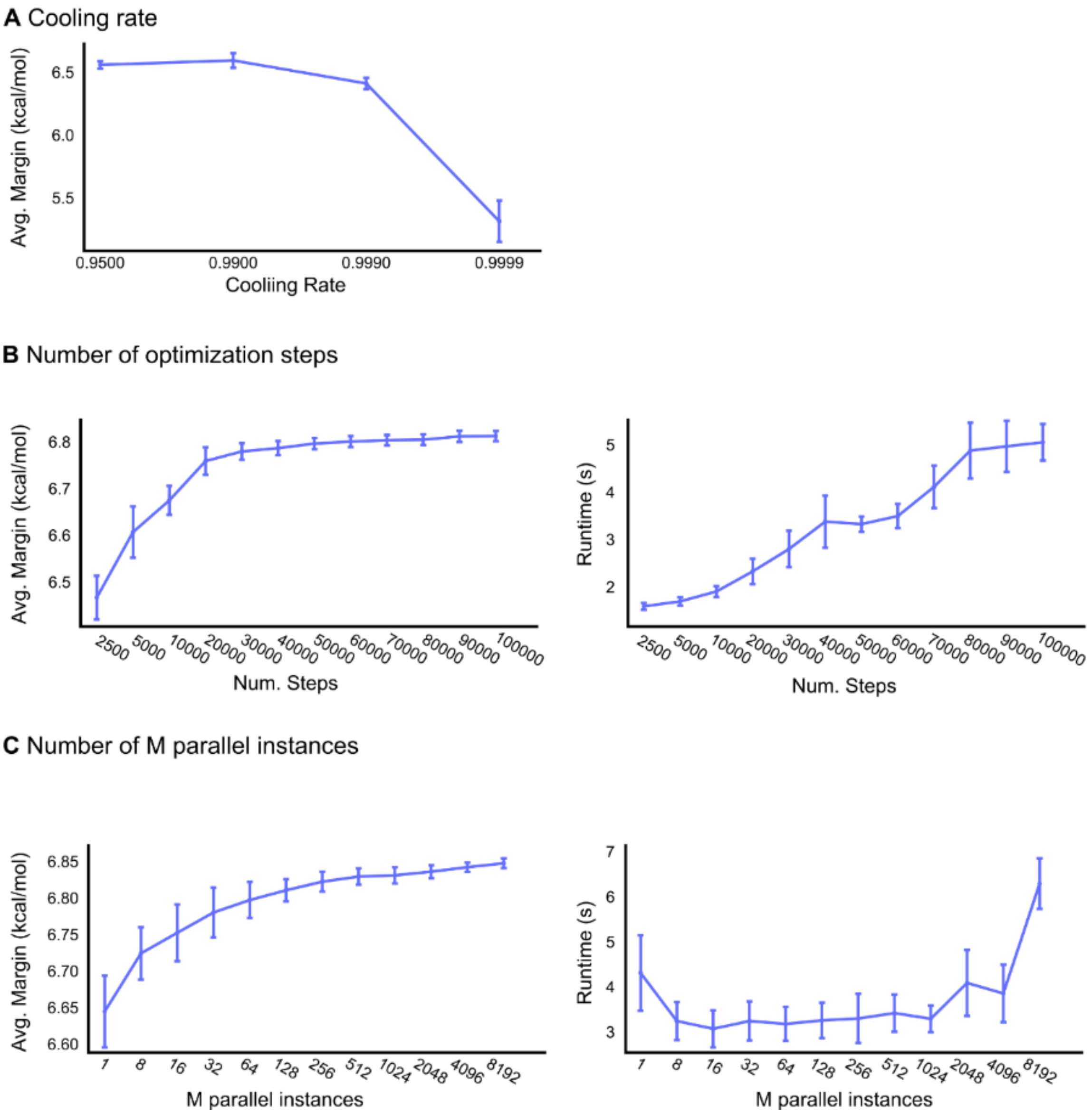


**Fig. S24.** Ablation study of optimizer hyperparameters for simulated annealing (fig. S23). (A) Geometric cooling rate has largest impact on the average margin found, whereas the number of (B) optimization steps and (C) parallel instances establish convergence towards a solution. Error bars show one standard deviation from N=10 random seeds per test case. Studies run on Ubuntu 24.04 using an RTX 4070 Super.

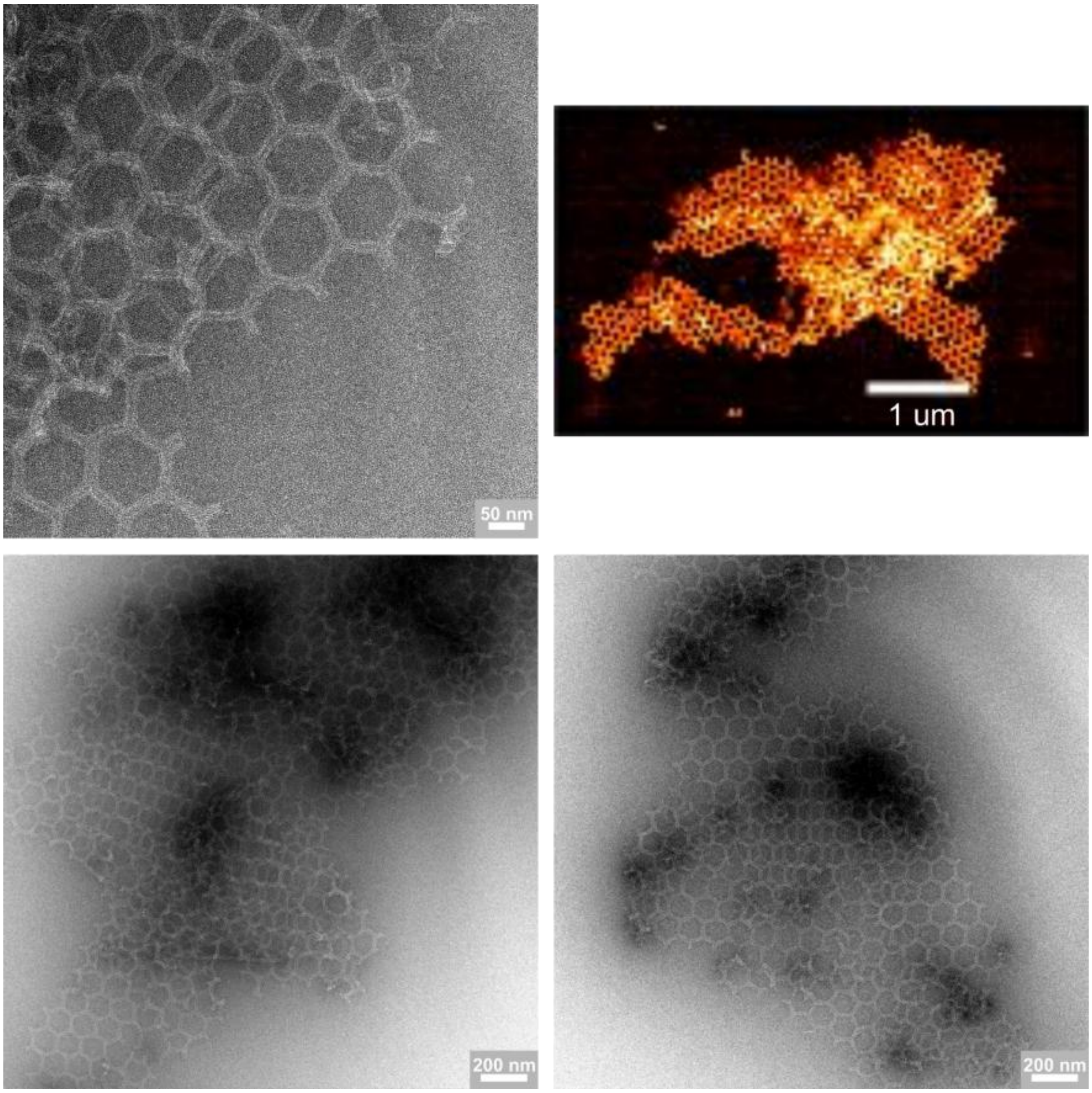


**Fig. S25.** Extended AFM and negative stain TEM images of the traditional honeycomb lattice formed by assembly of the 30 HB Y-junction. Scale bars as labelled.

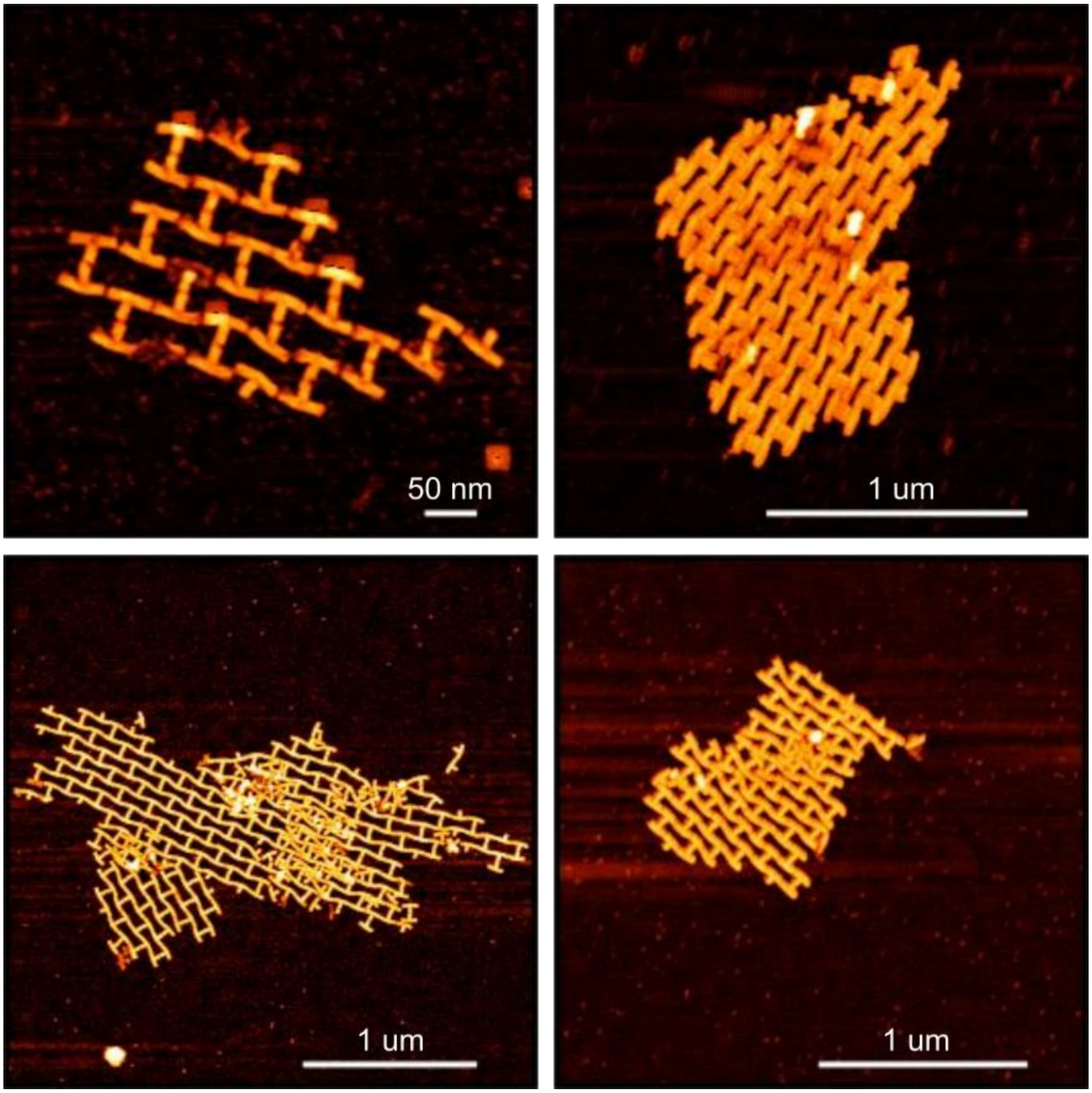


**Fig. S26.** Extended AFM images of the reentrant honeycomb lattice formed by assembly of the 18 HB arrow. Scale bars as labelled.

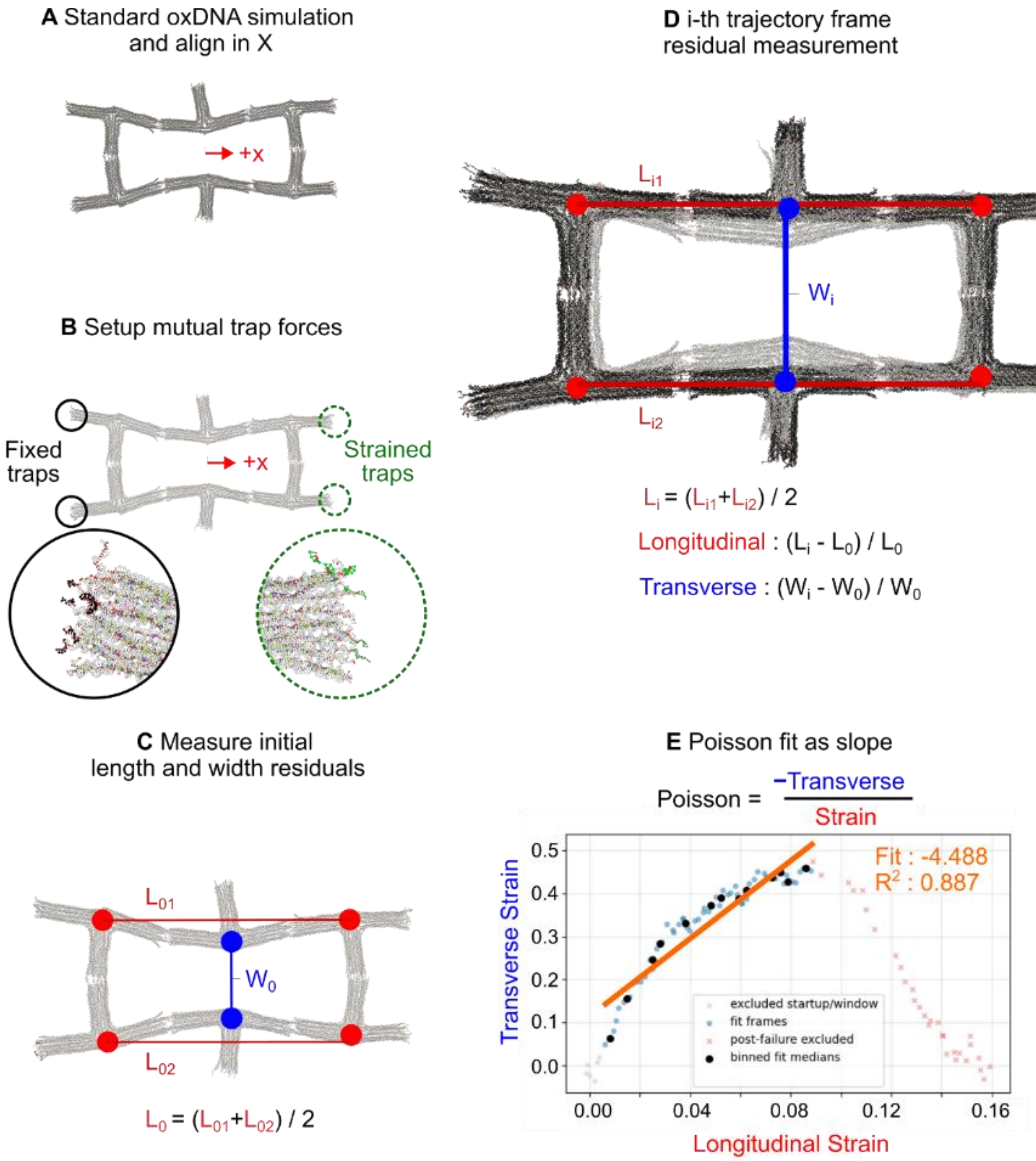


**Fig. S27.** (A) A unit cell is first simulated using a standard minimize-relax-simulate (see Materials & Methods) and then aligned in the X direction inside of oxView (38). (B) Harmonic trap forces are added to the 8-nucleotide long sticky end nucleotides on each face as circled and shown. (C) The centroid configuration is measured in width and height using the average position of a group of nucleotides at the central vertex of the arrow. Here, only the directional components are used to study the uniaxial loading case. (D) An oxDNA simulation is run for 1e7 steps and the per-frame longitudinal and transverse strains are calculated from the original length and widths. (E) The transverse and longitudinal strains are plotted against each other, and the Poisson ratio is the negative slope of the line fit through the individual frames.

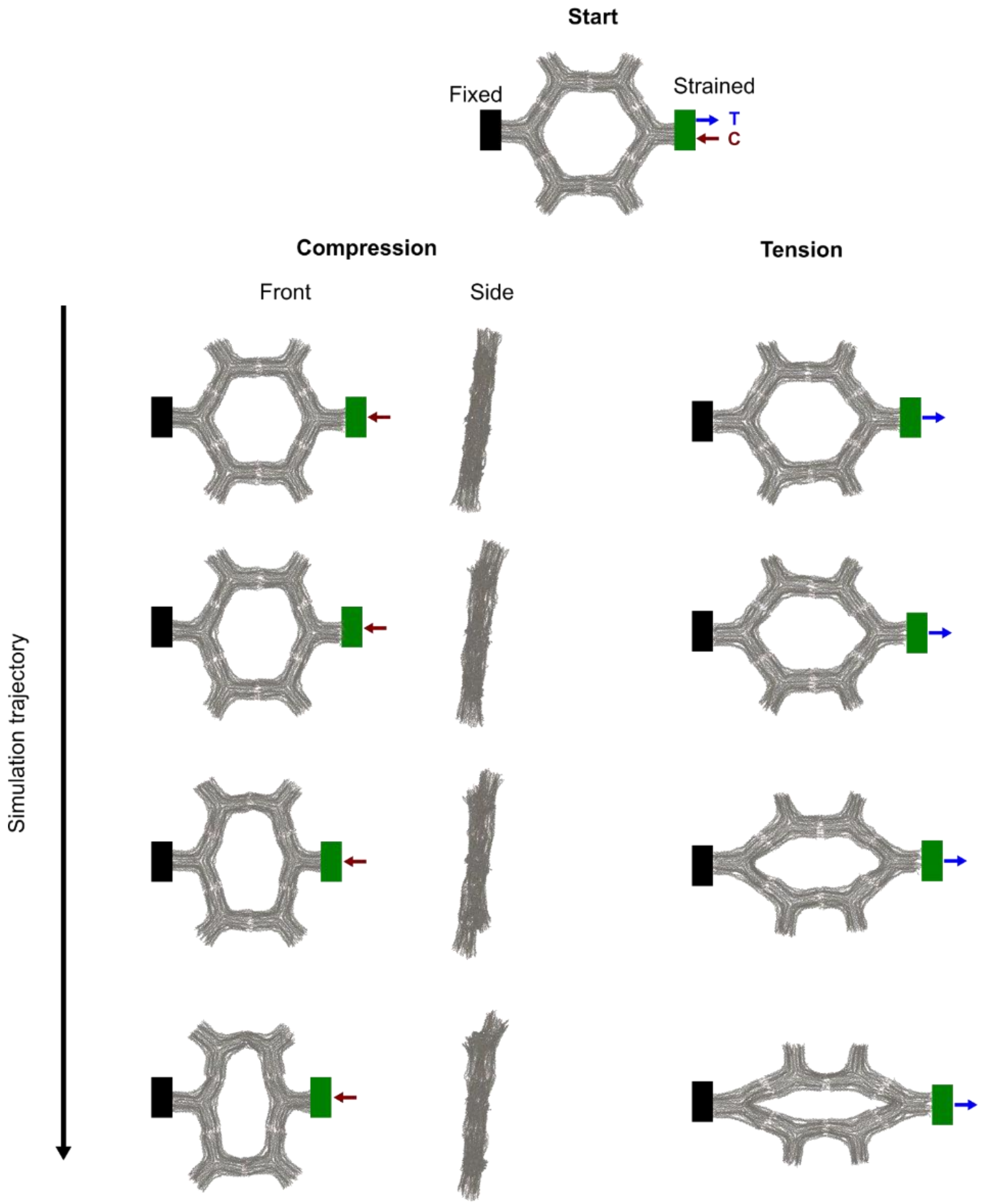


**Fig. S28.** Select states shown during compression and tension simulations of a single traditional honeycomb unit made up of six 30 HB Y-junction units (supplementary video 1).

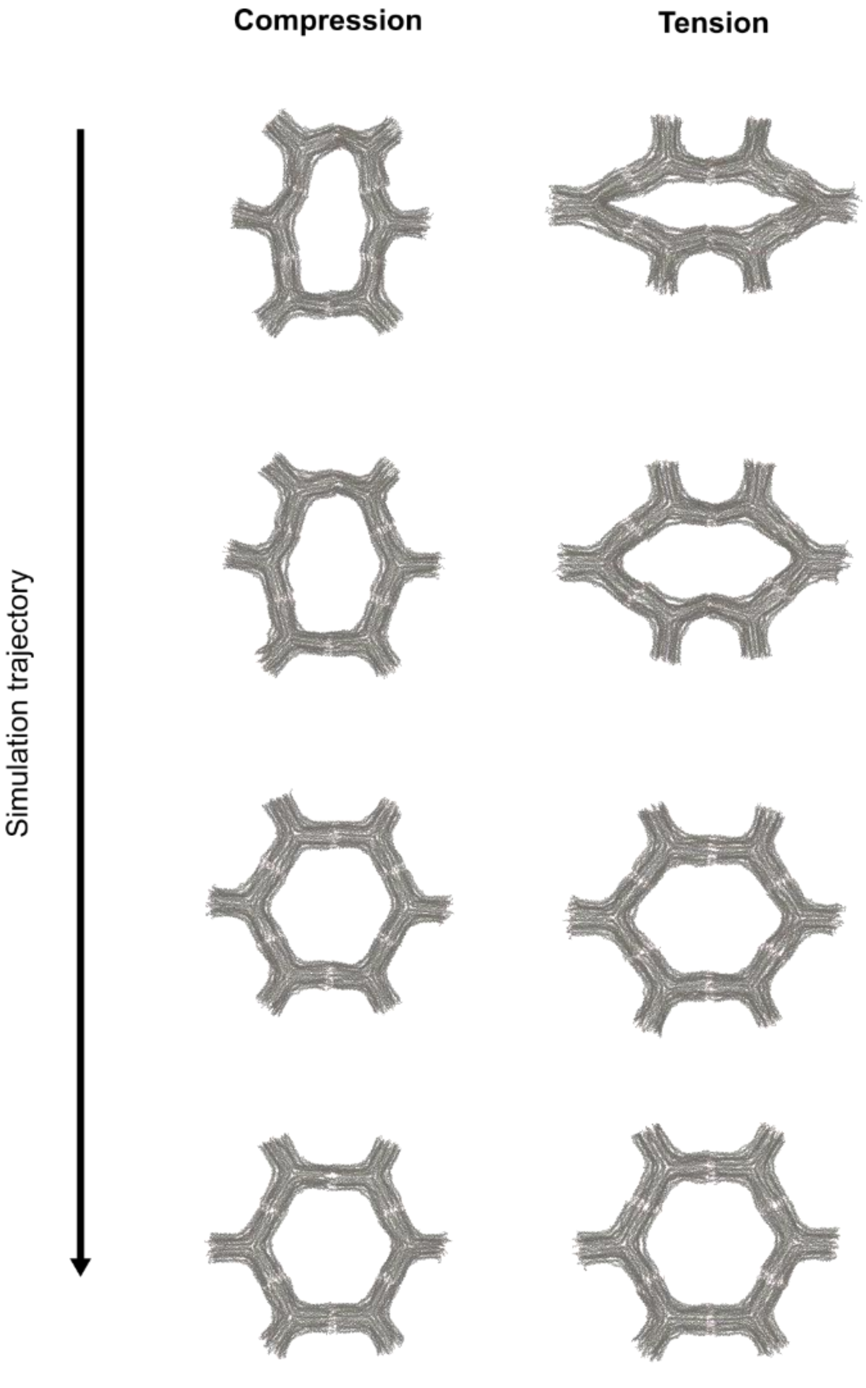


**Fig. S29.** Standard oxDNA simulations showing the inherent recovery of the initial geometry starting from the most compressed and tensile states found during the strain test of the single traditional honeycomb unit (fig. S28) (supplementary video 2).

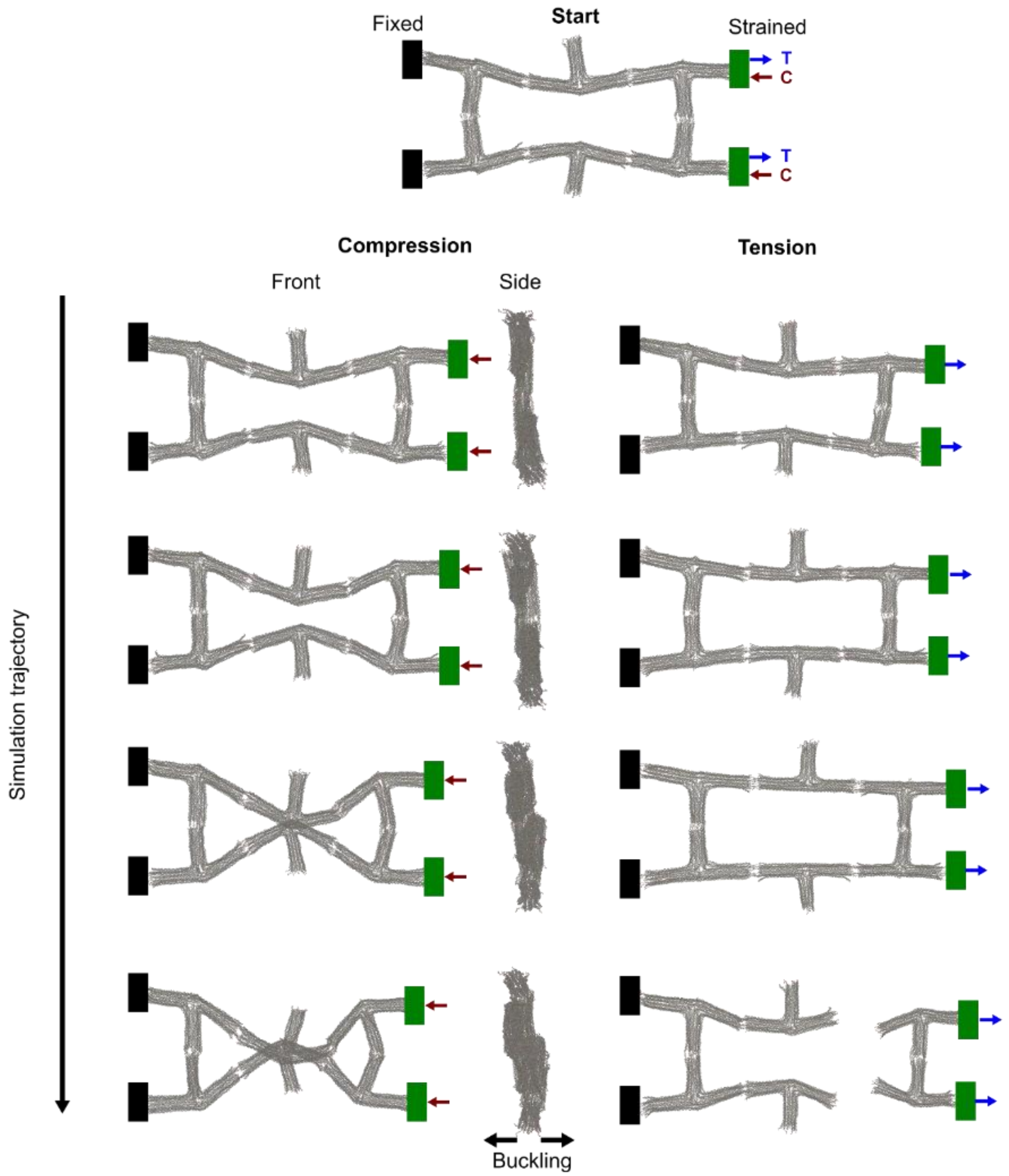


**Fig. S30.** Select states shown during compression and tension simulations of a single reentrant honeycomb unit made up of six 18 HB arrow units (supplementary video 3).

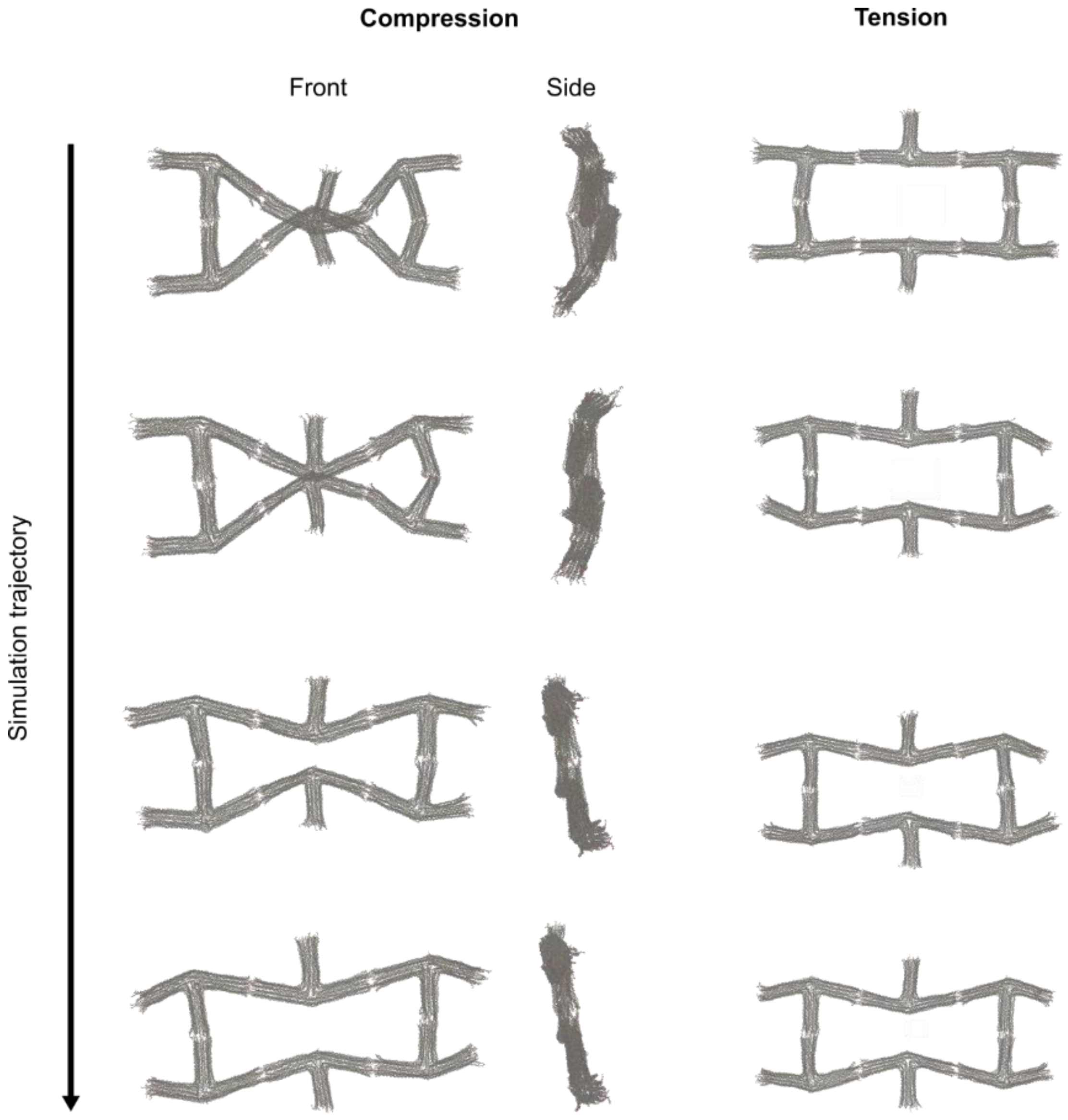


**Fig. S31.** Standard oxDNA simulations showing the inherent recovery of the initial geometry starting from the most compressed and un-failed tensile states found during the strain test of the single reentrant honeycomb unit (fig. S30) (supplementary video 4).

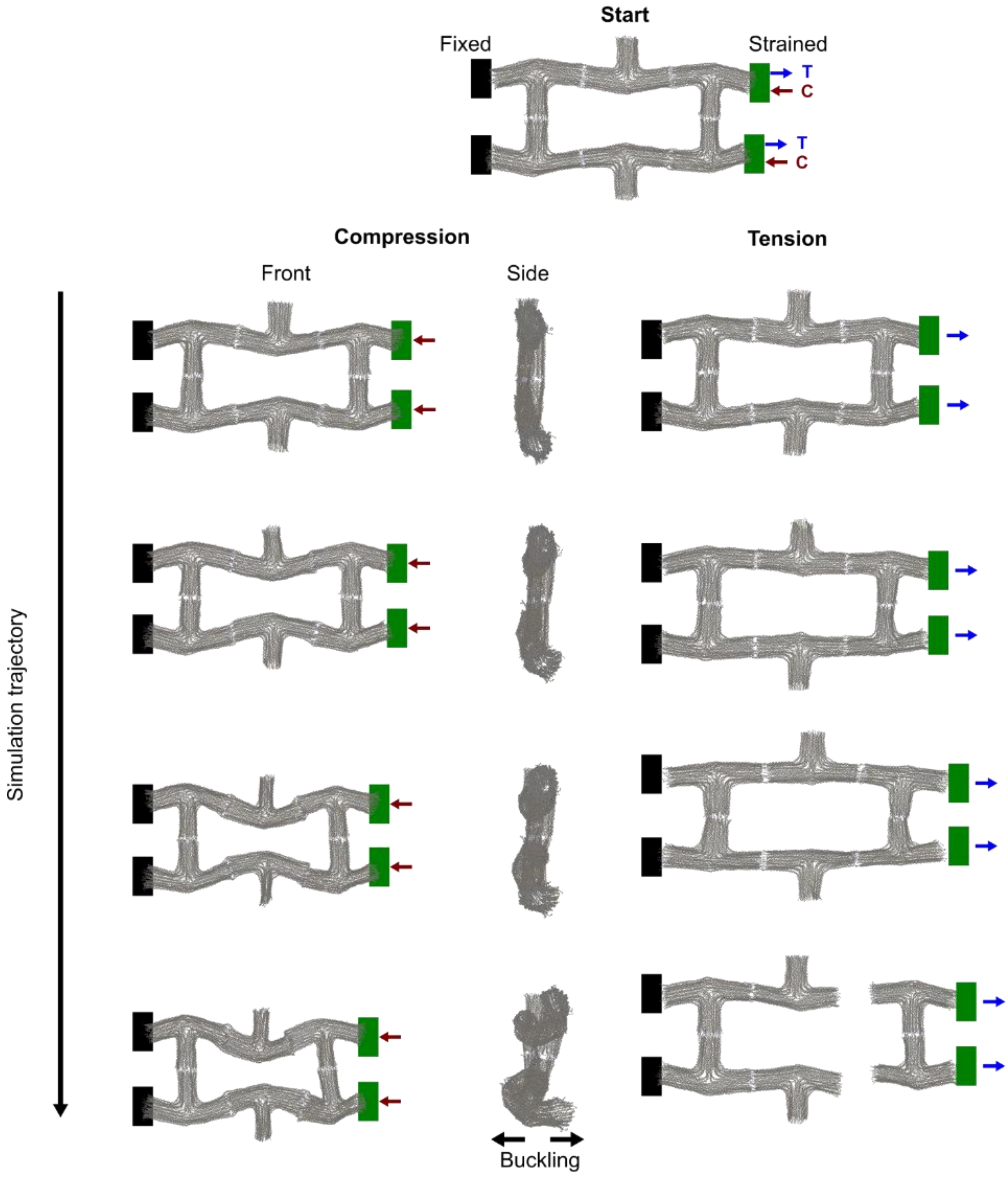


**Fig. S32.** Select states shown during compression and tension simulations of a single reentrant honeycomb unit made up of six 30 HB arrow units.

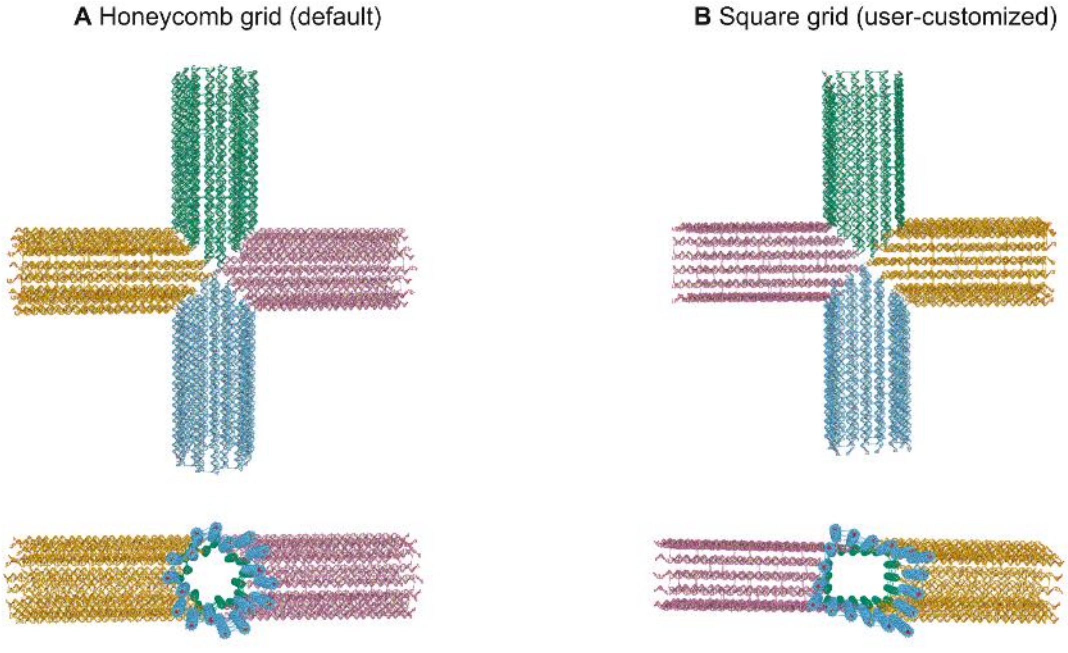


**Fig. S33.** The presented paradigm can be used with non-honeycomb style grids, for example the square grid can be used in place.

**Table S1. 18 HB arrow staple oligonucleotide sequences**

| Count | Name | Sequence |
|---|---|---|
| 1 | Arrow_Core | GTTACATATACTGCGTAAATAAACAGCCATATTAATTTGCTA |
| 2 | Arrow_Core | TATTTCAATTCTGCGAACGAGTAGATTTACGGGAGAAATCACCA |
| 3 | Arrow_Core | AAAGGGATTTGACGAGCACGTAAGCCGGAATCGGAAGAATAGG TTTGAT |
| 4 | Arrow_Core | GGTAAATCGTCTGAAATGAACGAACGCAGCAAGCGGAATATA |
| 5 | Arrow_Core | GAAGAAAACCGGAGTGTTGCCCCAGCAGGCGAGCGCCAGGGT |
| 6 | Arrow_Core | TCGGGTAACACTGAGTTTTTGTCGTCTTTCCAAAACACTTGTG |
| 7 | Arrow_Core | AACAGTTAGTACAAACTACAATTAGCGTAACGATCAGC |
| 8 | Arrow_Core | TTTTCAGAATTTCTGGCATCATAAAGCCTTTTCAGAGCATAAA |
| 9 | Arrow_Core | GACGTGTCACACCCACAAGAATTGAGTTAAGCGAATAAGTTT |
| 10 | Arrow_Core | CCTCAATCAATTATTAGTCTTTAATTAAAAGGCTTTGATGAT |
| 11 | Arrow_Core | ATAGGTCACGTGGATTCTAAATCAGTTGTAAAAGAGAAT |
| 12 | Arrow_Core | ATTAGGCAACAAGTTGCACCACTATTAAGAGCAGATTAAGAA |
| 13 | Arrow_Core | CACGTCACCAGTGAGACGATTGCGTTGCGCTCCGAGCCGCTG |
| 14 | Arrow_Core | GAATACCACATCAAAGCGAACCAGATTTGATAAGAGGTCAAT |
| 15 | Arrow_Core | CGCGTAACCACGCAAGTGAACTAATCGTCAATAGAT |
| 16 | Arrow_Core | TCAATAGAGTAGAAGAACTCTACATTGGCAGATTACATCGCGCA |
| 17 | Arrow_Core | CCACAGCCAGAGAGAAGGATTAGGAGCCACCCTCAGAGCACA |
| 18 | Arrow_Core | CAAATCCAACATGTTATTAAACTCATTAAGGAATTACGAGGCGAA |
| 19 | Arrow_Core | GCCCGAAAGACTTGCGGGATCGTCATTCGGTCTACATTTTAT |
| 20 | Arrow_Core | CACCTCCTCAAATGGAAAGCGCAGTAGCCGCCACCAGAACAGAAT |
| 21 | Arrow_Core | TAAACATTAAAAATACCGGATTATTAAACTATACAGTCAAGCCTT |
| 22 | Arrow_Core | TTAGAGCTGAGGCTTGCAGGCCGCTTTTCAAATCGCCAAATAC |
| 23 | Arrow_Core | ACCCCTGCGGAATTAGCCTTACTCGTCATAAACCAACCTAAA |
| 24 | Arrow_Core | GAGGCATATGGGCGCTAAGGGAGAATTAACTGAAAA |
| 25 | Arrow_Core | TAGCAACGGCCGACAATTTGGGGCGCGTTTTAGCTGAAAAGGTTA |
| 26 | Arrow_Core | TTTTTCAGAGTTTTATAGAACCCTTAGCGTAAAATATCTAGG |
| 27 | Arrow_Core | TATACCCTCAGTTAAGAGATAGTAAATTTTAAGAACTGGCCAAGTA |
| 28 | Arrow_Core | CGATGAACAGGAACGGTACGCCAGGGCGCGTACTAAAG |
| 29 | Arrow_Core | TATAGAAGGCTCAACATGTAATTTAATACCGATTAATGGTTA |
| 30 | Arrow_Core | CAAGTTTGCGACTTGCATATAAAATGAAAAGTTACAAAATCGCAAA |
| 31 | Arrow_Core | AGACTATCAGAGAGATAAATCAATAGAAAATTGAGGGA |
| 32 | Arrow_Core | ATAATTCCCAACGCAAGGGAGAAAGGCCGGAGCGGCCTTGCT |
| 33 | Arrow_Core | TATCACCGTACAATCTCCAAAAAAATTATCAGTAGTAGCAAG |
| 34 | Arrow_Core | AGATCTAGCAAAATTAAGCAAATTCTACTAATAGCTTGCTTTCG |
| 35 | Arrow_Core | AGGTAAATGTCATAGTCAGAGCATGATACCCTGCCTATT |
| 36 | Arrow_Core | TTCAGAAATTCCATTAAACGGGGTGAGAACAGAACCTTA |
| 37 | Arrow_Core | GATTTTGTATCCCAACTTGGCGCATAGGCTGGGCACCC |
| 38 | Arrow_Core | ACAATAATTGAATTACCTTTTCCGTGTGATAAATAACA |

| | | |
|---|---|---|
| 39 | Arrow_Core | CGCCCCTTATCTTGGGAAATGCAGATACATAAATCGCGTTTT |
| 40 | Arrow_Core | TGTTCCGTGGGAACAAACCCGTAATCTTCTGGCAGGCAAACG |
| 41 | Arrow_Core | TTGCGAACCCTCAGTTTTAACCGCCACCCTTAGAAAGGAA |
| 42 | Arrow_Core | GAGAAACCAAAATAGCGAGAGTTTGCTTTTGCAAAAGA |
| 43 | Arrow_Core | CATGGCTTTTGCGCCACCCTCAGAACATCGGCATTTTCGATT |
| 44 | Arrow_Core | GTTTGGAAACGTCAAAGGGCGAGGAAGAAAGCGAAAGCC |
| 45 | Arrow_Core | GAGCCACCACGTCAGACTGTAGCAAGGTGACACCACGCCAATAA CGA |
| 46 | Arrow_Core | GCCATTCAGGCGCAAGGCGATTAAGTCGACTCTAGAGGATTC |
| 47 | Arrow_Core | GCATTTTTTTTCGAGCCAGTACCTAAATTTATTTTATGGTTTGAAG GCAGAG |
| 48 | Arrow_Core | CATTTTGCGGACTTGCTGAACCTCACTGATAGCCCTAAACAC |
| 49 | Arrow_Core | CTTTGAGGACTTTTTAAAGACTTTTTCATGTAATTTTAGGAA |
| 50 | Arrow_Core | CCCTGAGAGGCTGATTGCCCTTCACAATACAACGGCGCCTGG |
| 51 | Arrow_Core | AGGCCTCCATGGGAACGAAGAACCGGATATTCCGGGAGGTTT |
| 52 | Arrow_Core | TCGATCATAATCATCAAGAGTAATATCAAGATTAGTTGCCA |
| 53 | Arrow_Core | AGTGAGCTAACTAAAGCCTGCCAGTGCCAGTCATTGAT |
| 54 | Arrow_Core | TGAATACCAGTATAAAGCAAAAGCCATGAAACGCAGAGGTAAGA GCA |
| 55 | Arrow_Core | CAGGTTGGGTAAGCGCCATTCGCCAGCTTTCCGGCACCGGGG |
| 56 | Arrow_Core | CCTGAAAAGAATACACTAGACGTTAATGTACCAACCTATATA |
| 57 | Arrow_Core | CCAGGGTTTTCCCAAGCTTGCATGCGAAGCATAAAGTGTCAC |
| 58 | Arrow_Core | TATCCATAAAGGTGGCAAAGCCATTTGGGAATGTAGCGACCA |
| 59 | Arrow_Core | CATATTACCGAGCCTGATTGCTTTGAATTACATTTAACAAAA |
| 60 | Arrow_Core | TTTTCTGACCTGGGAACCGAACTGAATCGCCTGATAAATCAT |
| 61 | Arrow_Core | GTACTGGTCGGAACCGCCTCCCCCCCCTTATTAGCGATT |
| 62 | Arrow_Core | AATTCGGATGGCTTAAAGGGAGTTAGAGCTTAATTGCTGATT |
| 63 | Arrow_Core | AGCGTGAATCCCCCTCAACGTAATGCCACTACACTTTCACAC |
| 64 | Arrow_Core | TTTCCCTAGACAAAGAACGCGGTCCAGACATCGAGTTATGCGGAG |
| 65 | Arrow_Core | CTTTGACCCCCTAAAGTTCGTCACCAATGCCCAGGAGT |
| 66 | Arrow_Core | ATTATTCATTTAGCAAAAGAAGATGTGTTTAGTATCATAAAA |
| 67 | Arrow_Core | CTCCTTATTACGTTTTCAGTATGTTAAAGTAAGGGAGAA |
| 68 | Arrow_Core | CAGTCAGGACGATTCCAAGAACGGGTCAGCTAATGCAGATGT |
| 69 | Arrow_Core | CATTAACAGAGCCGCCGCCAGCATTTTTGACAGGAGGTTGA |
| 70 | Arrow_Core | TAGGGCTTACCGGAACAAAATTAATACCATAGCAATAGC |
| 71 | Arrow_Core | TAAACTCATTTTTTAACCAGTAACAACCCGTCTGGTGTACCA |
| 72 | Arrow_Core | TTTGCGAGCTTTCAACTAGAAAAATCTACGTTAATCGGCTGT |
| 73 | Arrow_Core | AGAAACAAGAAACGCAAAGAATTATCACCGTCACCCT |
| 74 | Arrow_Core | TCCAAAAGGGTATAAAAATTTTTAGCATATAACAGTTGATGC |
| 75 | Arrow_Core | AGTATTAGACTTTACAAACAATTCGTTTTACAACTCGTAAATTTTG GAAT |
| 76 | Arrow_Core | AGGTCGTTGAATCAGGAGACCAGGCCACAAACAAATAATTTTAT CCT |
| 77 | Arrow_Core | CGTGGCGCTAGGGCGCTGCACACCCTATAATCGTC |

| | | |
|---|---|---|
| 78 | Arrow_Core | GCAACTGTTGGAGGAAGATCGCACTGATGGGCGCATCGTGTG |
| 79 | Arrow_Core | TTCAGGGAGATTTTGCTAAACAGAAGGCATATTCATTCC |
| 80 | Arrow_Core | GTTTCAGCGGATAAAATAATGCTTTAAATGTTTCATTCAGTG |
| 81 | Arrow_Core | ATATTCCCACGCTGAGAGCCACACCAGCAGAAGATGACGCTCATA |
| 82 | Arrow_Core | AGTTTGACCATAAATCAAAATTTTATCAGGTCTTTACCCAGG |
| 83 | Arrow_Core | GGCAAACCATCGATAGCATTTTGCACCGTAATCATAGAGCCATA |
| 84 | Arrow_Core | ATTTGAAATTATTCATTAGCGTTTTCCGCCACAAGCGTCTAT |
| 85 | Arrow_Core | AGCTACAAGTCTTTCCAGAGCCTATTTATAGCCTTTATT |
| 86 | Arrow_Core | GGTTGCTTTCCAGTCGGGCTCACAATCCCCGGATGTGCTTGC |
| 87 | Arrow_Core | TGTCCATCACGCTTTAAATTAACCGGTAGTTTTCTATTTTTGAG |
| 88 | Arrow_Core | TGTGAGTGAATAATTTCATCTTCTGAATAAGACATTACCCTT |
| 89 | Arrow_Core | TCCTCCCGAGATAGGGTTTCTATCAAGCACTACGAACG |
| 90 | Arrow_Core | AATACATTTGTTAGGAGCACTAACTAGCGGTCACTGAACTGACCG CTG |
| 91 | Arrow_Core | TCATCGGATTCAGCCCTTTTTAAGAAGCAAACGTAGAAAAGC |
| 92 | Arrow_Core | TTGGCAAATCAACAGTTGATTAAATACGTCAGATGAATAATA |
| 93 | Arrow_Core | TTCTGCCTTAACTTGACAGGCGCAGACGGTCAAAATCCGCGA |
| 94 | Arrow_Core | ACGCTATCCGGAGGCTTGAGGGGGTAATAGTAAAACAG |
| 95 | Arrow_Core | AGATTTTCCTTCTGTAAATCGTTTTTTCAAATATATGACAAAATTT |
| 96 | Arrow_Core | TAATTCTAGAAAACCGCTATTAAAGAAATTTTAAAAGTTTGAAAC |
| 97 | Arrow_Core | TGGCGAGATGGTTGCTTTTAGACGGTAATAGCAAATATT |
| 98 | Arrow_Core | CCGCACTCGACGACAATAAACAATCGCATAGAATCATTTGCATAT |
| 99 | Arrow_Core | TACCTTTAGTTAACCTTGAGGTTTACCTTTGCCCGAACGCAG |
| 100 | Arrow_Core | CGAGTTTTAGTAAATTGGGGCGCCCAATATTTTGCAAGCAAATCA CCAGAA |
| 101 | Arrow_Core | ATAAGAAACTGCCGGTATAGCCCAAAAGGAGCATTGAGGCGGCT TT |
| 102 | Arrow_Core | ATTGCCTGAGAGTCTGGACTGAGAAGTGTTTTGCCGCG |
| 103 | Arrow_Core | AAAATCACCATTTGTAGCACCATTAATGATTAAAACCGAAGA |
| 104 | Arrow_Core | AGCTCGAAGAAATTGTTATCCGAAACCTGGTATTGGAAA |
| 105 | Arrow_Core | GCGGGGTTTTTTTGCTCAGTGTTTAGTACCGCCATAATAATT |
| 106 | Arrow_Core | CATCGCAGATAGTTTTCCGAACAAAGTTACCAGAAGGAGA |
| 107 | Arrow_Core | CAACACTATCATTGCATCAAAAAGACAGCGAAAGACAGCCGC |
| 108 | Arrow_Core | AAATTGTACCAAAAACATCGCAAATGGTCAATGCATAACCGA |
| 109 | Arrow_Core | AGCGAATAGGATTGTATACGTAAAACTAGCATGCCGATT |
| 110 | Arrow_Core | TGAGGAAGGTTATCTAAGAATACGTGGCATTTCAGACAATA |
| 111 | Arrow_Core | ACTTGACATTCTGGCCAAGAATGGCATCTGGTTTATTAATTGCGT |
| 112 | Arrow_Core | CATCAAGAAAATCATAATTACTAGACAACGCTCGACTTGATT |
| 113 | Arrow_Core | TTAGCCCTCAGCTCTGAATAAGAGGCTGAGACCTCATT |
| 114 | Arrow_Core | AAATGATAGCTAGAACCTAACCACCAGAAGGAATGAAAAATC |
| 115 | Arrow_Core | ATGGAAGGGTTTAGATTAAGACGCTGTTATATAACTATAACG |
| 116 | Arrow_Core | TAAGAATAAACAATTGAGGAGGCGTAAGCTGCTAGACTGGAT |
| 117 | Arrow_Core | CGGATAACCCTCATTGTGAATTACCAACAAGCAAGCCGTGGTAA AG |

| | | |
|---|---|---|
| 118 | Arrow_Core | TGTAGCTCAACTGCTCCTCCGGAAGTGAGATTCGAACTAAC |
| 119 | Arrow_Core | GCTACAGAATCGCAAACACGTTAATATTTTGTATTAAATTTT |
| 120 | Arrow_Core | CAGTCTTGCCTTGATATTCTGTAATACTTTTGGTTTGACCA |
| 121 | Arrow_Core | AATTTTAACCTCCGGCTTAGATAAGAATCAATAATAAAATAG |
| 122 | Arrow_Core | TCACACAAAGAACCATATCAAAATTCTTGAAAACATAGCGCTGATG |
| 123 | Arrow_Core | CTTTTGTTTATCAACAATAGGTTGGGAGAAGACTTCTGATATCATC |
| 124 | Arrow_Core | AATATATTCTAAGAACGCAATCGCCATATTTAAGGCGTTATT |
| 125 | Arrow_Core | ATAAATTCGCTAAAATCATACAGGCAAGGCAATTAACATCCA |
| 126 | Arrow_Core | AATTACAAAGGCTATCAGAGTGAGGCCACCGAGTAAAAGAGTC |
| 127 | Arrow_Core | TATTCTTTAATCGTTTACCAGACGATATAGTCAGAAGCACGAGGG |
| 128 | Arrow_Core | ACGATGTATGGTAGCAAGAAAGTATTTTACCGTTCCAGTCCTCA |
| 129 | Arrow_Core | CTTAATGCGGAGCGGGACTCCACAAGAGTGCAAGCGGTC |
| 130 | Arrow_Core | TCAATTACAGTAACTTTTAGTACCTTAAACAGTACTTTATAAA |
| 131 | Arrow_Core | GGCCCCAATGAGGTCAGATGATATTGGATAAGTGCCGTCGTG |
| 132 | Arrow_Core | GTAAGCGCGAAAATATCAGTAACATCGTAAAACAGAAATAATTAAT |
| 133 | Arrow_Core | TCTGAAACATGCCCAATAGGAACCCGTAAATGAATTTTCAAG |
| 134 | Arrow_Core | CCACAACCTGTAATCGGTTAATGCCGGAGAGGTTGTAGCAAT |
| 135 | Arrow_Core | ATTTGGAAACGCAATAATCCAAAAGAACTGGCCCATTAGCAA |
| 136 | Arrow_Core | AACAGCTTGATACTTTTCGATAGTTGCGCTACAGTGACTATCGA |
| 137 | Arrow_Core | ATAACATACCTGCAATTAAAAACAGGGAAGCGCACAGAGAGA |
| 138 | Arrow_Core | AATAAAATCAACGTAACATTTAGCGAACCTCCCAACAG |
| 139 | Arrow_Core | TAAATGGTTTAATTTCAATTCATCGTAGGAATGAATATAAAG |
| 140 | Arrow_Core | GCTATTAGCTATATTTTCATGACAACAACCATATCGGAAAAG |
| 141 | Arrow_Core | CAACAGGAAGTACGAGAATTGCCAGCCCTGACGAGAAACAGA |
| 142 | Arrow_Core | GACCCAACCGTTCTAGCTTAGTAATAACATCACACACGACCA |
| 143 | Arrow_Core | CACACAACATAACTGCCCTTTCTTTCTGGTTTGTTCCA |
| 144 | Arrow_Core | AGGAAGAACGCCATCAAAAATAATTCGCAAAG |
| 145 | Arrow_Core | CCTCTAGGTAAAGATTCAGAACAA |
| 146 | Arrow_Core | AATTGTATCGGTAGGCTCCGGAATAGGAGAGGGCGACGTTGT |
| 147 | Arrow_Core | TATTACCGAATACCTACATTTTAAAACAGAGGTGA |
| 148 | Arrow_Core | GGAACAGCATGTAGAAACCTCCTGAACAAGAAATAC |
| 149 | Arrow_Core | GGCCTCGATTAAAACGACGGGGGTGCCTCGTCAGGAAAAATG |
| 150 | Arrow_Core | ACAGGAGAGTACCTTTAATATGTTT |
| 151 | Arrow_Core | GATTGTTTGGATTATAGTCAATAGTG |
| 152 | Arrow_PolyT_Overhang | TTTAAAGTCAGAGGGTAATTGATTTACCAGCGCCAAAGACATTT |
| 153 | Arrow_PolyT_Overhang | CGGCCTCGAAGGGCCTGGCGAAAGGGGGGTACCG |
| 154 | Arrow_PolyT_Overhang | GGTGGTTGAGAGGCGGTTTGCTCGTGCCAGCTGCATTAATGAATC GGTTT |
| 155 | Arrow_PolyT_Overhang | TGAAAATAGCCCCAATCCAAATAAGAAACGATTTTTTTT |
| 156 | Arrow_PolyT_Overhang | TTTACCAACGCTAACGAGCTTTTATCCTGAATCTTTTT |
| 157 | Arrow_PolyT_Overhang | TTTCCAACGCGCGGGCCGAAATCCTTATAAATCAAAACCCTAAAGG GAGCCCCCGTTT |
| 158 | Arrow_PolyT_Overhang | TTTCCTTGAGTAACAGTGCTTTAACGGGGTCAGTGTTT |

| | | |
|---|---|---|
| 159 | Arrow_PolyT_Overhang | TTTATTTAGAGCTTGACGGGGAATAACGTACAGGAGGTCAATCCAAAAAC |
| 160 | Arrow_PolyT_Overhang | TTTATCACCGGAACCAGAGCCACCACAATAAGTCCGTATA |
| 161 | Arrow_PolyT_Overhang | TTTAAAGTACAACGGAGATATACCAAGCGCGAAACTTT |
| 162 | Arrow_PolyT_Overhang | TTTATAATCAGAGTCTGGCCTTTT |
| 163 | Arrow_PolyT_Overhang | TTTATTAACACCGCCTGCAACAGTGCTGATTATCAGATGATGGCAATTCTTT |
| 164 | Arrow_PolyT_Overhang | TTTTTATTAGTCAGGTTTTT |
| 165 | Arrow_PolyT_Overhang | TTTATGCCTGAGTAATGTGATATATTTTAAATGCATTT |
| 166 | Arrow_PolyT_Overhang | TTTCTGAGAGACAATAATATCCCATTT |
| 167 | Arrow_PolyT_Overhang | TTTTGGGACGACGACAGTAT |
| 168 | Reentrant_Red17to8_5p | GCGGTAACTTCCTGTAGCCAGCTTTCATCAACATTAAATAACCGTGCAT |
| 169 | Reentrant_Red8to8_3p | CTGCCAGTTTGAGTTTTTTTTTGTGCCAG |
| 170 | Reentrant_Cyan7to14_Both | GTTACCGCTTTTTAAATCCGGCATTTTTCTGGCACA |
| 171 | Reentrant_Yellow15to6_5p | GGTTGACCTTTTTTTTTTCGCTATTACGCCAGGATCGGTGCGGGCCTCTTT |
| 172 | Reentrant_Black11to2_5p | GGTCAACCTTTTTTTTTAGCTGTTTCCTGTGTTTCGTAATCATGGTCATTT |
| 173 | Reentrant_Yellow13to12_3p | CCCATATGTACCCCGGTTGTCCTGAACC |
| 174 | Reentrant_Black5to16_3p | TTTTCAGAGCGGGAGCTAAGCTTTCCTCGTTAGAATGGTTCAGG |
| 175 | Reentrant_Cyan40to49_5p | TGGAAGCCTTACGGTGTCTGGAAGTTTCATTCAAC |
| 176 | Reentrant_Red20to33_5p | GGCTTCCATTTTTTGTTTAACGTCAAACAC |
| 177 | Reentrant_Cyan53to53_3p | TAAATATGCAACTAAAGTTGCAGAGC |
| 178 | Reentrant_Red33to33_3p | CCTGAACTTTTTGCTCTGCA |
| 179 | Reentrant_Cyan34to23_5p | CGCTACGTTCAGACAGCCCTCATAGCGCCT |
| 180 | Reentrant_Red50to50_5p | ACGTAGCGTTTTTTTTATCAATATAATCCT |
| 181 | Reentrant_Cyan23to23_3p | GTAGCATTCCATCGAGACGT |
| 182 | Reentrant_Red46to39_3p | CTTTTATCAAAATCATAGGTTTTTTTTTACGTCTCG |
| 183 | Reentrant_Yellow26to35_5p | ACCATCGGTTTTTAAAGGGCGACATTCAACCGTTTGCC |
| 184 | Reentrant_Black42to37_5p | CCGATGGTTTTTTGGTAGAAAGATTCATCAGTCAAACTCCA |
| 185 | Reentrant_Yellow35to35_3p | ATCTTTTCATAATCAAATCCACTTGG |
| 186 | Reentrant_Black38to51_3p | TTTTCCTAATTTACGAACATTATTACATTTTTCCAAGTGG |
| 187 | Reentrant_Yellow52to41_5p | CCTCACGTTGAAAAACGCTCATGGACCAGCCATTGCAACAGTTT |
| 188 | Reentrant_Black32to25_5p | ACGTGAGGTTTTTTTTAACGGTGTACAGACCATGAAA |
| 189 | Reentrant_Yellow45to45_3p | GGCGGTCAGTTTTTTTTTCCATGCTG |
| 190 | Reentrant_Black25to25_3p | GAGGACAGATGTTTTTTTTCAGCATGG |

**Table S2. 30 HB Y-junction staple oligonucleotide sequences**

| Count | Name | Sequence |
|---|---|---|
| 1 | Y_Junction_CoreStaple | TGTCTGGAAGTGCGAACGAGTAGATTTTTTAGTTTGACCATTAG |
| 2 | Y_Junction_CoreStaple | CCATAACAAGAGAATCGATTAAATTCAAACGGTCAGGAAG |
| 3 | Y_Junction_CoreStaple | AGACAAGGATAAAAATTTATTAACAGTTTAGCGAATAT |
| 4 | Y_Junction_CoreStaple | ATCGCACGCCAGCAGAGGATGGGGTGCCTAATGACGGGGAG |
| 5 | Y_Junction_CoreStaple | AATGCTGTCAGGTCAGGATTAGGAAAATTAAAGGTGGCA |
| 6 | Y_Junction_CoreStaple | TCTTTCCAGAATCATAGAACGGAATCGCCATATTTAATG |
| 7 | Y_Junction_CoreStaple | TGGTCAACATTAAATGTGAGCTTTTGAGTAACA |
| 8 | Y_Junction_CoreStaple | CAGTCTCTGAATTGTAAGCGTCATTTTACATGGC |
| 9 | Y_Junction_CoreStaple | CTAAAACGATTTTTTGTTCCCACAAGAATTGATAAGCAGCAT |
| 10 | Y_Junction_CoreStaple | AGGGAAGCGGAGAATTAACTGAATACCCACATGATTCGA |
| 11 | Y_Junction_CoreStaple | GAGCACGAGGAGCGTGAGTGTCAGCAAGCGGTCCACAA |
| 12 | Y_Junction_CoreStaple | GATAAGTCCTGACCTCCCGACTTGCGAAGCCTAAAAAC |
| 13 | Y_Junction_CoreStaple | TTCATTTGCTAATAGTAGTAGCTTAGAACAGGGTGAGA |
| 14 | Y_Junction_CoreStaple | CAAAGCTTTGAACCAGACCGGAAGCAAGCGCCAAAAAC |
| 15 | Y_Junction_CoreStaple | AAGGCTAATCGTAAAACTAGAAGATTTGGGATAGGGACG |
| 16 | Y_Junction_CoreStaple | AAATTGTTAAAGAAATCTATCCGCTCCAGTGAGACAAACAAGA TAGGGCA |
| 17 | Y_Junction_CoreStaple | CGCGAGGCGTTTTGCTATTTTTTTGCACCCAGCTACAGGC |
| 18 | Y_Junction_CoreStaple | AATGAAAATAGCAGTTTTCCTTTACAGAGAGAGAACAAA |
| 19 | Y_Junction_CoreStaple | ATAGTAAAAGGATAGCATGACCAATCAGGTCTTTACCAGC |
| 20 | Y_Junction_CoreStaple | CGCGGTGAGCTTGCAGGTCGACTCTTGGCGAAAGGGGGAATC |
| 21 | Y_Junction_CoreStaple | GAGCTCTTTGAATTCGTAATCATGGTCATAGGGCGATCGG |
| 22 | Y_Junction_CoreStaple | CCTGAGAGTCTTTGTTAAAATTCGCATTAATTTATTTTTGTTA |
| 23 | Y_Junction_CoreStaple | TAATGCAGAACAAATAAATCCTCATATGATACTAAGAG |
| 24 | Y_Junction_CoreStaple | CTGCGCGTTGCCCAGTGCCAAGCTTCAAGGCGATTAAGTGTT |
| 25 | Y_Junction_CoreStaple | GCAATTCGCAACATGTTTTAAATATGCAACTAAATTTTGTACGG |
| 26 | Y_Junction_CoreStaple | CAGTTATAGTCAGAAGCGCATCAAAAAGATTTATT |
| 27 | Y_Junction_CoreStaple | ACCCTCAAATGCCCCCTGCCTATTTCGGGGTTTAGTACCGCC |
| 28 | Y_Junction_CoreStaple | AGGCGGTTTGCGTATTGGTTTGCGCCAGGGTGGTTTTTCCCC |
| 29 | Y_Junction_CoreStaple | TGGTTTAATGATTTTAAGAACTGTTTTTTACTTTCATGCC |
| 30 | Y_Junction_CoreStaple | CAATCCACCAATAATAAGAGC |
| 31 | Y_Junction_CoreStaple | CCATGTACCGTTTTTAACACTGAGTTTCGTCACCACCA |
| 32 | Y_Junction_CoreStaple | AGCTTCTTTAATTGCTCCTTTTATTTTGTCACAATCAATAAGAG TACAATTGCTTATATT |
| 33 | Y_Junction_CoreStaple | AACGTTAATATTGGAGCACAATATGGAGAAGCCTTTATTCAG |
| 34 | Y_Junction_CoreStaple | TTTTGTAAAGCCCATGTTCAAAATAATATCCCAT |
| 35 | Y_Junction_CoreStaple | ATACATGGCAAAGAATTAGCAAATTTTATTAAGCAATAAAGAAA |
| 36 | Y_Junction_CoreStaple | GGTCACGCTCGCCGCTCAATA |
| 37 | Y_Junction_CoreStaple | CCTCGTTTACAAAAGGAATTACGAAACAACAGCTCATTGAC |

| | | |
|---|---|---|
| 38 | Y_Junction_CoreStaple | CGTCTGAAATGGCCTAAATTCCAATCGAGAGACTACCTTTGAT |
| 39 | Y_Junction_CoreStaple | TGCGGGTTTTCCTCTTCGCTATTACTCCAGCATTCTCCGTGGGA<br>AGTA |
| 40 | Y_Junction_CoreStaple | GAAATATTAATGCGGGGTTTTAATTAAAATTGCTCAGTAGTA |
| 41 | Y_Junction_CoreStaple | TAAACCACTACGAAGGCAGAAACAAAGTACAAGTGTACACAG |
| 42 | Y_Junction_CoreStaple | ACGACAGTTGTGCTGGCATGCCAACTCACATTAATTATT |
| 43 | Y_Junction_CoreStaple | TGAATAAGGCTAAACGAACTAACGGGGCATAGACTATCATAAC |
| 44 | Y_Junction_CoreStaple | ACATCTTTTTAAGAAAAGGTTAAGCAATAAGACGAGCG |
| 45 | Y_Junction_CoreStaple | CATTCCGAATTTGCGTGAATTACCATCAAAGTCTGGCCATT |
| 46 | Y_Junction_CoreStaple | AGAATCAGACGGCATTAAGTTTGCCTTTAGCGCAGTAGCGAC |
| 47 | Y_Junction_CoreStaple | TATAAAAATAAGAATTTTTAAACACCGGAATCATAAGCCA |
| 48 | Y_Junction_CoreStaple | CGCGTACTGGCAAGTAAGAATATTGCCCCAGCAGGCCAG |
| 49 | Y_Junction_CoreStaple | ACAGCGTCTATCAGTGGCGAGTTTTAAAGGAAGGGAAGAAAG |
| 50 | Y_Junction_CoreStaple | GGAAACCAGGCGTTGGGAAGCTGTTCACACAACATACGAG |
| 51 | Y_Junction_CoreStaple | ATATTATAGTTGCGCCGACACAACTT |
| 52 | Y_Junction_CoreStaple | GCTTTGAGGACCCTGCTCCATGTTAACGAGGCTTAAT |
| 53 | Y_Junction_CoreStaple | CAAACCTCATATATTTTAAATTCTAGGGCGCGATGGCTTAG |
| 54 | Y_Junction_CoreStaple | CCAGCAGCGGCGGTCAGTATTAACCGAACGAACCA |
| 55 | Y_Junction_CoreStaple | GGCCCGGATTGACCGTAAGTATAAGCAAATATTGAACGGCGG |
| 56 | Y_Junction_CoreStaple | AAGAAACCATATTATTTATCC |
| 57 | Y_Junction_CoreStaple | AATAAAAGTATAGGAGTGTACTGGTCACAAACGCGCCTGATA |
| 58 | Y_Junction_CoreStaple | TTCTGACCGACCAGTAATAAAAGGGACATAGAGATAGAACCC |
| 59 | Y_Junction_CoreStaple | CGAATATAACGCGAGTAACAATATTACCGCCATTACTAGGGC |
| 60 | Y_Junction_CoreStaple | AGCTTAGATTACTTGCTTACAATTTCATTTGATGAAACAAAC |
| 61 | Y_Junction_CoreStaple | CCAGCAGAGAGGTGAAAATGAACATCACCTTGCTGAGTCAGT<br>TG |
| 62 | Y_Junction_CoreStaple | TATTACCACCAAATTTTAAAAGTTTTCGACAACTCGTATCAT |
| 63 | Y_Junction_CoreStaple | ACGAAAGAGGCCCCAGCGATTATACGCTGGCTAACGTAA |
| 64 | Y_Junction_CoreStaple | CAGCCCTTTTCATAGTTAGCGTGGAATTGCGTTTTAATAATAA<br>TTTTTTTCCACAGA |
| 65 | Y_Junction_CoreStaple | TTAAACAGCTTGGAGTGACTTTCCAGACGTTAGCCACCACCC |
| 66 | Y_Junction_CoreStaple | ACCCGTCGGCAGCTTTCCGGCTTTACCGCTTC |
| 67 | Y_Junction_CoreStaple | AGGTTGGGTGGTAACAAAATATATAACTCGTGTGATAA |
| 68 | Y_Junction_CoreStaple | TCATACAACGCTCCTTATCATTCCATACCGCGCCCAATAACG |
| 69 | Y_Junction_CoreStaple | GGCGAGGATTATAATTTTCCTTCCATACCGCTTAGAATCCTT |
| 70 | Y_Junction_CoreStaple | GACGGTCAATCTCCGCGATAAAGACCAGCATCGGAACGAAAA |
| 71 | Y_Junction_CoreStaple | TCAACAGTTTCAGCGATACCGCGGTCGCAAA |
| 72 | Y_Junction_CoreStaple | AATAACCTTCCAATAAATCATATCAACGCAGTCAAATCA |
| 73 | Y_Junction_CoreStaple | CACTACCGCCTGGCTTTTCCTGAGAGAGTTGTGTTCCAGTTTGG<br>AACT<br>TTAAGAGTC |
| 74 | Y_Junction_CoreStaple | CGGGCTCACTGCCCGCTGAAACCTGTCGTGCGAAAATCCCCTT<br>ATAAA<br>TCAAGTAGC |

| | | |
|---|---|---|
| 75 | Y_Junction_CoreStaple | CGGTACGCCATTAAAGGGATTTTAGCCAACTCTGGACAGGAA |
| 76 | Y_Junction_CoreStaple | TTGAAGATTAGTGAGGAAGGTTATCATATCTGACCTCAATGAGAG |
| 77 | Y_Junction_CoreStaple | TTGCAATTTCAGGAAAAACGCTGGCGTTAGTACCGAAGTAATTCTG |
| 78 | Y_Junction_CoreStaple | ATCCATACCTACATTTTGTACCGACATATGTACCGGCTT |
| 79 | Y_Junction_CoreStaple | GTAGAAAATAATAGCCGAACAAAGGAGATAATAACGTCCCT |
| 80 | Y_Junction_CoreStaple | TCAGAGAACCGCCACCCTCACAACATTTATGAGCCACCATTG |
| 81 | Y_Junction_CoreStaple | GAATCTTACCAGCAAGCACTGTCTTCAACATGTAATTTAAAA |
| 82 | Y_Junction_CoreStaple | TGAGGAAGTTTTTCCATTAAACGGGTAAAATATTGTATC |
| 83 | Y_Junction_CoreStaple | GAGGCGATAAAAACCAAACTAATGCAGATACAAAGATTCCCA |
| 84 | Y_Junction_CoreStaple | GTTAACCTACCATATCACACGTAAAACAGAAGTCAGATAGAAACAA<br>TAACGGAAATCG |
| 85 | Y_Junction_CoreStaple | AAAGACGAGAAACACCAGAACTTTTGAGTAGTA |
| 86 | Y_Junction_CoreStaple | TTATTAGAAACGCATTTTCGAGCCAGTAATAATTTGAGAA |
| 87 | Y_Junction_CoreStaple | ATATAACCAAAATAAATAGTTGATTCGCATAAAGCAGACTTGTTTTAAAT |
| 88 | Y_Junction_CoreStaple | CCGGAAGCTTTTATAAAGTGTAAAGCCTCCCCGGGTACC |
| 89 | Y_Junction_CoreStaple | TTGAGGGAGGGCAAATATCGCGTTTTAATTCGAGCTT |
| 90 | Y_Junction_CoreStaple | ACAGCTGATTGTTTTCACACAATTCTCCTGTGTG |
| 91 | Y_Junction_CoreStaple | TCGAAACTAAAAACGATCTAAAGTTGATAGCAAGCCCAACGG |
| 92 | Y_Junction_CoreStaple | TCTGACTACAAACGGATAATGGAAAATTGCGTAAAAAATCCAAGATT |
| 93 | Y_Junction_CoreStaple | CTGTAGCCAGACCAATAGGAACGCCTTATGCTTCAACTGCA |
| 94 | Y_Junction_CoreStaple | ATAACATGGAATGAGAAGTGTTTTTATATTTTATCAGTGAG |
| 95 | Y_Junction_CoreStaple | GCCTTGATATTAATAAGTTATTCTGAAACATGGGTGTATCAC |
| 96 | Y_Junction_CoreStaple | AATGAATCGGCCGCTGGTGCCCGAGATAGGGTGGCGCT |
| 97 | Y_Junction_CoreStaple | AGGACTGACTATTCAGAAAACGAGAGTCCAATACTGCGGCCA |
| 98 | Y_Junction_CoreStaple | GGCTATTACGTCCTCCAAGTCTTTAATGCGCGATTTTTGAAT |
| 99 | Y_Junction_CoreStaple | CGGTTGTACCACCTCAGACCAATTCTTTCATTCC |
| 100 | Y_Junction_CoreStaple | AGCACCATTACCGAGCCAGATATTGACGGAAATAAG |
| 101 | Y_Junction_CoreStaple | CAAATGTAGCATCACGTTGAAAATCAGGCTCCGGGTAGCGAG |
| 102 | Y_Junction_CoreStaple | CAAAGCTTTATTACAGGTAGATAACGCCCAGACGAGGGTA |
| 103 | Y_Junction_CoreStaple | ACCAAGAAAACACATTTACTGTAAATCGTCGCACATAGCTTA |
| 104 | Y_Junction_CoreStaple | ACATGAGAGGGTAGCAACTAAGAGCTATTTTTGTTAATATGC |
| 105 | Y_Junction_CoreStaple | ACCTAATGCTGATGCAAATTAATGGTTTGAAAACGCTCAAT |
| 106 | Y_Junction_CoreStaple | CCTGAGGACAGATGAACGCGGAGATCGTAATGGGCCGCTTT |
| 107 | Y_Junction_CoreStaple | AGGGCGCTATGGTTGCACGCAACCTTGCTGGTAATAGTA |
| 108 | Y_Junction_CoreStaple | AGAGCAATCAATAATCGGAATCAGATATAGAAATTTTATAAA |
| 109 | Y_Junction_CoreStaple | TTCATTAAAGAACGTGGAAAAGGGCGAAAAACGAGGCCGAGA |
| 110 | Y_Junction_CoreStaple | CCGAAAGACTTAAGGTAACAAAATCACACTGGAAAGACAGT |
| 111 | Y_Junction_CoreStaple | GAAGGAAACCGTTTTAGGAAACGCAATAATAACGGAACA |
| 112 | Y_Junction_CoreStaple | CAGGCGCATAGCAAGCGCCCAACCTTGAGGCTTGCAGGGTTC |

| | | |
|---|---|---|
| 113 | Y_Junction_CoreStaple | TGCGGGATCGTCACTTTTCCTCAGCAGCGAAAGATTTTTCA |
| 114 | Y_Junction_CoreStaple | TTCAGGTTTAACATAAAGAGGGTTAGTGGATTATACT |
| 115 | Y_Junction_CoreStaple | CCTAATTTTTTACGAGCATGCCGGTATTTTTTCTAAGAA |
| 116 | Y_Junction_CoreStaple | ATCCCGTTATTGAAGGAGCGCCGGACTTAGGAATTATCATCA |
| 117 | Y_Junction_CoreStaple | GCCACTGCTTTCCTCGTTAGAATTTTCAGAGCGGGAGCTAA |
| 118 | Y_Junction_CoreStaple | GCAAATAAGGAATAGCCGTCAATTCGCAATAATAGATAATATAA |
| 119 | Y_Junction_CoreStaple | AATCAGCTCAGCTCATTATACCAGTCAGGATTTCGTTGGGAAG |
| 120 | Y_Junction_CoreStaple | GCTGAGACTCCTTTTTCAAGAGAAGGATTGATAAGTGC |
| 121 | Y_Junction_CoreStaple | GTTAGCAGACAAAAGTTTTGGCGACATTCAACAAGACTCCTTAT<br>TACGTTTCAGTAT |
| 122 | Y_Junction_CoreStaple | CATCCATTTGGGAATTAATTAGCAAGGCCGGCCGTAATTCAG |
| 123 | Y_Junction_CoreStaple | AAGCCTGTTTATCCAGAAAAGAGTCTGTCCATCTTTGAC |
| 124 | Y_Junction_CoreStaple | AGGAGCCTTTAATTGTATCGGTTTTTTATCAGCTTGCTT |
| 125 | Y_Junction_CoreStaple | AATTGGGCTAACCGAACTGACTTTCAACTTTG |
| 126 | Y_Junction_CoreStaple | AATCGACCTTCATCAAGATTTGACCAAAAGAAATAACCGAT |
| 127 | Y_Junction_CoreStaple | TCATCTCAGGAAACCTATTTTAACGGGGTCAGCAGACGACCC |
| 128 | Y_Junction_CoreStaple | CGTTATACAAACTATCGGATTAACCGTTGTAGACAGGG |
| 129 | Y_Junction_CoreStaple | TCTTTTCAGTTTTGGGAGTAATCAAAATCACCGGGTTTGCCA |
| 130 | Y_Junction_CoreStaple | AAAAATCTACGAGAGATCTACAAAGGCTATTTTCAGGTCATTG |
| 131 | Y_Junction_CoreStaple | GACCCTGTATTTATACTTTTGCGGATATTC |
| 132 | Y_Junction_CoreStaple | CCCTATAACATTAAATCAAGATTAGTTAGCGAAACAAGAAGC |
| 133 | Y_Junction_CoreStaple | ACATCATATGGTTTACCAACTCCAAAGCTCAAATGGTC |
| 134 | Y_Junction_CoreStaple | TCCAGACGACGTTTACAATAAACAAAGAATGGATTTTAAGCG |
| 135 | Y_Junction_CoreStaple | ATCGCCTGATTTTTAAATTGTGTCGAAAATAAGGGTGAGA |
| 136 | Y_Junction_CoreStaple | CGTATTTCAGGTTGTCGTGAATAGAAAGGAACGGTGAATAGT |
| 137 | Y_Junction_CoreStaple | TGAGGGTCACGTTGGTGTAGCCCCAAAAACAGGCATGTCATT |
| 138 | Y_Junction_CoreStaple | CGTCGAGATTTGGGTTGATATAAGTATAGCCTAGGAAC |
| 139 | Y_Junction_CoreStaple | AACCGTTCTAGTTTTCTGATAAATTAATGCCGTAT |
| 140 | Y_Junction_CoreStaple | GTCAGAGGGTTTTAATTGAGCGCTAATATCAGATTACCA |
| 141 | Y_Junction_CoreStaple | CGCCAGTTGACAACATTCAGGCTGCGCAACTAAAGCGCCTTC |
| 142 | Y_Junction_CoreStaple | ATCAAGTTACAATTCGCCTGATTGCTTACGCCCTACATGAAT |
| 143 | Y_Junction_PolyT_Overhang | TTTCCCGCCGCGCTTAATGGCGCGTAGCAAAATCTGTTTGATG<br>GTGGTTTTT |
| 144 | Y_Junction_PolyT_Overhang | TTTATAAACAGCAATGAAATATTT |
| 145 | Y_Junction_PolyT_Overhang | TTTTTTACAAACAATGAGTAACTGCGGAACAAAGAACCTGATT<br>GATT |
| 146 | Y_Junction_PolyT_Overhang | CAAGTACCGCACTCATCGAGATTT |
| 147 | Y_Junction_PolyT_Overhang | TTTATTACCTGAGCAAAAGCGAATTACATCGGGGAATATACAG<br>TAAC<br>AGTTT |
| 148 | Y_Junction_PolyT_Overhang | TTACATCAGTTGAGATTTAGGAATTT |
| 149 | Y_Junction_PolyT_Overhang | TTTTACCACATTCAAATAGCGACAAAAGAAGTTTTGAATCGTC<br>TAAA |
| 150 | Y_Junction_PolyT_Overhang | CTTCTTTGATTAGTAATATTT |

| | | |
|---|---|---|
| 151 | Y_Junction_PolyT_Overhang | ACTGTAGCGCGTTTTCATCTTT |
| 152 | Y_Junction_PolyT_Overhang | TTTCGGAATAAGTTTGATATTTGCGGAGCTGAAAATTT |
| 153 | Y_Junction_PolyT_Overhang | TTTTTTTGCTAAAATGACACCCACGCTACACTAAATTT |
| 154 | Y_Junction_PolyT_Overhang | TTTGATAATCAGAAAAGATGGGCCGTGCATCTGCCATGGGTA AACGA |
| 155 | Y_Junction_PolyT_Overhang | TTTGAATTTATCAAAATCATAGGTCTGCAAGACAAAG |
| 156 | Y_Junction_PolyT_Overhang | TTTCTCAACAGTAGGGCTTAATTGAGGTATTAAAC |
| 157 | Y_Junction_PolyT_Overhang | TTTCAGTCACACTGAAAGCGTTTT |
| 158 | Y_Junction_PolyT_Overhang | CGCAGAGGAAGATGAATTACCTTTTTTAATGGAAACAGTTT |
| 159 | Y_Junction_PolyT_Overhang | TTTTCATCAATATAATCCTATCAGATGATGGCAATTTT |
| 160 | Y_Junction_PolyT_Overhang | CCCTCAGCCGCCACCAGAACCACCACCAGAGCCGCTTT |
| 161 | Y_Junction_PolyT_Overhang | TTTGTCACGACGTTGTAAACGCCAGGGTTTTCCCATTT |
| 162 | Y_Junction_PolyT_Overhang | TTTCCTCCCTCAGAGCCGCCA |
| 163 | Y_Junction_PolyT_Overhang | TTTACACTCATCGTAATCTCGGATATTCA |
| 164 | Y_Junction_PolyT_Overhang | AACGCGAGAAAACTTTTTCTTT |
| 165 | Y_Junction_PolyT_Overhang | TTTATTAAAAATACACCGCGCCACGCATATCAAACTTT |
| 166 | Y_Junction_PolyT_Overhang | TTTTAAACAGTTGAACCGCCATTT |
| 167 | Y_Junction_PolyT_Overhang | TTTGGTGGCATCAATGCAAATGTGTAGGTAAAGAATCATATGT ACC CCGGTTTTT |
| 168 | Y_Junction_PolyT_Overhang | TTTCCTCAATCATAAAATACACTAACAACTAATGGATTTAGAAGT ATTAGACTTT |
| 169 | Y_Junction_PolyT_Overhang | TTTACCATCGATAGCAGCAAAACGTCACTTGAGTAAAGGTGAA TTATCATTT |
| 170 | Y_Junction_PolyT_Overhang | TTTTCCCCCTCAAATGCTTATAAATATTCATTGAATTT |
| 171 | Y_Junction_PolyT_Overhang | TTTGCAATAGCTATCTTACCGAAGCCATAAAAGAAAC |
| 172 | Y_Junction_PolyT_Overhang | TTTCCCTCAGAACCGCCACCCTCAGAGTAAATGAATT |
| 173 | Y_Junction_PolyT_Overhang | TTTAAGAATACGTGGCACAGACAATAACTGATAGCCC |
| 174 | Honeycomb_Red24to21_Both | TCTCACGGTTTTTCATTTATTATTTTTTGTCCAACC |
| 175 | Honeycomb_Cyan15to15_3p | AGAGTCAATAGTTTTTTTTTGGTTGGAC |
| 176 | Honeycomb_Cyan28to15_5p | CCGTGAGATTACATAAATCAATATATGTGAGTGAATAACAGAC GCTGAGA |
| 177 | Honeycomb_Black27to27_3p | TGGCAGATTCACTTTTTTTTCCACACTG |
| 178 | Honeycomb_Black22to27_5p | ACCTCCTGTAAATATATTTTAGTTAATTTCATCTTCTGAATTATT TACAT |
| 179 | Honeycomb_Yellow12to29_Both | CAGGAGGTTTTTTAGGAGTCTTTTTTTTCAGTGTGG |
| 180 | Honeycomb_Pink19to19_3p | TAAAACATCGCCTTTTTTTTCCGTACCT |
| 181 | Honeycomb_Brown10to23_3p | TTTACAGTCTGCATTTTTAGGTACGG |
| 182 | Honeycomb_Magenta20to9_5p | CGATCACCTTACCTTTTATTCATTTCATTT |
| 183 | Honeycomb_Green16to5_5p | GGTGATCGTTTTTATTTGAAATTTTT |
| 184 | Honeycomb_Red54to51_Both | GCGGTAACTTTTTTTTTGGAGGCTTTTTGGAACTCG |
| 185 | Honeycomb_Cyan75to75_3p | ATAAAGCCAACGTTTTTTTTCGAGTTCC |
| 186 | Honeycomb_Cyan88to75_5p | GTTACCGCTACATCACTTGCCTGAGTAGAAGAACTCAAATTCTT ACCAGT |

| | | |
|---|---|---|
| 187 | Honeycomb_Black57to57_3p | ACAGTGCCCGTATTTTTTTTTGGAAGCC |
| 188 | Honeycomb_Black52to57_5p | CGTCATCCTCGCCAGCATTGACAGGAGGTTGAGGCAGGTTGCCTTGAGTA |
| 189 | Honeycomb_Yellow72to89_Both | GGATGACGTTTTTGAGTATGCCTTTTTTGGCTTCCA |
| 190 | Honeycomb_Pink49to49_3p | TTCTGTATGGGATTTTTTTTCTGCGAAG |
| 191 | Honeycomb_Brown70to83_3p | TTTTCATTAGAGGTTTTTCTTCGCAG |
| 192 | Honeycomb_Magenta50to39_5p | ACCATCGGTCCGTCACCGACCAATGAATTT |
| 193 | Honeycomb_Green76to65_5p | CCGATGGTTTTTTGTCGGTTCCATTT |
| 194 | Honeycomb_Red84to81_Both | TGTGCCAGTTTTTCGTAACGCATTTTTTCGCTACGT |
| 195 | Honeycomb_Cyan58to45_5p | CTGGCACATGGCATTTTCGGTCATAGCCCCCTTATTAGCAACCAGAGCCA |
| 196 | Honeycomb_Cyan45to45_3p | CCACCGGAACCGTTTTTTTTACGTAGCG |
| 197 | Honeycomb_Black87to87_3p | GCCAGTTACAAATTTTTTTTCCATGCTG |
| 198 | Honeycomb_Black82to87_5p | CCTTGACCTACAAGCAAGCCGTTTTTATTTTCATCGTAGGAGCCTAATTT |
| 199 | Honeycomb_Yellow42to59_Both | GGTCAAGGTTTTTAGAACTGACATTTTTCAGCATGG |
| 200 | Honeycomb_Pink79to79_3p | GCAAAGACACCATTTTTTTTCTCAGTCG |
| 201 | Honeycomb_Brown40to53_3p | TTTCATCGACAACTTTTTCGACTGAG |
| 202 | Honeycomb_Magenta80to69_5p | AGCTCTGCTCCGAAATCGACCACCACATTT |
| 203 | Honeycomb_Green46to35_5p | GCAGAGCTTTTTTGGATTAAAGCTTT |

**Table S3. 18 HB tetrapod staple oligonucleotide sequences**

| Count | Name | Sequence |
|---|---|---|
| 1 | Tetrapod_CoreStaple | CTAGTCTTTGATAAAAGAGTCTGTCATCAGAGCGGGAGCGCTTTG |
| 2 | Tetrapod_CoreStaple | AATCAACTCATCGAGAACATTTACGAGCATGTAACGCGCCTAC |
| 3 | Tetrapod_CoreStaple | CTTTAATGCGGCCCTAAAACATCGACTAATAGATTAGAGAT |
| 4 | Tetrapod_CoreStaple | GGAAGGCAAAAGCAGCAAGCGGTCCACATACGAGCCGGACGT |
| 5 | Tetrapod_CoreStaple | ACATGTTGTACAGGCTATAATATCCAGAACAATATTTTTACCG |
| 6 | Tetrapod_CoreStaple | ACATTATTACAGGTAAGGCATAGTAAGAGAGTAGCATTA |
| 7 | Tetrapod_CoreStaple | AAAACCAAAATAGCGAGAACAGTTCAGCATTTATTTTAAAAC |
| 8 | Tetrapod_CoreStaple | CAGAATGGATGATACAGGAGTGCCACCCTACCGTAAGAA |
| 9 | Tetrapod_CoreStaple | CAACAGTTGAAAATTCATAGTAACACCAGAGCACGGGAGAA |
| 10 | Tetrapod_CoreStaple | GGAATTGCCACTGAGTTTCGTCCACCCTCAGAACCGTAC |
| 11 | Tetrapod_CoreStaple | GAGATCTTTCCAAATGAAGCCCACGCATAACCTAAAGGCCGC |
| 12 | Tetrapod_CoreStaple | CCTGAGAAGAGACAGGAACGGTACGTGATACGAACCCAGAAT |
| 13 | Tetrapod_CoreStaple | ATATCGCGAAAATCAGGTCTTTTTCATTGGCCAGAGAGA |
| 14 | Tetrapod_CoreStaple | AAAATATCATACAACGCCAACATGTAAGAGAATATAAAGTGT |
| 15 | Tetrapod_CoreStaple | CAAAGCGGAACAAAGAAAGTATTAAAAACCGTGTGGACTGCGTATT |
| 16 | Tetrapod_CoreStaple | TTATCAACAATATCCTAAAGCAAGCCGCCCAATAAATCAAG |
| 17 | Tetrapod_CoreStaple | AATGGTTTGAATCAAATACTGATGCTTTTAACAGTCAATAGT |
| 18 | Tetrapod_CoreStaple | TAATTAGCGACTCAAAATCACCGGACACAAACAAATAAAAAT |
| 19 | Tetrapod_CoreStaple | GAAACCATACCGGAAGCAAACTACCTGTTTAGCTATGGG |
| 20 | Tetrapod_CoreStaple | ACGCTAATTTTCCCTTAGAATTTTAAATTAATCTTTTAGTTTG |
| 21 | Tetrapod_CoreStaple | TTGCCTGATAAAGCTAAATCGCCAATAAAATTACGGAA |
| 22 | Tetrapod_CoreStaple | CAAAGCGCAGACGGTCAATCATTTTTAAGGGAACCGACAGTA |
| 23 | Tetrapod_CoreStaple | TCAGTGATATTACCAGAGAGCACCGTAATCAGTCGAGCTTCA |
| 24 | Tetrapod_CoreStaple | AGATGAACCGGATATTCAAACTTTGAAAGAGGGCCAGTTT |
| 25 | Tetrapod_CoreStaple | AATGTGCTGGCATAGGAGCGAGTAAAGGTCACGTTGGTGTCG |
| 26 | Tetrapod_CoreStaple | GCGCTCTTTTACTGCCCGCTTGTAAAACGACGTTTTGCCAGTGCCA |
| 27 | Tetrapod_CoreStaple | AGATTTTCAGGGATTATACTTCTGACCCTCAAGCCACGCCTG |
| 28 | Tetrapod_CoreStaple | AACCCCGCCTGCAACAGTTCAATATCTGGTCATTGTTTGTTT |
| 29 | Tetrapod_CoreStaple | ATGCGATTTTAGGGGTAATAGTAAAAATCGTCATAAATAACC |
| 30 | Tetrapod_CoreStaple | CCCGGGCAGGTGCCTCCCTCAGAGCCGCTTTCACCCTCAGA |
| 31 | Tetrapod_CoreStaple | CCATTTTAATTCCATTAGCAAGGCTTTCGGAAACGTCACCAAT |
| 32 | Tetrapod_CoreStaple | ACCAGAAATCCGCGACCTTGCCGGAAAACACTAGAATCCAGGC |
| 33 | Tetrapod_CoreStaple | AAAGCGCCATCTTCTGGGCTCCATTTGCCCTGTTGGGAAGA |
| 34 | Tetrapod_CoreStaple | TGATATTCTAAAAATTTTTAGAATAACAGTTAGAGCATT |
| 35 | Tetrapod_CoreStaple | CCGTAAAGCACATTTGAGGATTTAGAAGTATTATTTTGACTT |
| 36 | Tetrapod_CoreStaple | CAATAAAGCCGCAAAGACAGATACTTAGGAACATCAAGAGT |
| 37 | Tetrapod_CoreStaple | GAGAATGACCAGCTTTAAGGCTTTTTCAGGACGACGAGAAAC |
| 38 | Tetrapod_CoreStaple | CATTCCATACCCTCAATCACCACGCTCAAACAGAGATAG |
| 39 | Tetrapod_CoreStaple | CCAGCCGAGAGGGTAGCTATTTTTTTTTGAGAGATCT |

| | | |
|---|---|---|
| 40 | Tetrapod_CoreStaple | CTTTCAACAGTTTCCACAGACAGCCCACCGTACTCAGGAGTG |
| 41 | Tetrapod_CoreStaple | AGCATTTTTGCGGATGGCTTGATTCCCAATTCGCAAGGAAAC |
| 42 | Tetrapod_CoreStaple | CTATCATTTTTAACCCTCGTTTACCAGACGACGGAGCTGAAAAG |
| 43 | Tetrapod_CoreStaple | AGTAACATTACGTTATTAATCCTTCGATAGCTTAGATTAAG |
| 44 | Tetrapod_CoreStaple | ACAACAAAAACATTATGACTTTTCCTGTAATACTTTTGCTGA |
| 45 | Tetrapod_CoreStaple | GGATAACCATCTTTTCTGTATGGGATTTTTTTGCTAAACAA |
| 46 | Tetrapod_CoreStaple | GCCATCGTCTGAAATGGACCGGAGACAGTCAATATATTTTAA |
| 47 | Tetrapod_CoreStaple | AATCAACGGTGTACAGACGCGCATCGTAACCGGAGGATCGGT |
| 48 | Tetrapod_CoreStaple | GGCAATTAACACTTCTGATACCTACATTTTGATCAATA |
| 49 | Tetrapod_CoreStaple | GGAATTATTTTTCATCATATTCCTGATGGGAGA |
| 50 | Tetrapod_CoreStaple | AACAATAACTTTTGGATTCGCCTGATTGAAGGAGC |
| 51 | Tetrapod_CoreStaple | TAAGTTGGGTAGGGGATGAGGGAGTGATATATGCAGAGGCAT |
| 52 | Tetrapod_CoreStaple | TTAGGCCTGGAGGCTTGCTGCTGCAAGTTCTGTCATCGCGAT |
| 53 | Tetrapod_CoreStaple | CCCCAAAGAATCCGCCTGGCCCTGAAGTGTAAAGCCTGGCCC |
| 54 | Tetrapod_CoreStaple | TATTACGCAGTACGTAGATTACCAGAAAAGGGAATTAGAGCC |
| 55 | Tetrapod_CoreStaple | GTGGCATCAATTTTTTCTACTAATAGTCAACA |
| 56 | Tetrapod_CoreStaple | GGGCGCCGGGAAACCTGTCGTGGTTTTCCCAGTCATATTACGGCG |
| 57 | Tetrapod_CoreStaple | AGATTCCCGACAAGGCTTATCCGGTGGGTATTAAACCAAATCAATAAT CGGCTAAC |
| 58 | Tetrapod_CoreStaple | CCTCCAATGAATGAACACCCTGAACAAATTTGTCAGAGGGTAATT |
| 59 | Tetrapod_CoreStaple | GCCTTGCCTTGAGTAACAGGTTTAGTGTAGCATTCAGCGAAC |
| 60 | Tetrapod_CoreStaple | ATTAGAGAGGCGAATTATTCATTTCAGAAGCCTTAGCAAGCA |
| 61 | Tetrapod_CoreStaple | AATACTAAATCTTGAGTGGCAACAGCTGATTGAATGAGTGAG |
| 62 | Tetrapod_CoreStaple | TAGCCGAAAGAACTGGCATGATAGGTGGCTCAATAGTTG |
| 63 | Tetrapod_CoreStaple | CGGGGAGAGGCATTAATGAATCGGCATATAGGGTTCAACGCG |
| 64 | Tetrapod_CoreStaple | CGGAGCTTTCCGGCACCGTCGCCATCTACAGATTTCTTAGAG |
| 65 | Tetrapod_CoreStaple | AAGCGAACCAGCGATAGCCCACCACCGGAACCCAGACGATTG |
| 66 | Tetrapod_CoreStaple | GAATCTGTAAATCGTCGCGTACATAAATCAATTAACAATAGA |
| 67 | Tetrapod_CoreStaple | AGTATAGCCCGGTTTTAATAGGTGTATCTCAT |
| 68 | Tetrapod_CoreStaple | ACCAGTACCGCGATATAGTTGCGGGAGGTTTTATTACCTGAG |
| 69 | Tetrapod_CoreStaple | ACGTCACATAGGAAAAAAGGGCGAAATCCTTTGCCCGAATCA |
| 70 | Tetrapod_CoreStaple | GTTGATATAAACAGTTAATGTTTTCCCCCTGCCTACGCCGCCAGCATTT TTTGACAGGAG |
| 71 | Tetrapod_CoreStaple | TGAATCTTTTTACCAACGCTAACGAGCGTCTTTGTACCT |
| 72 | Tetrapod_CoreStaple | ATGCTGTCTGGTAATGCTTCACCGTAGGGAAGGTAAATATTA |
| 73 | Tetrapod_CoreStaple | CGTAAAACAGGAAGATTGTATTTAAGCTCATTTAGGAACTTA |
| 74 | Tetrapod_CoreStaple | GCAAGAAAAAGAGAAGGATTAGGTTTTATTAGCG |
| 75 | Tetrapod_CoreStaple | AAATAATATCCCAGATAAGCAGTAATAATTTAGTCGGTCGCTT |
| 76 | Tetrapod_CoreStaple | GAGCGCTAATTAACCCACAAGAATCTCAGTACCAGGCGGTC |
| 77 | Tetrapod_CoreStaple | TCAGTTGAGATATAACGCCAAAAGGATCATACAGGCAAGTCA |
| 78 | Tetrapod_CoreStaple | GAGGCGAGGCCGATTAAAGTACAGGGCGCGTACTGGCAAGCGG |
| 79 | Tetrapod_CoreStaple | TTAACATAGCAATAGCTATCTTACCGGCATTAGCTAATT |
| 80 | Tetrapod_CoreStaple | TTCGACAACTCCCACCAGCTTTGAACAATTTTATCC |

| | | |
|---|---|---|
| 81 | Tetrapod_CoreStaple | GTGAAGGCTTTGAGGACTAAAGACTTTCAGCTTACTAAA |
| 82 | Tetrapod_CoreStaple | AGTCAGGGTGGTTTTTCTTTTTTTCACCAGTTAATTGCGTT |
| 83 | Tetrapod_CoreStaple | GCAAGCGAGAAAACTTTTATACCGAAACACCGTCTTACCATT |
| 84 | Tetrapod_CoreStaple | TTGATGTAGCGGTCACGCGCCGCGGATTTTTGTTTTTTGAGTAGA |
| 85 | Tetrapod_CoreStaple | TAATATTGTATCGGTTTATTTCATGAGGAAGTACGAAA |
| 86 | Tetrapod_CoreStaple | TCTAAAATATCTTTTTTTAGGAGCACTAACACCATTAAAA |
| 87 | Tetrapod_CoreStaple | TGCGCCGTTTACAATGACAACCGTCACCCTCAGCAGCTTTTGAAAG |
| 88 | Tetrapod_CoreStaple | TTTTGTCACAAAACATATAAAAGAAACCCAAACAAAGTTATA |
| 89 | Tetrapod_CoreStaple | CCTTTTGATAAGAGTAGATTTAGTTGGGAGAAAATGCCGATT |
| 90 | Tetrapod_CoreStaple | TACAAAATCAAGTTTTTTTTTTGGGGTCGAGGTG |
| 91 | Tetrapod_CoreStaple | AATCATGGTCAATGGGATCAACCCGATCAACAGCCATCAAAA |
| 92 | Tetrapod_CoreStaple | ACGTCAGGAAAAACGCTCCTAGCTGATAAATTGCCTTTATTT |
| 93 | Tetrapod_CoreStaple | CACCACCAATAGCCGTACCCGCCGCGCTTAATTGCGCGTAGC |
| 94 | Tetrapod_CoreStaple | CCAGGCAAAAGAAGTTTTAATCCCCCTCAAATTAAATCATTT |
| 95 | Tetrapod_CoreStaple | GCGCATAAAAATCTACGTTCAGTGAATAAGGCGTTACTTAGC |
| 96 | Tetrapod_CoreStaple | AAGTTTTAACGGAACCGCACCAGTAAGGAACAGCTTTCGAG |
| 97 | Tetrapod_CoreStaple | AGGGCACCGACTTGAGCCTTAATTGCTGAATAAAGTTT |
| 98 | Tetrapod_CoreStaple | GGAACCCATGTCAGAGCCACCACCCGCTTTTGAAAGCGCCTT |
| 99 | Tetrapod_CoreStaple | TTTTCCAGCTGGCGAAAGACGCCAGGCCAGCTGCGGTTTCCA |
| 100 | Tetrapod_CoreStaple | CAACTGCGAACGAGGTCAAAATCACCACCACCCTAAGACCGC |
| 101 | Tetrapod_CoreStaple | GGGTACCGAGCTAGATGGCAGGCGCTGACCTTTACCACATTA |
| 102 | Tetrapod_CoreStaple | CTGACGAAAGACTTTAGCGCGTTTGCCATCTTTAAAGC |
| 103 | Tetrapod_CoreStaple | TGGTTCCTCATTTCATAAAGAATCAAGTTTGCCTTCAA |
| 104 | Tetrapod_CoreStaple | AAAGGGACATTTGAGAGCCAGCAGCCCTCAAATATCAAAATA |
| 105 | Tetrapod_CoreStaple | CGTTATGGAAACCTGAAAGCGTAAGGTGAGGCGGTCAGTAAT |
| 106 | Tetrapod_CoreStaple | ACCGCCACCCACCACCACCAGAGCTTTCGGAACCTATTACT |
| 107 | Tetrapod_CoreStaple | GAGGGGCATGCCTCTCACATGAGACGGTTGTTCCAGTTT |
| 108 | Tetrapod_CoreStaple | GAGGCAAAAAAACACTTGTGTCGAACGAGGCTCATTATA |
| 109 | Tetrapod_CoreStaple | AGTTAGTTTTCGTAACGATCTAAAGTTTTGTCGGGGTTGATATA |
| 110 | Tetrapod_CoreStaple | TTTACATCTATCAGATGATGGCAGGAATTAACAGAGAAT |
| 111 | Tetrapod_CoreStaple | ATTAGTCAGACTGTAGCGATTAAGAGGAAGCCCTATTATAG |
| 112 | Tetrapod_CoreStaple | TGCCAGTTTCAGATGAATATACCAATATAATCCTGAGTT |
| 113 | Tetrapod_CoreStaple | AGCTTGACGACGAACTGACCTTACCCAAATCAATTTCGTAA |
| 114 | Tetrapod_CoreStaple | GGTTTAAACAGCACCGAGTTAGTAATAACATCGAACGGTAAT |
| 115 | Tetrapod_CoreStaple | CACAGAGCCGCAACATGAAAAGCAAATGTTGTATTAAGAGGC |
| 116 | Tetrapod_CoreStaple | ACAGCAAAGGGCGATCGTTTTGTGCGGGCCTCTTCGCCGACGTTTCC |
| 117 | Tetrapod_CoreStaple | CGACTATTTAATGCGTTATACAAATGAATCATAATTACTTTT |
| 118 | Tetrapod_CoreStaple | GGGTTTTGTGAGTTAAGCCCAATTTTTAATAAGA |
| 119 | Tetrapod_CoreStaple | GCAGCAATCCAAATAAGAAAAACAGAAATAAAAAGGGTTAG |
| 120 | Tetrapod_CoreStaple | GAGAATCGCCAAAAAGGTAAAGTAATCAGCTAATGCAGAGAA |
| 121 | Tetrapod_CoreStaple | AGCTTGATACCGAGGGTAGCAACGGTCAGGCTGCGCAACCCA |
| 122 | Tetrapod_CoreStaple | GAGCAGAGTCTCAAACTATCGGCCTTGCTGGTCAGGTCA |
| 123 | Tetrapod_CoreStaple | GCCAACGAGGCTGCTCATTAATAAAACGAACTAACGTTTGAACA |

| | | |
|---|---|---|
| 124 | Tetrapod_CoreStaple | TGAGAATAGAACAAACTACAACGCCTACCGCCACCCTCAGGG |
| 125 | Tetrapod_CoreStaple | AGAACTGGAGCAAACAAGAGAATCGATACTTGCCATAATCAGT |
| 126 | Tetrapod_CoreStaple | ATAATAAGACTAAAAGTAATAACATAAACAGCCATATTACGT |
| 127 | Tetrapod_CoreStaple | TAGTAGACGTTTTCGAGCTCCTGAACAGTGCCGATAAAGAA |
| 128 | Tetrapod_CoreStaple | ATACCGTTTTAACGAACCACCAGCAGAAGATAAGAGGAAGGTTA |
| 129 | Tetrapod_CoreStaple | AACGACAAAATAAAAACAGGGAAGCAAGCCCTTTTTAAGCCT |
| 130 | Tetrapod_CoreStaple | GCAAGGCACAGACAATATTTTTTTGAATGGCTATTAGT |
| 131 | Tetrapod_CoreStaple | TTTTATCGCGCTTGCTATTTTGCACCCAGCTATACCAAGTTA |
| 132 | Tetrapod_CoreStaple | TGGGCGACATTCAACCGAAAAATTCATATGGTAAATACATAC |
| 133 | Tetrapod_CoreStaple | GCCTCCCTTCAAGCCCGAGATAGGGGGAACCCTAAAGGGAAC |
| 134 | Tetrapod_CoreStaple | TCAATACCAACAGGTCAGGATTTTTTAGAGAGTACCTAGATACATTTT TTTCGCAAATGG |
| 135 | Tetrapod_CoreStaple | TGAGATTCTGACACCAGATCAGAGCCAGTAGCACCATTAGCT |
| 136 | Tetrapod_CoreStaple | ACTGTAGTAAATTGGGCTCATCGCCTGATAAATCATCTTTGA |
| 137 | Tetrapod_CoreStaple | ATAATTTTTGTTAAATCAATTGTAAACGTTAAGCCCCAAAAA |
| 138 | Tetrapod_CoreStaple | TCATCAGAGAATCAGGTAGGAATCATTACCGCGTTTTTATTT |
| 139 | Tetrapod_CoreStaple | GTTCCGAAATCAGCCGGCGAACGTGGGGCGCTAGGGCGCTAT |
| 140 | Tetrapod_CoreStaple | GGAACATTTAGAGTCCACTATTAAAGAACCTATCATCACCCAACAA |
| 141 | Tetrapod_CoreStaple | TCGGCCTTTTTCAGGAAGATCGCACTTGTTGGGTCGGAACGATAGT |
| 142 | Tetrapod_CoreStaple | CTAAGCAGGTCGACTCTATGCATCTACAGATGTTGACAATCA |
| 143 | Tetrapod_CoreStaple | CCAATTTTACAACTAATGATTAGCAAAGCAAATATAAATTAAG |
| 144 | Tetrapod_CoreStaple | CAAATTCATTTGAATTACGGAAACATATTAATTGAGAAGCTCCGG |
| 145 | Tetrapod_CoreStaple | CTTAGGTTACTATATGTAAATGTATTTTAGTTAATTACCTAAAAGA |
| 146 | Tetrapod_CoreStaple | ATGGGAAATTGTTTATCCCCTTTACAGAGAGAAGCAGA |
| 147 | Tetrapod_CoreStaple | AATTAGCATAAGAGAGTTTCCCTTATAAATCAGATTTAGAGC |
| 148 | Tetrapod_CoreStaple | GACCGTATAGCTGTCCACACAACGCTGGGTG |
| 149 | Tetrapod_CoreStaple | AGTACGGAATGCCTAGAAAGGTTATTTAATA |
| 150 | Tetrapod_CoreStaple | GGCACCACAGCGATTTTGTATTGAGATGCTT |
| 151 | Tetrapod_PolyT_Overhang | TTTAACGCAATAATAACGGAATACGCAAAGACACTTT |
| 152 | Tetrapod_PolyT_Overhang | TTTGCGTCCAATAGCAAAGCGGATTTT |
| 153 | Tetrapod_PolyT_Overhang | TTTAGAAAGCGAGTATAACGTGCTTTT |
| 154 | Tetrapod_PolyT_Overhang | TTTATAACCTTGCTTTTATCAACTGAGAGACTACCTAAATCCAATC |
| 155 | Tetrapod_PolyT_Overhang | TTTACAATAAACGTCTTTCCTTATTTT |
| 156 | Tetrapod_PolyT_Overhang | TCAGAACTGCGGATGTTTAGACTGGATATTTTTT |
| 157 | Tetrapod_PolyT_Overhang | ATGTTTCTGTCCAGACGACGTTT |
| 158 | Tetrapod_PolyT_Overhang | ACGAGCACAAGGAGCGCGAGAAAGGAAGGGATTT |
| 159 | Tetrapod_PolyT_Overhang | TTTTTCCTCGTTAGACATCACGGTTGTAGCAATACTCATGTCAATCAT ATGTACCTTT |
| 160 | Tetrapod_PolyT_Overhang | TTTATTCGCATTAAATTCGCGTTGTAGCCAGCTTTCTCGGATTCTCCGT GGGAACTTT |
| 161 | Tetrapod_PolyT_Overhang | TTTATAGCAAGCCCAGAAAATCAGGCTCCAAAAGGAAAACGGGTAA AATACGTAATTT |
| 162 | Tetrapod_PolyT_Overhang | TTTTTATTTGCACGTAACGATTCGTCAAAAATGAAAACCAGAAGGAAA CCGAGGATTT |

| | | |
|---|---|---|
| 163 | Tetrapod_PolyT_Overhang | TTTGCATCACCTACCATATCAAAATTT |
| 164 | Tetrapod_PolyT_Overhang | TTTTGCATCAAAAAGCGTTTTCTCGGTCATAGCCCCAGTCTCTGAATTTACCGTTTTT |
| 165 | Tetrapod_PolyT_Overhang | AACCTTGCTGAAAAATGAAAAATCTAAATTT |
| 166 | Tetrapod_PolyT_Overhang | TTTCACGGAATAAGTTTGACGGTTAAAGGTGAATTAGTAGCTCAACATGTTTTAATTT |
| 167 | Tetrapod_PolyT_Overhang | TTTCATTCCAAGAACATTCTAACGTTTTAGCGAACCGATGAAACAAACATCAAGATTT |
| 168 | Tetrapod_PolyT_Overhang | TTTTTATTCAAAATTTTTTT |
| 169 | Tetrapod_PolyT_Overhang | TTTTTTAGGTAATCATTTTT |
| 170 | Tetrapod_PolyT_Overhang | TTTTTCTTCCCTGGCTTTTT |
| 171 | Tetrapod_PolyT_Overhang | TTTTTAAAAATCCAATTTTT |
